# Assessment of a western blot signal for the Bcnt/Cfdp1, a tentative component of Srcap chromatin remodeling complex; trial to overcome off-target problems

**DOI:** 10.1101/739862

**Authors:** Shintaro Iwashita, Takehiro Suzuki, Yoshimitsu Kiriyama, Naoshi Dohmae, Yoshiharu Ohka, Si-Young Song, Kentaro Nakashima

## Abstract

The BCNT (Bucentaur) protein family is characterized by a conserved amino acid sequence at the C-terminus (BCNT-C domain) and plays an essential role in gene expression and chromosomal maintenance in fungi, fly, and chicken. The mammalian Bucentaur/Craniofacial developmental protein 1 (Bcnt/Cfdp1) is also a tentative component of the Srcap (SNF2-Related CBP Activator Protein) chromatin remodeling complex, but little is known about its properties, partly because there are few suitable antibodies to detect the endogenous protein. We used multiple anti-Bcnt/Cfdp1 antibodies against unrelated immunogens derived from BCNT-C domain and mouse-specific N-terminal peptide. To assign western blot signals and evaluate these antibodies, we utilized a stem cell line from mutant embryos of mouse *Bcnt/Cfdp1*, whose mRNA expression levels were reduced to 75% of the parental cells. In western blotting of these mutant and parental cell extracts with the anti-Bcnt/Cfdp1 antibodies, mouse Bcnt/Cfdp1 was detected as a doublet of approximately 45 kDa. LC-MS/MS analysis of the corresponding doublet for the Flag-tagged mouse Bcnt/Cfdp1 constitutively expressed in T-REx 293 cell (a HEK293 derivative) exhibited that the upper band was much more phosphorylated than the lower band and that there was preferential Ser phosphorylation in the WESF motif in the BCNT-C domain. Western blot with these validated antibodies indicated a preferential expression of Bcnt/Cfdp1 in the early stages of brain development in mouse and rat, which is consistent with the expression of Bcnt/Cfdp1 mRNA. This article describes the evaluation of anti-Bcnt/Cfdp1 antibodies, including a scheme to prepare a potential negative control for western blot, and discusses immune-cross reactions with off-target proteins, particularly immunoreaction probabilities.

## Introduction

The BCNT family members in yeast, *Drosophila*, and chicken have been shown to play essential functions in gene expression and chromosomal maintenance [1]. Mammalian Bcnt/Cfdp1 is also presumed to be a tentative component of the SRCAP (SNF2-Related CBP Activator Protein) chromatin remodeling complex (Human soluble protein complexes; http://human.med.utoronto.ca/php/search_complex.php?clusterid=595) based on the analysis of Swc5, a budding yeast ortholog of Bcnt/Cfdp1, in Swr1 (yeast Srcap) chromatin complex [2–4]. Although Swc5 is not integrated with the Swr1 complex, it participates in activation of the remodeler ATPase and in the ATP-dependent histone exchange reaction, which replaces nucleosomal H2A–H2B with H2A.Z–H2B dimers by recruiting the variant H2A.Z in transcription and DNA repair [5–7]. *Swc5* is not essential for survival, but its deletion mutant *swc*5Δ cells result in genetic instability, hypersensitivity to drugs, and transcriptional misregulation because Swr1 binds to chromatin but lacks histone replacement activity [8][The *Saccharomyces* Genome Database https://www.yeastgenome.org/].

The protein structure of the Bcnt family members generally consists of an acidic N-terminal region, a highly conserved C-terminal region with about 80 amino acids (BCNT-C domain), and a hydrophilic region between them (S1 Fig and [1]). Recently, the BCNT-C domain of Swc5 was found to be essential for the histone exchange reaction [9]. On the other hand, a point mutant of the *Drosophila* ortholog, *Yeti*, which entirely lacks the BCNT-C domain, shows substantial chromosomal abnormalities, resulting in lethality before pupation [10]. Thus, the metazoan *Bcnt/Cfdp1* is essential for survival in contrast to the yeast *swc5*. Furthermore, the chicken ortholog CENP-29 has been identified to be a kinetochore-associated protein [11]. Given a report that CENP-B protects centromere chromatin integrity by promoting histone deposition [12], these results imply that the Bcnt members may play a broader role in the maintenance of the structure and function of the chromosome.

A RNA sequence analysis, as shown by the dramatic influence caused by genetic mutations of *swc5* in fission yeast [13], has revealed complex mechanisms and dynamic processes such as embryonic development and stress adaptation. However, recent studies have shown that there exists discordance between mRNA and protein expression in such dynamic processes and argue that analysis at the transcriptional level is insufficient to predict protein levels [14]. While these processes are quite complex, involving both noncoding RNAs and antisense RNAs, the western blot analysis is the most straightforward way to examine changes in target molecules at the protein level. Because the Bcnt family members may function preferentially in these dynamic processes, analysis of their protein dynamics is essential. However, most of the currently available antibodies against Bcnt/Cfdp1 are challenging to assign the correct signal in western blot analysis.

We previously characterized human BCNT/CFDP1 (hBCNT/CFDP1) using a constitutively expressed His-tag molecule [15] and the custom-made antibody generated against an 18-mer peptide (EELAIHNRGKEGYIERKA) located in the BCNT-C domain (anti-BCNT-C Ab) [16]. Despite a calculated mass of 34.9 kDa (33.6 kDa plus His-tag) of His-hBCNT/CFDP1, the immunoreactive signal was detected around 50 kDa as a doublet band on SDS-polyacrylamide gel electrophoresis (SDS/PAGE), and we showed that the difference in its apparent molecular mobility is mainly due to the acidic stretch located in the N-terminal region and Ser^250^ phosphorylation in the BCNT-C domain [15]. However, we failed to identify the endogenous hBCNT/CFDP1 due to high background caused by anti-His Ab reactive proteins and to accurately assess the specificity of the anti-BCNT-C Ab to endogenous hBCNT/CFDP1. Furthermore, we recently found that the anti-BCNT-C Ab cross-reacts with a completely unrelated target, glutamine synthetase (NP_001035564.1; EC 6.3.1.2, which is also known as γ-glutamate: ammonia ligase)[17]. In this paper, we evaluate and validate the anti-Bcnt/Cfdp1 Abs and assign western blot signals using various target-related materials, including *Bcnt/Cfdp1* knockdown cells. We also present a scheme to prepare a potential negative control for western blot to detect Bcnt/Cfdp1. Then, we demonstrate high expression of Bcnt/Cfdp1 at an early developmental stage of the brains of mice and rats by using the above-evaluated Abs. We also discuss off-target proteins in terms of immune reaction probability based on the analyses to solve the tasks of the present study.

## Results

### Detection of Flag-tagged mBcnt as a doublet band

To characterize mammalian endogenous Bcnt/Cfdp1, we first expressed Flag-tagged mouse Bcnt/Cfdp1 (F-mBcnt) in T-REx 293 cells (a derivative of HEK 293) as a reference. We did this because although we previously expressed exogenous His-tagged hBCNT/CFDP1, the relatively high background by cross-reacting proteins with anti-His tag Ab made it difficult to assign the signals in western blot [15]. Therefore, we used Flag-tagging to expect the lower tag specific background, though having negatively charged sequence (DYKDDDDK).

The mBcnt/Cfdp1 is composed of 295 amino acids, which is four amino acids less than the human counterpart. The N-terminal region has low homology between mouse and human (75%) and can be used as species-specific immunogens, while the C-terminal 82 amino acid sequence of the BCNT-C domain is identical except for two amino acid residues (S1 Fig, [1]). We isolated T-REx cell colonies that constitutively expressed F-mBcnt using G418 selection. Both the number and size of the antibiotic-resistant colonies from F-mBcnt transfectants were significantly lower and smaller than those from F-multi-cloning site (F-MCS) transfectants as a control. After growing each colony in the presence of G418, the extracts were prepared and evaluated by western blot using either anti-Flag Ab or anti-BCNT-C Ab (Fig 1, S1 Fig). Whereas all signals with anti-Flag Ab showed doublet bands, the anti-BCNT-C Ab detected one and three bands in the extracts from the F-MCS-derived and F-mBcnt derived colonies, respectively (S1 Fig). Compared with the doublet pattern between transiently expressing cells and constitutively expressing cells, the upper band (Upper) in the transient expression was significantly stronger than that of the constitutive expression (Fig 1). These features are similar to those of His-tagged hBCNT/CFDP1, as previously reported [15].

**Fig 1.**
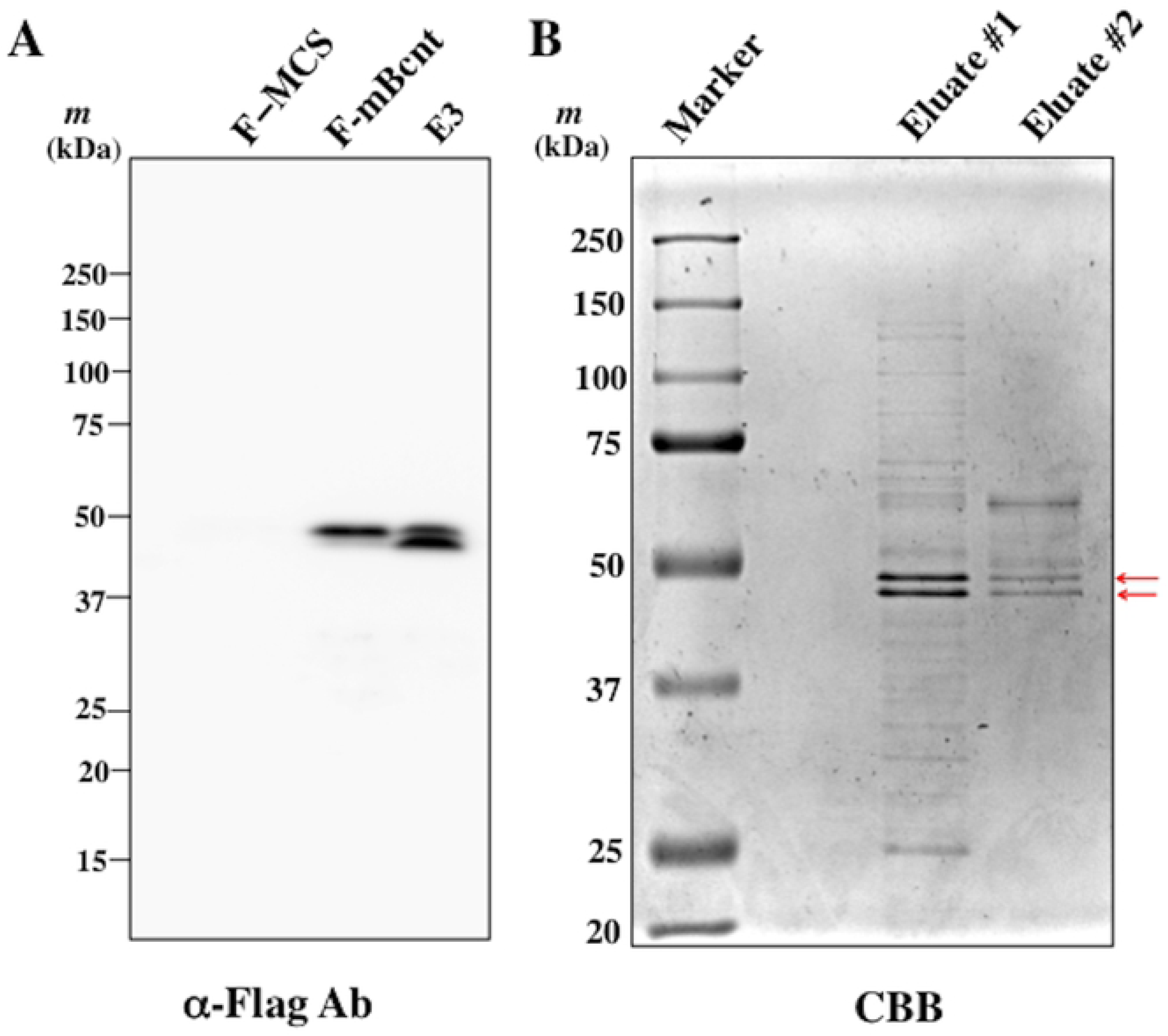
F-mBcnt expression as a doublet band and isolation of each band. (A) Flag-tagged mBcnt was detected as a doublet in both transient and constitutive expression. The F-mBcnt or F-multi-cloning site (MCS) in the vector was expressed in T-REx cells, and the extracts were prepared after culturing for 46 h as transiently expressed samples (Lanes: F-MCS and F-mBcnt). In addition, the extract of the E3 colony that constitutively expressed F-mBcnt was prepared (~5 × 10^4^ cells). These proteins were subjected to western blot analysis with anti-Flag Ab. The image in this figure is a replica of part of S1 Fig. (B) Isolation of the upper and lower bands from F-mBcnt doublet expressed in the E3 colony. The supernatant of the E3 extract isolated by centrifugation was applied to anti-Flag-tag agarose beads, and the adsorbed fraction was eluted, as shown in S2 Fig. After evaluation of the chromatogram, the more massive Eluate #1 and #2, which had been isolated by sequential elution with Flag peptide, were separated on SDA/PAGE and detected by Coomassie Brilliant Blue staining. The arrows indicate a doublet band.

### Phosphorylation with serine^246^ preference in the upper band

To reveal the molecular differences between the upper and lower bands, the bands were isolated from lysates of T-Rex-derived E3 cells that constitutively express F-mBcnt using anti-Flag Ab conjugated agarose beads (S2 Fig, Fig 1). Each band excised from the gel was digested with three different proteases—that is, *Achromobacter* protease I (API), AspN, and chymotrypsin—and each digest was subjected to LC-MS/MS analysis. In the analysis where each digested fragment covered 60-67% of the entire F-mBcnt/Cfdp1 sequence, the ratios of the upper to lower bands for each fragment amounts were listed (S1 Table). Furthermore, we focused on phosphorylation sites and presented them systematically (Fig. 2). The upper band is much more phosphorylated than the lower band, and, in particular, the serine 246th mBcnt/Cfdp1 that corresponds to the S250 of hBCNT/CFDP1 is apparently the preferential site of phosphorylation. Their characteristics are very similar to those of His-tagged hBCNT/CFDP1, as previously described [15].

**Fig 2.**
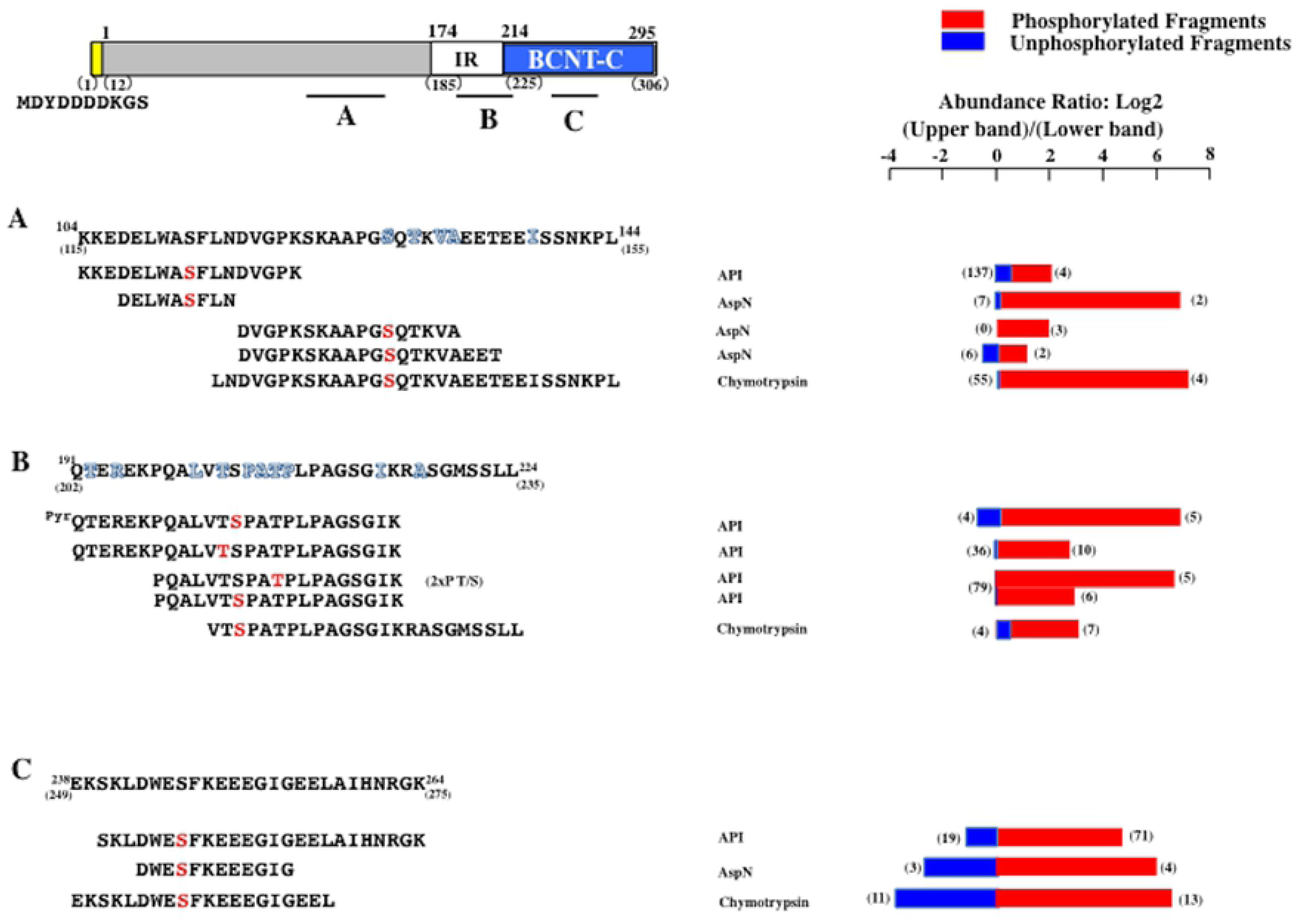
Differential phosphorylation between the upper and lower bands of F-mBcnt. Top panel: Molecular architecture of Flag-tagged mouse Bcnt/Cfdp1. The protein shown in a large outline comprises the acidic N-terminal region, Lys/Glu/Pro-rich 40 amino acids (named intramolecular repeat [IR], white box), and a highly conserved C-terminal region (BCNT-C domain, blue box) in addition to the Flag-tag at the N-terminus (yellow box). The numbers above or below the outline show the amino acid residues of the protein with (below) or without (above) the Flag tag, respectively. Three black bars A, B, and C indicate the regions of focused phosphorylation sites. Each upper and lower band of F-mBcnt from Fig. 1B were digested with three proteases (API, AspN, and Chymotrypsin) and subjected to LC-MS/MS analysis. All of the identified peptides are listed in Table S1, and their typical phosphorylated fragments and their unphosphorylated counterparts are represented. Each top amino acid sequence in A, B, and C represents each focused region, and the numbers at the N-terminus and the C-terminus correspond to the amino acid residue of mBcnt, respectively. **X**s are different amino acid residues from human BCNT/CFDP1. Red letters show the identified phosphorylated sites. Red bars indicate the ratios of the amounts of identified phosphorylated fragments in the upper band compared to those in the lower band, and blue bars show the corresponding ratios of their unphosphorylated fragments, respectively. The numbers of peptide-spectrum match (PSM) values are presented in parentheses.

### Evaluation of the anti-BCNT-C antibody

We had assumed that the ~50-kDa signal above the F-mBcnt doublet detected by anti-BCNT-C Ab corresponds to hBCNT/CFDP1 from previous results [15]. However, the band migration was judged to be too slow on SDS/PAGE, because F-mBcnt is expected to be a slow migration due to the acid Flag tag. Therefore, we reexamined the previous data of His-hBCNT/CFDP1 [15] in the western signal by introducing anti-Bcnt/Cfdp1 Abs. We chose the two commercial anti-BCNT/CFDP1Abs from the following criteria; the description of the immunogen is clear and the candidate signal is detected in a region significantly smaller than 50 kDa in western blot. They are 26636-1-AP9 (Proteintech, (https://www.ptglab.com/products/CFDP1-Antibody-26636-1-AP.htm) and A305-624A-M (Betyl, https://www.bethyl.com/product/A305-624A-M/CFDP1+Antibody). According to each catalog, the former Ab has been generated using the larger immunogen (172-299 hBCNT/CFDP1); it detects a single band with 45-50 kDa, and the specificity is validated by siRNA knockdown. The latter Ab recognizes the region of 249-299 hBCNT/CFDP1, has immunoprecipitation ability and detects ~48, ~37, and ~19 kDa signals. However, we found that A305-624A-M detected only a major signal around 48 kDa, whereas 26636-1-AP9 revealed other signals, including a signal of a ~50 kDa (as shown later). Based on the results, we compared western signal patterns between the anti-BCNT-C Ab and A305-624A-M using cell lysates of parent T-REx and its derivative G11 clone that constitutively expresses His-tagged hBCNT/CFDP1 [15]. Both Abs efficiently recognized exogenously expressed His-hBCNT/CFDP1 in the G11 extract but showed distinctly different patterns in the T-REx extract. It is noteworthy that the anti-BCNT-C Ab reacted to a band above the doublet detected with A305-624A-M (Fig 3, left panel). The difference between the two patterns was confirmed by reprobing with each of the replaced Abs (Fig 3, right panel). Whereas the anti-BCNT-C Ab revealed a few other bands, A305-624A-M detected a doublet but not the bands of ~37 and ~19 kDa demonstrated in the catalog. These results strongly suggest that the distinct ~ 50-kDa band detected with anti-BCNT-C Ab as shown in S1 Fig. is an off-target signal.

**Fig 3.**
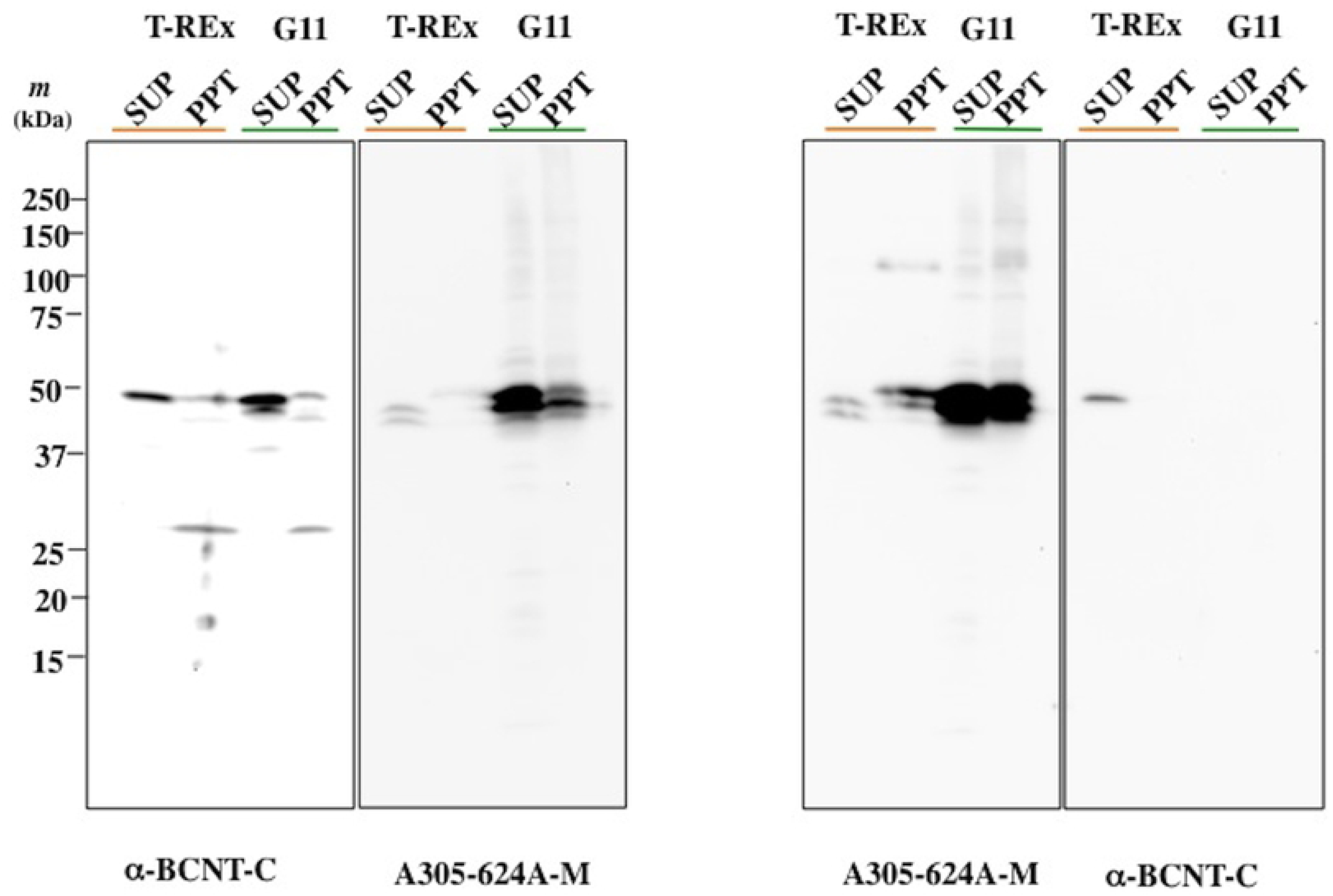
Comparative assessment of western blot signals between the anti-BCNT-C antibody and A305-624A-M. The supernatants (SUPs) and their pellets (PPTs) of T-REx or its G11 cells (His-tagged human BCNT/CFDP1 constitutively expressing clone) were prepared from each cell lysate by centrifugation at 25,000 × *g*. Equal amounts of protein (20 μg) were subjected to western blotting analysis with either anti-BCNT-C Ab or A305-624A-M (left panel). After obtaining their images, each filter was stripped and re-probed with the exchanged Abs (right panel).

### Assigning the candidate signal of mBcnt/Cfdp1 using ES mutant cells

To assign the appropriate western blot signal of endogenous Bcnt/Cfdp1 and evaluate its Abs, we utilized a mouse embryonic stem (ES) cell line that is listed as homozygous m*Bcnt/Cfdp1* mutant cells (i.e., double-knockout cell line) (Cfdp1-K1)[18]. The gene trap vector is inserted in m*Bcnt/Cfdp1* intron 5 (Fig 4A) (GenBank: accession number AG999723.1). As the BCNT-C domain is encoded by exons 6 and 7 (Fig 4A), the Cfdp1-K1 cell lysate could be used as a potential negative control for the validation of Abs generated with the BCNT-C domain as an immunogen, assuming the cells are homozygous m*Bcnt/Cfdp1* mutants.

**Fig 4.**
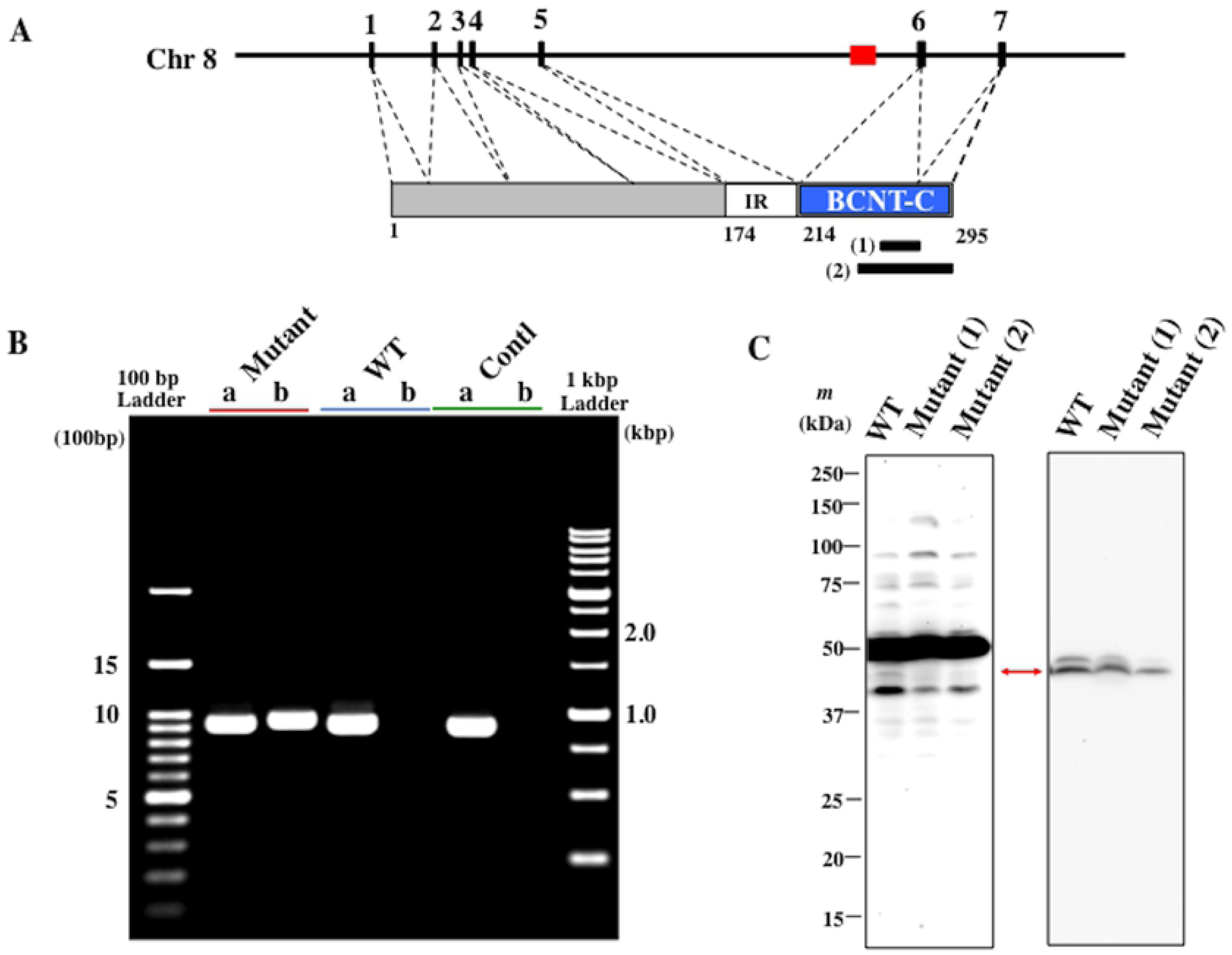
Assigning western blot signals of endogenous Bcnt/Cfdp1 using mutant ES cells. (A) Location of the inserted gene trap vector in the Cfdp1-K1 cell. Mouse *Bcnt/Cfdp1* consists of 7 exons, and the dashed lines show their corresponding regions to Bcnt/Cfdp1. A red box indicates the gene trap vector. Two black bars under Bcnt/Cfdp1 predict each location of immunogens for Ab production of anti-BCNT-C Ab (1) and A305-624A-M (2), respectively. (B) RT-PCR analysis of *Bcnt/Cfdp1* mRNA from Cfdp1-K1 and its parental cells. Using each cDNA from Cfdp1-K1 (Mutant), vdR2-4 (WT), or mouse brain (Contl), RT-PCR was carried out to examine the products that correspond to (a) the full ORF of m*Bcnt/Cfdp1* (928 bp) and (b) the fused region of m*Bcnt* exons 1-5 and a part of *hygromycin phosphotransferase* in the gene trap vector (948 bp), respectively. The products and DNA size ladder markers were accessed by separation in agarose gel, followed by staining with ethidium bromide. (C) Assessment of western blot signal of endogenous Bcnt/Cfdp1 using extracts of Cfdp1-K1 and its parental cells. Cell extracts from equal number (~2 × 10^5^ cells) of vdR2-4 (WT) or Cfdp1-K1 (Mutant) cells that had been serially passaged in the presence [Mutant (2)] or absence [Mutant (1)] of G418 and puromycin were subjected to western blot analysis with either anti-BCNT-C antibody (left filter) or 305-624A-M (right filter), respectively. A red arrow indicates a candidate signal of endogenous mouse Bcnt/Cfdp1.

First, we confirmed whether Cfdp1-K1 is knocked out on the expression of *Bcnt/Cfdp1* by reverse transcription-polymerase chain reaction (RT-PCR). After subculturing in the presence or absence of G418 and puromycin, which can delete the feeder layer cells, we prepared cDNAs from Cfdp1-K1 and vdR2-4 cells; the latter is a parent cell line of Cfdp1-K1. We then examined the target m*Bcnt/Cfdp1* mRNA by RT-PCR and analyzed their products by DNA sequencing. In the cDNA from Cfdp1-K1, we detected PCR products corresponding to not only the fusion gene coding *Bcnt/Cfdp1* exons 1-4 and *hygromycin phosphotransferase* derived from the gene trap vector but also the full-length *mBcnt/Cfdp1* ORF (Fig 4B). This result indicated that the gene trap vector was inserted adequately into intron 5, but Cfdp1-K1 cells were not *Bcnt/Cfdp1* double knockout. Then, a comparative transcriptome analysis by NovaSeq 6000 was carried out between Cfdp1-K1 (mutant) and vdR2-4 (wild-type), and it has been shown that 694 genes were differentially expressed with a 2-fold difference, 188 genes were upregulated, and 506 genes were downregulated (S1 Appendex and S2 Table). Among them, *Bcnt/Cfdp1* mRNA in the Cfdp1-K1 cells was reduced to 74.4% compared to the parent cells (101.75 vs. 136.79 FPKM). Besides, the mRNAs of several housekeeping genes that are frequently used as internal controls, the flanking genes *Bcar1/Cas* and *Tmem 170A*, were also significantly altered (S3 Table.).

Next, we examined whether anti-BCNT-C and A305-624A-M Abs detected the differential expression of Bcnt/Cfdp1 between mutant and parental cells in western blot analysis. Of the several bands detected by the anti-BCNT-C Ab, one ~45-kDa band was significantly reduced in the Cfdp1-K1 cells compared to the band intensity in vdR2-4 cells by both Abs (Fig 4C). These results suggest that the ~45-kDa band is a candidate signal derived from endogenous mBcnt/Cfdp1 in mouse ES cells.

### Validating anti-mBcnt-N Ab and assigning the mBcnt/Cfdp1 signal

As mentioned earlier, whereas the C-terminal region of BCNT members is highly conserved, the N-terminal region is variable among species. To further investigate whether the ~45-kDa band was a valid signal, we generated an anti-mBcnt/Cfdp1 Ab using a mouse-specific N-terminal peptide consisting of 16 amino acids as an immunogen (named anti-mBcnt-N Ab, Fig 5A).

**Fig 5.**
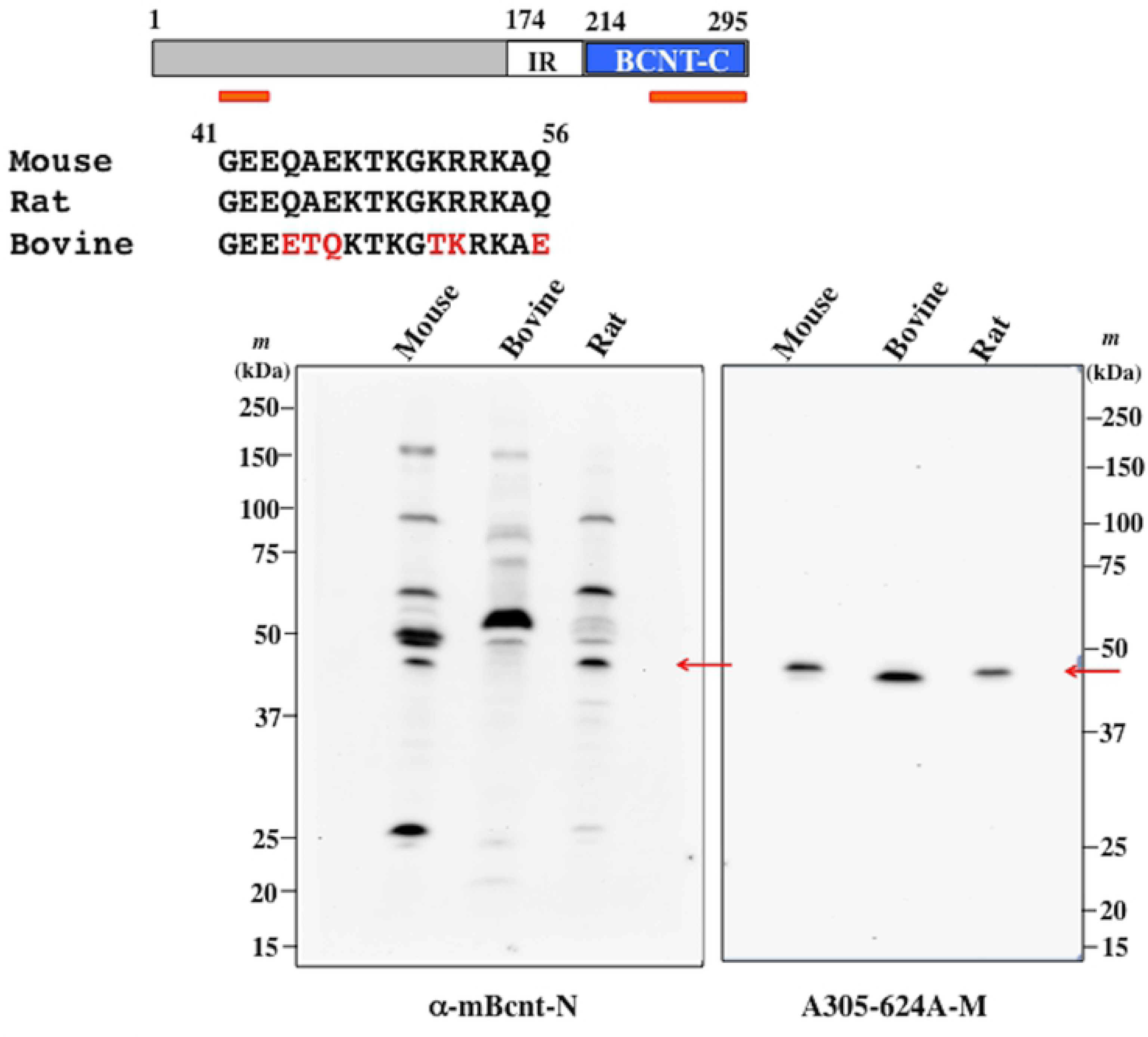
Validation of anti-mBcnt-N antibody. (A) Amino acid sequence alignment of the immunogen for anti-mBcnt-N Ab generation and its counterparts of rat and bovine. Mouse Bcnt/Cfdp1 is schematically shown in Fig 4A. The underlined two red bars present each location of the immunogens for the generation of anti-mBcnt-N Ab and A305-624A-M, respectively. The rat and bovine counterparts are aligned, and red letters indicate the different amino acids from the immunogen peptide. (B) Equal amounts of mouse and rat brain extracts (15 or 20 μg) and the enriched bovine placenta extract (30 or 40 μg) were separated on SDS/PAGE followed by western blot analyses with either anti-mBcnt-N Ab (left filter) or A305-624A-M (right filter). Each first Ab was used at a final concentration of 500 ng/mL and 1 μg/mL, respectively.

While the rat Bcnt/Cfdp1 has the same sequence as the immunogenic peptide derived from mBcnt/Cfdp1, the bovine counterpart has an entirely different amino acid sequence (Fig 5A), implying that the probability of cross-reactivity with anti-mBcnt-N Ab was expected to be very low. Indeed, we could use rat and bovine tissue extracts as potential positive or negative controls, respectively, to evaluate anti-mBcnt-N Ab specificity concerning endogenous Bcnt/Cfdp1 in western blot analysis. However, since it was difficult to obtain a bovine source containing high Bcnt/Cfdp1 content, we concentrated an extract of bovine placenta with Phos-tag agarose [19], which allows enrichment of phosphorylated protein (S3 Fig), and used it for the western blot analysis (Fig 5).

As a result, while A305-624A-M detected a ~45-kDa signal in both tissue extracts, anti-mBcnt-N Ab detected the band in mice and rats but not in cattle. This result strongly suggests that anti-mBcnt-N Ab specifically recognizes endogenous mBcnt/Cfdp1 despite the fact that a nonspecific cross-reaction with unknown proteins was observed. We further confirmed that the ~45-kDa signal was a valid target signal using another anti-hBCNT/CFDP1 Ab, 26636-1-AP (S4 Fig). Finally, we performed western blot analysis with two Cfdp1-K1 cell lysates prepared after passage in the presence or absence of G418 / puromycin using anti-mBcnt-N Ab or A305-624A-M. The results showed that both Abs recognized a ~45-kDa band with significantly reduced intensity in both Cfdp1-K1 lysates as compared to that in the parental cell lysate (Fig 6). From these results, we concluded that the ~45-kDa signal is derived from the endogenous mBcnt/Cfdp1.

**Fig 6.**
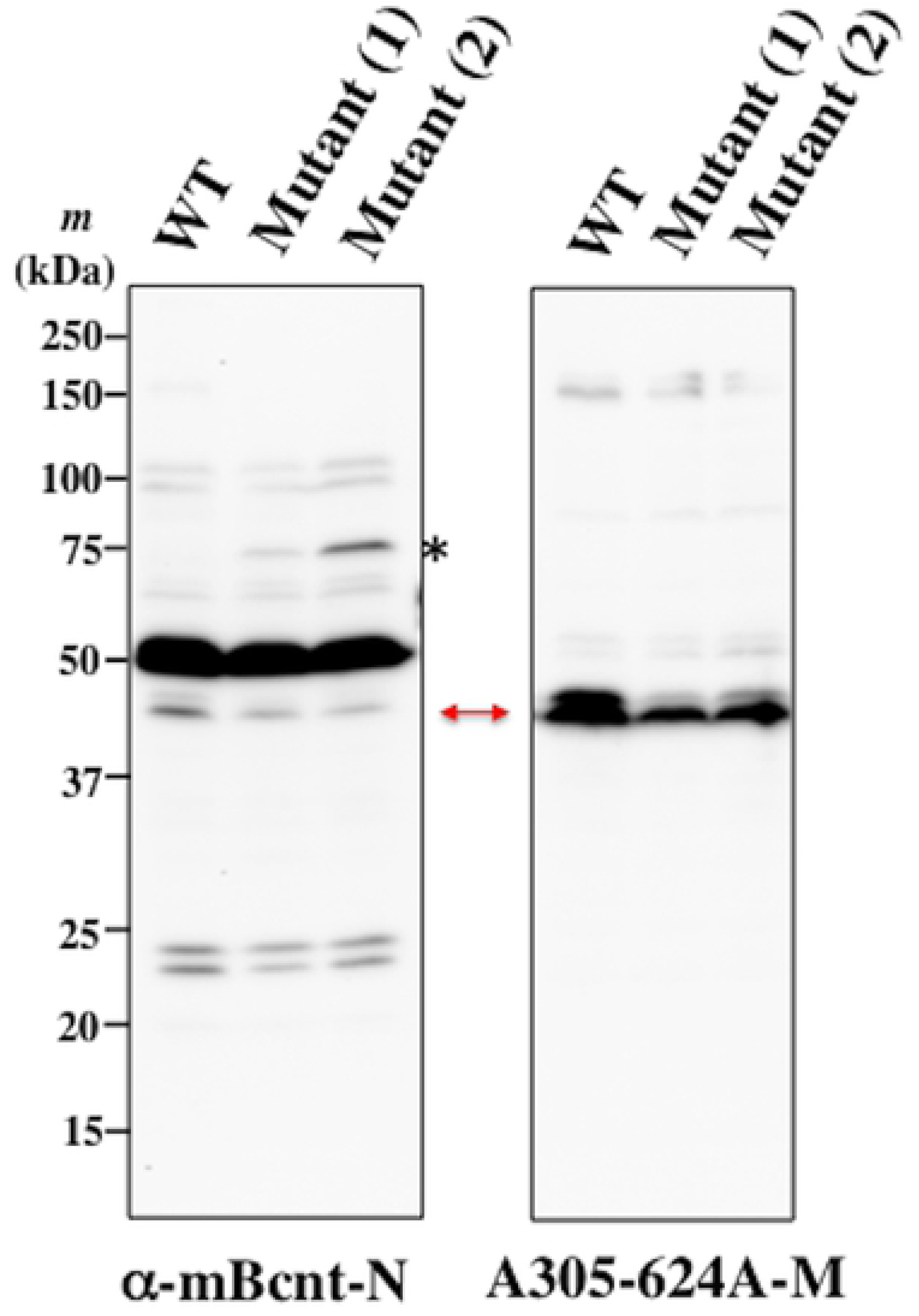
Assignment of western blot mBcnt/Cfdp1 signals. Equal amounts of cell extract (20 μg protein) of vdR2-4 (WT) or Cfdp1-K1 (Mutant) cells serially passaged in the presence [Mutant (2)] or absence [Mutant (1)] of G418 and puromycin were subjected to western blot analysis with anti-mBcnt-N Ab (left panel) or A305-624A-M (right filter). A red arrow indicates a signal of mouse Bcnt/Cfdp1. A band detected with anti-mBcnt-N Ab at ~75 kDa (shown as *) is probably the fusion protein of a part of mBcnt (derived from exon 1-5) and hygromycin phosphotransferase. It is 63.9 kDa as a calculated molecular mass but may run slowly on SDS/PAGE due to the acid stretch located in the N-terminal region of the mouse Bcnt/Cfdp1.

To confirm whether the ~45-kDa band detected by the both Abs is identical, the filter that had been detected with anti-mBcnt-N Ab was reprobed with A305-624A-M. The result indicated that the signal was identical.

### Expression of mBcnt/Cfdp1 in the early stage of brain development

RNA profiling data of the mouse and rat ENCODE (The Encyclopedia of DNA Elements) projects show that Bcnt/Cfdp1 mRNA expresses ubiquitously and preferentially in the early stage of development. However, recent studies have revealed pervasive discordance between mRNA levels and protein levels, especially in embryonic development [14]. Therefore, we examined mBcnt/Cfdp1 expression in the cerebrum of mouse and rat, focusing on developmental stages using the evaluated anti-Bcnt/Cfdp1 Abs above. The results showed that mBcnt/Cfdp1 preferentially expresses in the early stages and significantly decreased according to the postnatal stages in the rat cerebrum (Fig 7).

**Fig 7.**
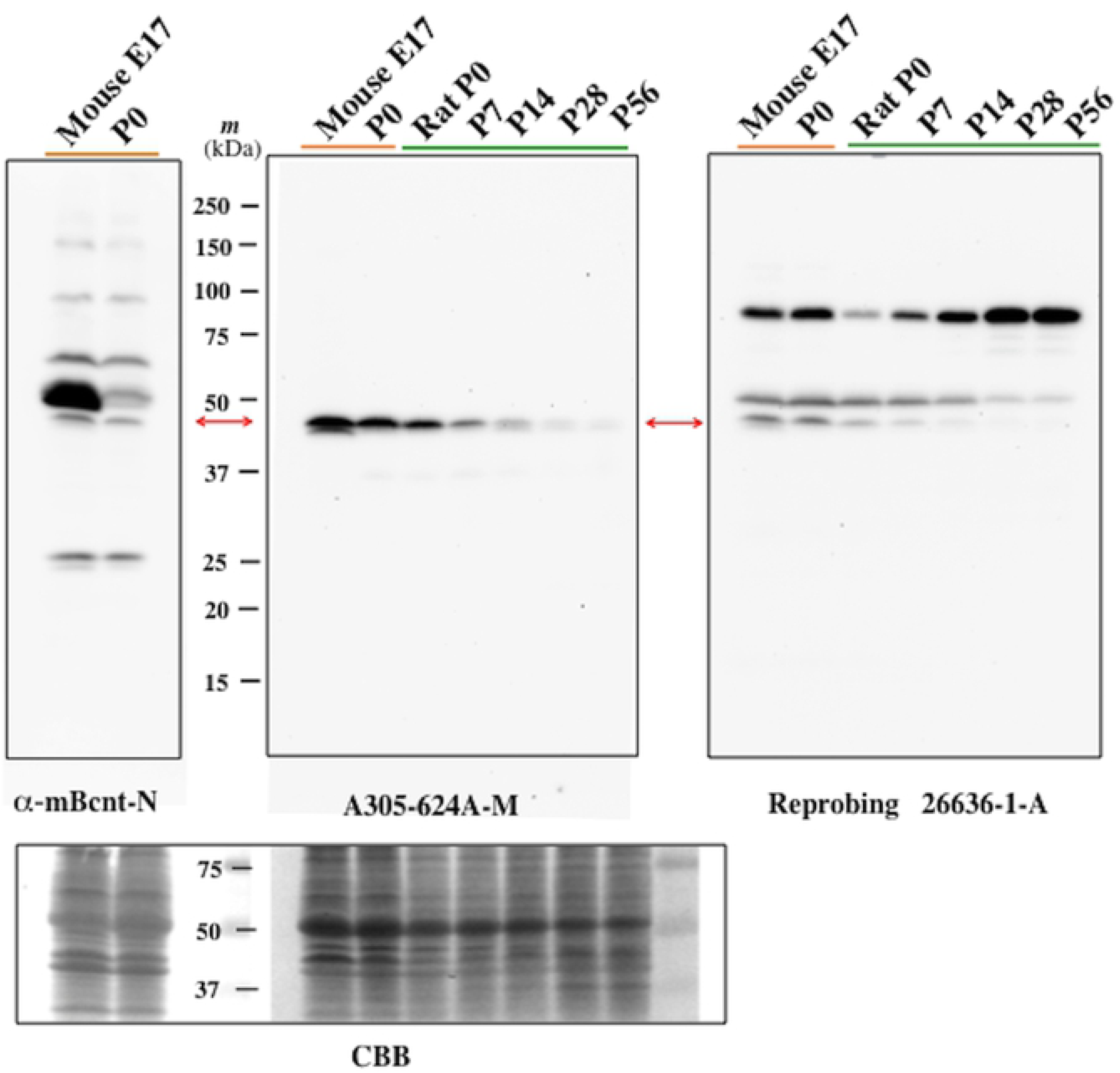
Bcnt/Cfdp1 expression of rodent brain in the early stage of development. Equal amounts (20 μg of protein) of cerebrum extracts of the embryo around 17 days (E17) and P0 mouse and rat postnatal day samples (denoted by P# on the top of each lane) were loaded and subjected to western blotting analysis with anti-mBcnt-N Ab (left panel) or A305-624A-M (middle panel). The latter filter was re-probed with another anti-BCNT/CFDP1 Ab, 26636-1-AP (right panel). The filters were finally stained with Coomassie Brilliant Blue (CBB) to check the amounts of loading proteins (bottom panel).

## Discussion

In this paper, we assigned the endogenous mBcnt/Cfdp1 signal in western blot, validated anti-mBcnt/Cfdp1Abs by utilizing various target-related materials, and showed that mouse and rat Bcnt/Cfdp1 expressed preferentially at early stages of brain development. Moreover, based on the problems encountered when attempting to solve these tasks in the present study, we discuss immune-cross reactions with off-target proteins from the viewpoint of immune reaction probability.

We generated F-mBcnt instead of His-tagged hBCNT/CFDP1 because the latter made it challenging to evaluate the specificity of anti-Bcnt/Cfdp1 Abs due to the difficulty to distinguish specific immune-positive signals from the high background of anti-His tag positive proteins [15]. Indeed, Nono (p54rnb)[20], which contains the HHQHHH sequence in the N-terminus region, could be isolated using anti-His-tag Ab-conjugated beads (S6 Fig). F-mBcnt appeared as a doublet, and the upper band was much more phosphorylated than the lower band-in particular, the phosphorylation of the Ser^246^ residue in the WESF motif of the BCNT-C domain was characteristic. The mBcnt/Cfdp1 doublet was probably caused by the presence or absence of Ser^246^ phosphorylation because this characteristic phosphorylation is very similar to that of His-tagged hBCNT/CFDP1, as previously described [15]. Though its biological significance is not yet clear, one interesting possibility is that this phosphorylation might be a determinant of the intracellular localization of Bcnt/Cfdp1.

We first evaluated a custom-made anti-BCNT-C Ab that detected a distinct band from those detected by anti-Flag Ab in F-mBcnt-expressing cell lysates (S1 Fig). We initially assumed that the band was hBCNT/CFDP1. By introducing another commercially available anti-hBCNT/CFDP1 Ab (i.e., A305-624A-M from Betyl laboratory), we compared the immune-positive signals detected with ant-BCNT-C Ab and found that their patterns were distinctively different in the parental cell extract. On the other hand, the exogenous hBCNT/CFDP1 in the extract of G11 clone were well recognized by both Abs. Therefore, we utilized a Cfdp1-K1 cell line that has been produced by a library of random mutations introduced by gene trap vector insertion in Bloom-deficient ES cells, selected for populations of homozygous mutant cells following mitotic recombination [18], and was listed as a cell line of mouse *Bcnt/Cfdp1* homozygous mutant (Japanese Collection of Research Bioresources (JCRB) Cell Bank_AyuK7D01). However, RT-PCR analysis of cDNA from Cfdp1-K1 revealed the presence of mRNA corresponding to the full-length ORF of the m*Bcnt/Cfdp1*. Indeed, a comparative analysis of transcriptome RNA sequencing between Cfdp1-K1 and its parent cells showed that *Bcnt/Cfdp1* mRNA levels were 74.4%. Splicing may efficiently occur by skipping the acceptor site in the trap gene vector, which is located in the intron 5 spanning over 50 kb (Fig 4A). Although the Cfdp1-K1 cell line was not *Bcnt/Cfdp1* double knockout, it is useful as *Bcnt/Cfdp1* knockdown mutant cells to detect a strong candidate signal at ~45 kDa. On the other hand, an attempt has been made to generate dog *Bcnt/Cfdp1* knockout MDCK (Madin-Darby Canine Kidney) cells by targeting its exon 1 according to the same strategy as the production of β- and γ-catenin double-knockout cells [21], but the expected cells were not obtained so far (W. Kobayashi, personal communication). Finally, we were able to assign the signal of endogenous mBcnt/Cfdp1 detected by two Abs raised against unrelated immunogens. One of the antigens is a mouse-specific N-terminal peptide, and the other is a peptide of BCNT-C domain, which is highly conserved in mammalian Bcnt/Cfdp1.

The following is evidences that the ~45-kDa band is the signal of endogenous Bcnt/Cfdp1 in western blot. First, among several signals detected by the anti-BCNT-C Ab, the ~45-kDa signal was significantly reduced in the western blot of the m*Bcnt/Cfdp1* mutant cell extract compared to the signal of the parent cell (Fig 4C). Second, the ~45-kDa signal was detected by two other antibodies (i.e., anti-mBcnt-N Ab and A305-624A-M), each of which was generated using mutually unrelated immunogens (Fig 6) as well as another anti-hBCNT/CFDP1 Ab, 26636-1-AP (S4 Fig). We confirmed the specificity of the anti-mBcnt-N Ab concerning the ~45-kDa signal by preparing enriched bovine Bcnt/Cfdp1 with Phos-tag agarose (Fig 5) as a potential negative control in a western blot.

The target band at ~45-kDa appears significantly smaller than signals reported by many available anti-Bcnt/Cfdp1 Abs, including our custom-made anti-BCNT-C Ab. The molecular behavior of ~50-kDa protein reported with many anti-hBCNT/CFDP1 Abs was significantly different from that of endogenous Bcnt/Cfdp1 (S1 Fig and Figs 3 and 4C); therefore, the 50-kDa signal is probably a common non-specific band(s). Since we used this anti-BCNT-C Ab, our initial conjectures regarding the 50-kDa signal (refer to abstract in [15]), the 43-kDa signal (glutamine synthetase, [17]) (refer to Fig 3C in [16] and Fig 1B in [22]) determined by western blot analysis, and the intracellular localization by immunostaing (refer to Fig 6b in [23]) were all misdirected. On the other hand, we did not recognize two bands at ~37 and ~19 kDa, which are shown in the catalog of A305-624A-M. Furthermore, although five isoforms of m*Bcnt/Cfdp1* have been reported (*Mus musculus*, NCBI Accession No. ID: 23837), we could not identify any isoforms during our western blot analyses.

It was evident that the anti-BCNT-C Ab detected a weak signal to the target molecule (Fig 4C). A similarly difficult situation must occur with many Abs, which is considered to be problematic (e.g., [24], [25]). Concerning the time-consuming validation of anti-Bcnt/Cfdp1 Ab using western blot, we consider the intrinsic issues that may have led to its inappropriate assignment and how these could be resolved. First, we should use the information of preferential expression of Bcnt/Cfdp1 mRNA—for example, mouse embryo brain is more suitable than the adult brain as screening samples for its Ab evaluation. Second, although we needed other independent anti-BCNT-C Abs and attempted to prepare them using the N-terminal region as immunogens, we failed to obtain the appropriate Abs that could be utilized at that time. Third, although non-specific band patterns detected by Abs raised against different immunogens are generally not identical, many available anti-Bcnt/Cfdp1 Abs including the anti-BCNT-C Ab and 26636-1-AP, commonly detected a relatively strong signal(s) near 50-kDa, and thus resulted in an incorrect assignment. Regarding the second and third points, the properties of Bcnt/Cfdp1 may be related since it mainly consists of structurally disordered regions. It is noteworthy that the epitopes from disordered antigens are smaller than those from the ordered counterparts and that they interact more efficiently with their Abs [26]. Thus, the anti-Bcnt/Cfdp1 Abs generated against most parts of the molecule as immunogens may cause cross-reaction with various proteins with high affinity, resulting in the poor quality of antisera. Fourth, complex migration of Bcnt/Cfdp1 on SDS/PAGE made it challenging to assign a valid signal. BCNT/CFDP1 as well as mBcnt/Cfdp1 is expressed as a doublet band and migrates slower on SDS/PAGE than expected from the calculated molecular mass (33.6 kDa of human BCNT/CFDP1 and 32.7 kDa of mouse Bcnt/Cfdp1). This feature may be mainly due to the acid stretch of the N-terminal region and the Ser phosphorylation of the BCNT-C domain [15]. Lastly, despite a lack of substantial evidence that *Bcnt/Cfdp1* is directly involved in craniofacial development, its attractive but misleading nomenclature (i.e., craniofacial development protein 1) may have caused confusion regarding its function in Ab providers and researchers. As a result, this may have prevented prior careful analysis of the molecule.

Off-target problems have been widely discussed in many Ab validation studies (e.g., [24], [25]), including specific in-depth efforts for Ab evaluation (e.g., [27]). Moreover, a strategy for Ab validation has been proposed [28]. However, the focus of this discussion appears to be blurred, at least regarding western blot analysis.

Immune cross-reactivity of Abs is based on a general chemical reaction determined by the reaction probability. Epitopes are conventionally divided into two categories: linear or sequential and discontinuous or conformational epitopes [29]. However, Abs do not recognize even linear epitopes as a series of amino acid residues but rather the physicochemical and stereochemical states that they constitute, and these properties as a whole constitute epitope [30, 31]. For example, the following two cases may reflect topological or stereochemical similarity of small environments that determine the common epitopes between completely different proteins: Bcnt/Cfdp1 and glutamine synthetase [17] and phosducin and β-actin [32], respectively. Thus, in principle, it is impossible to exactly match the properties of the epitope with their amino acid sequences, even in linear epitopes. Furthermore, it has been shown that linear epitope peptides that reveal apparent off-target binding at the peptide levels have a strict conformational component at the protein levels [33]. The classification of two types of epitopes is therefore not easily producible.

Western blot signals detected by a certain Ab are strongly influenced by the target concentration in test samples, as shown in Fig 3. The anti-BCNT-C Ab recognized the exogenously expressed His-tag hBCNT/CFDP1 but scarcely detected an endogenous counterpart. The result implies that the extract of cells overexpressing a target protein does not qualify as a positive control for Ab evaluation in some experiments. However, a good Ab means that it is useful to reveal new evidence regardless of the type of experiment, which depends mainly on the target concentration to be analyzed and the extract preparation method (Figs 3 and 5 and [25]). The experimental materials are quite different with respect to their species, tissues, and developmental stages. Besides, when the target gene has a strong influence on other genes or when the target is a lowly expressed protein, some efforts may be required to assign their appropriate target signals even using lysates from knockout or knockdown cells as a negative control. As shown by the effect of Bcnt/Cfdp1 knockdown in Cfdp1-K1 cells, it is not uncommon that a single gene mutation dramatically alters the expression levels of other genes (S3 Table and S4 Table). Furthermore, it is possible that many proteins showing similar mobility with the target molecule on SDS/PAGE overlap the migration region of the target molecule, making the signal assignment challenging [34]. There is no guarantee that the once validated Ab will work with other samples.

Of course, what we can do is generate antibodies using many unrelated immunogens and also eliminate troublesome off-targets by rigorously screening candidate antibodies (e.g., [28]). However, it is not necessary to be concerned with a single signal, and even if a few off-target signals are detected, these kinds of Abs can be useful if we have evaluated their limitation correctly, as shown in Fig 5 and in many reports (e.g., [35]). Abs do not act within the confines of all-or-nothing modes via specific reactions; therefore, it is critical to understand the efficacy and limitations of the antibody used in any experiments.

Although sensitivity of western blot analysis to crude extract is much better than that of mass spectroscopy, these tools are fundamentally different—that is, individual Ab is not a tool for identifying a molecule but rather a tool for checking a contradiction. Recently, mass analyses have become remarkably advanced and widely available. Thus, it is now much easier to identify molecules that have been considered to be false targets due to wrong-cross reaction with Abs. These trials may provide byproducts of the excellent Abs, such as conformation-specific antibodies against the proteins.

As a reliable anti-Bcnt/Cfdp1 Ab (i.e., A305-624A-M) becomes clearer at present, it is possible to characterize endogenous Bcnt/Cfdp1 more accurately, including subcellular fractionation and tissue distribution. On the other hand, the comparative transcriptome analysis of Cfdp1-K1 cells revealed that only a 25% decrease in Bcnt/Cfdp1 mRNA resulted in a marked up-regulation or down-regulation of many genes (S1 Appendix, S2 Table), though the confirmation is required by generating their revertant cells by removing the mutagenic vector sequences through Flp-FRT recombination [18]. Among them, it is noteworthy that mRNA expression of intermediate filaments of keratins 8, as well as 18 and 19, was dramatically suppressed —that is, 1% relative to the parental cells (S4 Table, [36]). Given the function of Swc5, which is required under stressed conditions and conditions requiring rapid transcription ([*Saccharomyces* Data Base], [6], and [7]), Bcnt/Cfdp1 may play an essential role for maintaining cell homeostasis, especially in processes such as developmental stage, cell differentiation, and DNA damage repair. The Cfdp1-K1 cell line may serve as a stable Bcnt/Cfdp1 knockdown cell in elucidating the functional role of Bcnt/Cfdp1 in the early developmental stage by using western blotting analysis with further reliable anti-Bcnt/Cfdp1 Abs.

## Materials and Methods

All of the reagents and materials, and primers used are listed as S5 Table and S6 Table, respectively.

### Ethical approval

All of the genetic recombination experiments and all of the animal experiments in the present study were approved by the Genetic Recombination Experiment Safety Committee and the Animal Care and Use Committee, respectively, of Tokushima Bunri University. All of the experiments were performed in accordance with NIH Guidelines for the Care and Use of Laboratory Animals.

### Cell culture

T-REx-293 cells (a HEK293 cell derivative, T-REx) and all clone/subcolonies including G11 clone cells that constitutively expressed His-tag hBCNT/CFDP1 [15] were routinely maintained in DMEM-GlutaMAX-1 (DMEM), which contained 10% Fetal calf serum, 50 μg/mL gentamicin, and with or without G418 (0.5 mg/mL) in 5% CO_2_ incubator. For subculturing, the medium was aspirated completely, and the cell layers (2 ml culture on 35-mm dish) were incubated with 0.75 mL accutase for 5 min at room temperature. Then, they were homogenized by pipetting using a 1000-μL pipet tip, and 40-60 μL of the suspension was directly plated on the dish preincubated with the 2 mL culture medium for at least 30 min, and G418 was added the next day when needed. For transfection or preparation of cellular protein extract, the cell layers (5 mL culture on 60-mm dish) were washed with 5 mL prewarmed Hepes buffered saline (HBS, [10 mM Hepes-NaOH, pH 7.5, 150 mM NaCl], treated with 1.5 mL accutase, homogenized as described above, and then 1 mL medium was added. The cell suspension was transferred to a 15-mL tube, and the dish was washed with another 1-mL medium followed by combining the suspension within a total 10 min (total ~2.5 mL). After taking out 20 μL to estimate the cell number with a disposable Hemocytometer (Watson Bio Lab), the suspension was centrifuged (Sakuma Model RSL-IV; 1, 000 rpm, 1 min, room temperature) and resuspended in the culture medium for further study. TrypLE Express was also used in the earlier stage of the study in isolation of T-Rex-derived colonies that constitutively expressed Flag-MCS or Flag-tagged mBcnt (see below).

### ES cell culture

Cfdp1-K1 and vdR2-4 cells were obtained from Japanese Collection of Research Bioresources (JCRB) Cell Bank and grown in ESGRO Complete Clonal Grade Medium plus GSK3β Inhibitor (50 μl/100 mL) supplemented with gentamicin (50 μg/mL) in a 35-mm dish that was precoated by incubation with recombinant human Laminin (iMatrix-511) at a final concentration of 5 μg/mL in PBS for either 2 h at room temperature or 1 h at 37 °C. For subculturing, the medium was aspirated completely, and the cell layers were washed with 2 mL of prewarmed HBS and incubated with 0.75 mL accutase at room temperature. After 5 min, the cell layer was homogenized by pipetting using a 1000-μL pipet tip, and 1-mL DMEM containing 0.1% polyvinyl alcohol (DMEM-PVA) was added. The suspension was then transferred into a 15-mL tube and the dish was washed with another 1 mL of DMEM-PVA and the suspension was combined (total ~2.7 mL) within a total of 10 min. After taking out 20 μl for counting the cell number, the suspension was centrifuged (700 rpm, 3 min, room temperature) and resuspended in the new medium and seeded at the density of ~ 2.5 × 10^5^ cells per 35-mm dish. For the preparation of protein extract and total RNA, the cell suspension obtained above were twice washed with chilled HBS, dispensed at ~1 × 10^6^ cells per 1.5-mL tube, and centrifuged (100 × *g*, 5 min, 4°C). After removing the buffer completely, the pelleted cells were softly vortexed, snap-frozen in liquid N_2_, and stocked at −80 °C until use.

### Generation of T-REx colonies expressing Flag-mBcnt

T-REx cells (~2 × 10^6^ per 100-mm dish in 10 mL medium) were cultured for 20 h, and each 5 μg of pcDNA3.1 plasmid carrying Flag-MCS or Flag-*Bam*H1-mBcnt cDNA was added in the culture using Lipofectamine 3000 (5 μL in 250 μL Opti-MEM) according to the manufacturer protocol. Just before transfection, each 5 mL medium was once removed, transfected, and the medium that had been saved was back to the culture 4-6 h after transfection. After culturing for a total of 44-48 h, each cell layer was washed with 10 mL prewarmed HBS and treated with 1.5 mL of TrypLE Express for 10 min at 37 °C, harvested using 1 mL of medium in a 15-mL tube, centrifuged and resuspended in a 1-mL medium.

The number of each cultured cell was counted, and ~60 cells in 50 μL were seeded in a 48-well plate preincubated with 150-μL medium. After 4 hours, G418 was added to a final concentration of 0.75 mg/mL. Colonies growing in 48-well plates were sequentially expanded to 12-well plates (on day 15 after media change twice) and 35-mm dishes (on day 21 after media change twice). Each colonies was then maintained in the presence of 0.5 mg / mL G418.

### Protein extracts of T-REx cells and their colonies

After washing with chilled HBS (10 mL/100-mm dish) followed by removing the buffer completely using a piece of filter paper, the cells were homogenized with a cell scraper (17-mm width) in 0.5 mL lysis buffer [20 mM Hepes-NaOH, pH 7.5, 150 mM NaCl, 1 mM 2-mercaptoethanol, designated L_buffer] supplemented with both inhibitors of proteinases and phosphatases. The homogenate was transferred into a 1.5-mL tube, sonicated by a Bioruptor (BM Equipment) in an ice-water bath (10-s pulses repeated 15 times at 10-s intervals) and centrifuged (25,000 × g, 30 min, 4 °C) using a centrifuge (Kubota 3780, rotor AF-2536A). After taking out five μL for measurement of protein concentration, the supernatants were aliquoted, snap-frozen in liquid nitrogen, and stored at −80 °C until use. On the other hand, the pellets were dissolved in 50-μL lysis buffer containing SDS [1%SDS, 1 mM EDTA in 10-mM Hepes-NaOH, pH 7.5, designated LS_buffer], sonicated (3 × 10-s pulses at 10-s intervals) and centrifuged (10,000 × *g*, 1 min, 4°C). After taking out 5 μl for measurement of protein concentration, the sample was boiled in SDS/PAGE sample buffer.

### Isolation of F-mBcnt by anti-Flag Ab-conjugated agarose beads

The frozen supernatant of E3 colony was thawed, and Nonidet P-40 (NP-40) was added to a final concentration of 0.05%. After sonication for 30 s in the ice-water bath (3 × 10-s pulses at 10-s intervals) followed by centrifuging at 10, 000 × *g* for 1 min at 4 °C, the sup (1 mL of 1.2 mg) was mixed with anti-Flag-tag agarose beads (20 μL settled volume) in a 1.5-mL siliconized tube and incubated in a rotary shaker for 2 h at 4 °C. The mixture was centrifuged for 30 s using a swing type centrifuge (Swing Man, Type ATT-101), and the supernatant was saved as the unbound fraction. The pellet was suspended in 100-μL of L_buffer containing 0.05% NP-40 plus inhibitors and transferred to a spin column (0.8 mL size) using a 200-μL wide-bore tip, and the tube was once more washed with 100 μL of L_buffer plus NP-40, and the suspension was recovered to the spin column. The through-flow fraction obtained by centrifugation was stocked as the first wash fraction. The agarose in the column was washed twice with the same buffer, followed by being washed once more with HBS, and then the bound proteins were eluted with 50 μL of Flag (DYKDDDK) peptide (150 μg in HBS) by incubation at 4 °C for 30 min (Eluate #1) and another 5 min (Eluate #2), sequentially. The agarose in a column was further treated with 50 μL of glycine-HCl (50 mM, pH 2.5), and its eluate and the agarose in the column were immediately neutralized with 2 M Tris. Finally, 50 μL of 1 × SDS/PAGE sample buffer was added to the column, vortexed, and boiled for 5 min. All of the fractionated samples except the fraction eluted with SDS/PAGE sample buffer were boiled in 1 × SDS/PAGE sample buffer by mixing with 4 × SDS/PAGE buffer. Each five μL (3.75 μL of the net eluate) were separated on 1.25 % SDA/PAGE and followed by western blotting analysis using anti-Flag Ab. For LC-MS/MS analysis, Eluates #1 and #2 described above (each ~40 μL) were concentrated with acetone according to the protocol (http://tools.thermofisher.com/content/sfs/brochures/TR0049-Acetone-precipitation.pdf). The dry pellet was once dissolved in 6 μL of 10-fold diluted LS_buffer, and two μL of 4 × SDS/PAGE sample buffer was added followed by boiling for 3 minutes. After separating the sample for 10 minutes longer than usual on 12.5% gel SDS/PAGE, the gel was fixed with 50% MeOH-10% acetic acid solution for 20 minutes, stained with 0.25% CBB for 10 minutes, and then de-stained in 10% MeOH-7% acetic acid solution.

### Protein extracts from mouse and rat brain

Cerebrum and cerebral cortices were dissected from mice (P0 of C57BL/6J, male) and rats (P0-P56 of Wistar rat, male) after euthanasia under anesthesia, respectively. These isolated samples were snap-frozen in liquid N_2_ and stored at −80 °C until use. Frozen samples were crushed with a hammer on dry ice and immediately transferred to a glass-Teflon homogenizer with 1 mL of LS_buffer per 100 mg of samples. Then, tissues were homogenized at 600 rpm using a digital homogenizer and boiled for 5 min. These homogenates were sonicated (12 × 10-s pulses at 20-s interval) and centrifuged (28,000 × *g*, 30 min, 20 °C). In the case of a mouse embryo on day around 17, a frozen piece (~40 mg) was wrapped with aluminum foil, crushed with pliers with the help of liquid nitrogen, transferred to a 1.5-mL BioMasher, and soaked in chilled 200 μL of LS_buffer. Then, the tissues were homogenized in ice, boiled for 5 min, sonicated for 2.5 min (15 × 10-s pulses at 10-s intervals), and centrifuged (15,000 × *g*, 10 min, 20 °C). The supernatants were used for western blot analysis. Protein concentrations in the supernatants were determined using a Bicinchoninic Acid (BCA) protein assay kit with bovine serum albumin as a standard.

### Enrichment of bovine Bcnt/Cfdp1 by Phos-tag agarose

Bovine placenta (from Holstein Day 116) is a gift from Dr. Kazuyuki Hashizume (Iwate University, Morioka) and was stored in liquid nitrogen until use. Each piece (~30 mg) was further subdivided with a razor blade and extracted in two ways: one for the whole extract preparation and the other for the concentration of Bcnt/Cfdp1 content. The minced pieces were placed in 1.5 mL of BioMasher, soaked in cold 200 μL of L_buffer and homogenized, and then 10 μl of 20% SDS was mixed, followed by boiling for 5 minutes. The extract was sonicated for 2.5 minutes (15 × 10-s pulses at 10-s intervals), centrifuged (15,000 × *g*, 10 minutes, 22 °C.), and the supernatant was used as a whole extract. For enrichment of Bcnt/Cfdp1, the fined tissue piece in the 1.5-mL BioMasher was homogenized in 100 μL of chilled RIPA buffer [20 mM Tris-HCl, pH 7.5, 150 mM NaCl, 0.5% sodium deoxycholate, 1% NP-40, 1 mM EDTA] plus both inhibitors of proteinases/phosphatases. After transferring the homogenate into a new 1.5-mL tube, the homogenizer was washed with another 100 μL of RIPA buffer, and the solution was combined and centrifuged (15, 000 × *g*, 10 min, 4 °C). After taking out 5 μL for measurement of protein concentration, the supernatant was aliquoted, snap-frozen in liquid N_2_ and stored in −80 °C until use (total 1.4 mg protein). After dilution of the supernatant with RIPA buffer, 100 μL (200 μg) was enriched by Phos-tag agarose (100 μL settled volume in a spin column, 0.8 mL size) according to the manufactured protocol with the following modification; use of a swing-type centrifuge (Swing Man, Type ATT-101) and softly tapping the spin column during washing (200 μL, three times) and elution (100 μL per tube three times). Each eluate was precipitated with TCA at a final concentration of 20% and washed twice with 50 μL of acetone according to the protocol (http://www.its.caltech.edu/~bjorker/TCA_ppt_protocol.pdf). After heating at 95 °, 50 μL of SDS/PAGE sample buffer was added and solubilized by a mixer (Tomy MT-360) for 30 min at room temperature. For larger preparation, the above 700 μg of the extract was diluted to 350 μL with RIPA buffer, applied to a 350-μL settled volume of Phos-tag agarose in a spin column. After rinsing with 0.5 ml of washing buffer three times, the bound proteins were eluted with 0.5 and 0.45 mL of elution buffer sequentially into one tube, as described above. After precipitation with TCA, the pellets were mixed with 50 μL of LS_buffer for one hour. The total recovered protein was 112 μg, with a yield of 16 %. On the other hand, to estimate the approximate Bcnt/Cfdp1 content in the pellet (~20 μL) of the above centrifugation (15,000 × *g*, 10 min, 4°), 6.7 μL of 4 × SDS/PAGE sample buffer was added, mixed vigorously, boiled for 5 min, and sonicated for 2. 5 min (15 × 10-s pulsed at 10-s intervals).

### Immunoblotting

Procedures of SDS/PAGE of 12.5% or 15% gel and blotting onto membranes were mostly the same as previously described [15, 17]. The blotted PVDF membrane was blocked in 5% skim milk in TBT buffer [10 mM Tris-HCl, pH 7.6, 150 mM NaCl, 0.1 % Tween 20] for 2 h at room temperature or overnight at 4 °. The first Ab was incubated for 2 h at room temperature or overnight at 4 ° and the second Ab was treated for one h at room temperature. All immunoreactivity was visualized by chemiluminescent using horseradish peroxidase (HRP)-conjugated secondary antibodies that are listed in the S5 Table. Their image was detected by a scanner (GeneGenome, Syngene BioImaging) using ImmunoSTAR Zeta or LD as a substrate. For re-probing, the bound antibodies on the used membrane were stripped by incubation of the filters in a solution of 62.5 mM Tris-HCl (pH 6.8), 2% SDS, and 100 mM 2-mercaptoethanol with stirring for 30 min at ~60 ° on hot block, followed by washing three times with TBT buffer.

### Mass spectroscopy analysis

The upper and lower bands of a doublet band that were detected by CBB staining were cut out from the gel and digested with API, AspN, and chymotrypsin, and each digest was analyzed by nano-LC-MS/MS using a Q Exactive mass spectrometer as described previously [15]. The quantification of peptides derived from the upper and lower bands was carried out using a label-free quantification method using Proteome Discoverer Ver 2.2.0.388 (Thermo Fisher Scientific).

### Isolation of total RNA

The frozen ES cells were homogenized with Trizol reagent, and total RNAs were purified using PureLink RNA Mini kit according to the manufacturer protocol [17]. The purified total RNAs were treated with TURBO DNase to eliminate contaminating genomic DNA, extracted with phenol/chloroform/isoamyl alcohol (pH 5.2), and re-purified using RNA Clean & Concentrator -25 kit. The concentration of the total RNAs was determined by the absorbance at 260 and 280 nm using NanoDrop One (Thermo Fisher Scientific), and the quality was estimated using a 2100 Bioanalyzer (Agilent). Two RNAs with the high quality shown below were subjected to RNA sequencing (Macrogen Japan Corp., Kyoto).

| RNA source | Concentration (ng/μL) | A260/A280 | RIN |
| --- | --- | --- | --- |
| Cfdp1-K1 (Bcnt/Cfdp1 Mutant) | 454.6 | 2.07 | 9.9 |
| vdR2-4 (Wild type) | 366.8 | 2.03 | 9.6~9.9 |

### Reverse transcription-PCR and plasmid construction

cDNAs were synthesized by Superscript III SuperMix according to the manufacturer protocol using oligo-(dT) 20 primer from purified total RNAs of Cfdp1-K1, vdR2-4 cells, and whole mouse brain of a P56 C57BL/6J male as previously described [17]. The full-length ORF of m*Bcnt* or the fragment of m*Bcnt* exons 1-5 fused with *h*ygromycin phosphotransferase were amplified from each cDNA (1 ng as a total RNA) by PCR using KAPA HiFi HotStart DNA polymerase under the following cycling conditions: denaturation at 95 °C for 3 min, followed by 35 cycles of 98 °C for 10 sec, 68 °C for 15 sec, and 72 °C for 90 sec. The primer sequences are listed in S6 Table. After confirmation of their amplicon size by 0.8% Tris-Acetate EDTA agarose gel electrophoresis and purification by Wizard SV Gel and PCR Clean-Up System (Promega), the sequences of their ORFs were confirmed using BigDye Terminator V3.1 Cycle Sequencing Kit (Thermo Fisher Scientific). For the construction of Flag-mBcnt expression plasmid, the PCR product was inserted into a mammalian expression vector, Flag-MCS-pcDNA3.1 (Accession No. LC311018), using restriction enzymes *Bam* HI and *Xho* I as previously described [17].

### Transcriptome analysis

The following is from a report of HN00101712 (Macrogen Corp Japan, S1 Appendix). The two cDNA libraries from RNAs of Cfdp1-K1 or dvR2-4 cells were prepared using the TruSeq Stranded mRNA LT Sample Prep Kit (Illumina). Their sequences were obtained using NovaSeq 6000 S4 Reagent Kit by a Nova Sequencing system (S1 Appendix, S2 Table). Paired-end reads (read length 101) were mapped to a mouse reference genome (UCSC GRCm38.p4/mm10, annotation RefSeq_2017_06_12). After trimming, 98.81% were mapped on 45,262,214 cleaned reads from Cfdp1-K1 RNA, while 99.09 % were mapped on 52,245,252 cleaned reads from vdR2-4 RNA.

## Supporting information

Supplemental Fifure 6

## Acknowledgments

We wish to thank Drs. Kyoji Horie and Eiji Kinoshita for providing technical suggestions and useful discussions on ES cell culture and Phos-tag agarose handling, respectively. We thank Drs. Kazuyoshi Hashizume and Takeharu Masaki for giving us bovine tissues and *Achromobacter* protease I, respectively, Dr. Wakako Kobayashi for sharing the interest in generating Bcnt/Cfdp1 knockout cells, and Dr. Sang Wan Kim for conducting the transcriptome analysis. We are grateful to Drs. Salvador Eugenio Caoili, Christopher A. MacRaild, and Brian McWilliams for their particularly useful discussions and opinions. We also thank Drs. Ed Luk, Takashi Yasuda for discussions on the chromatin remodeling complex and on repair of damaged DNA, respectively. We were indebted to Professor Hiromi Nochi for her encouragement.

