## Supplementary figures and images for "Assessment of a western blot signal for the Bcnt/Cfdp1, a tentative component of Srcap chromatin remodeling complex; trial to overcome off-target problems"

### Supplemental Fifure 6

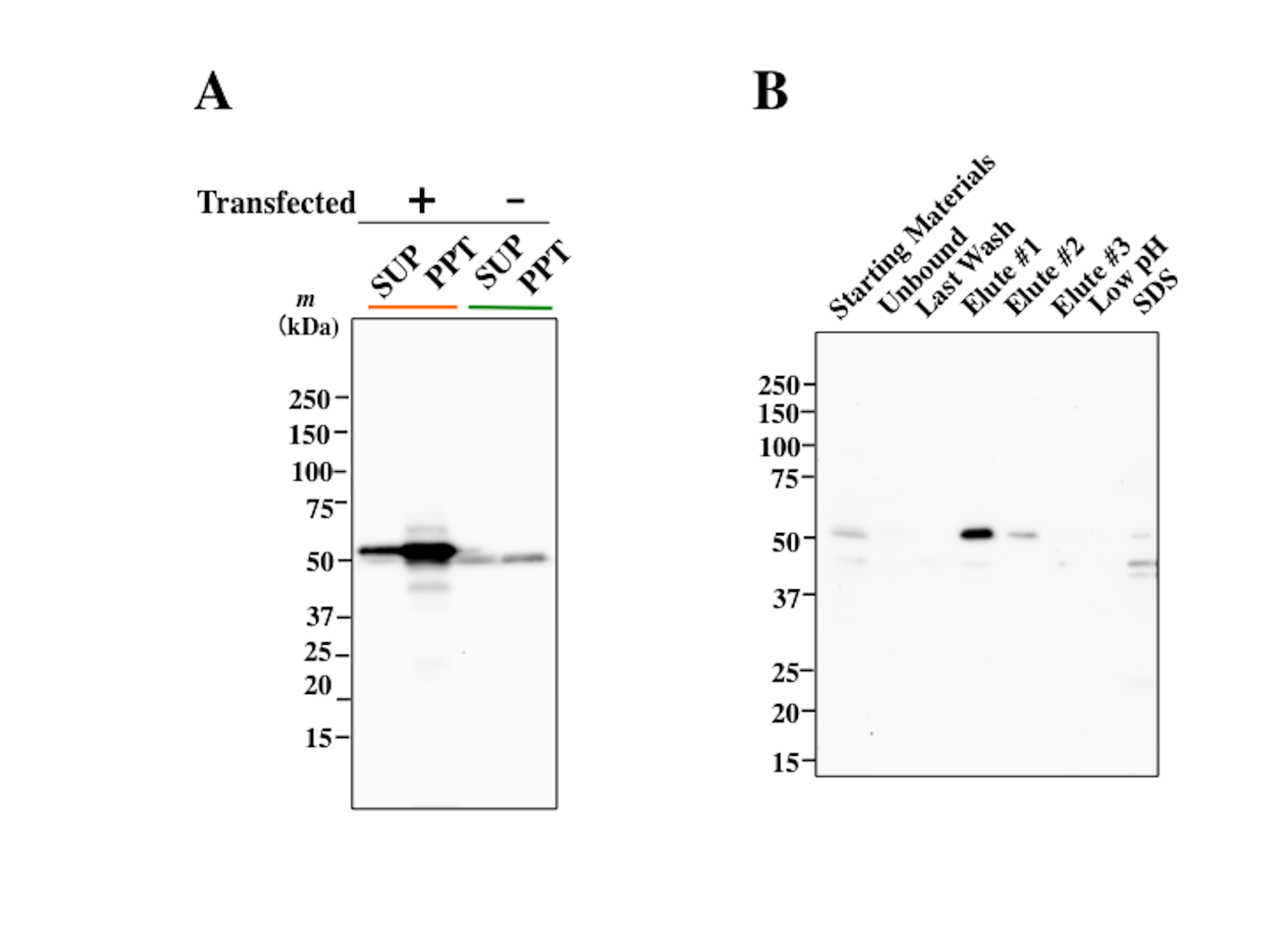

### Supplemental Figure 1

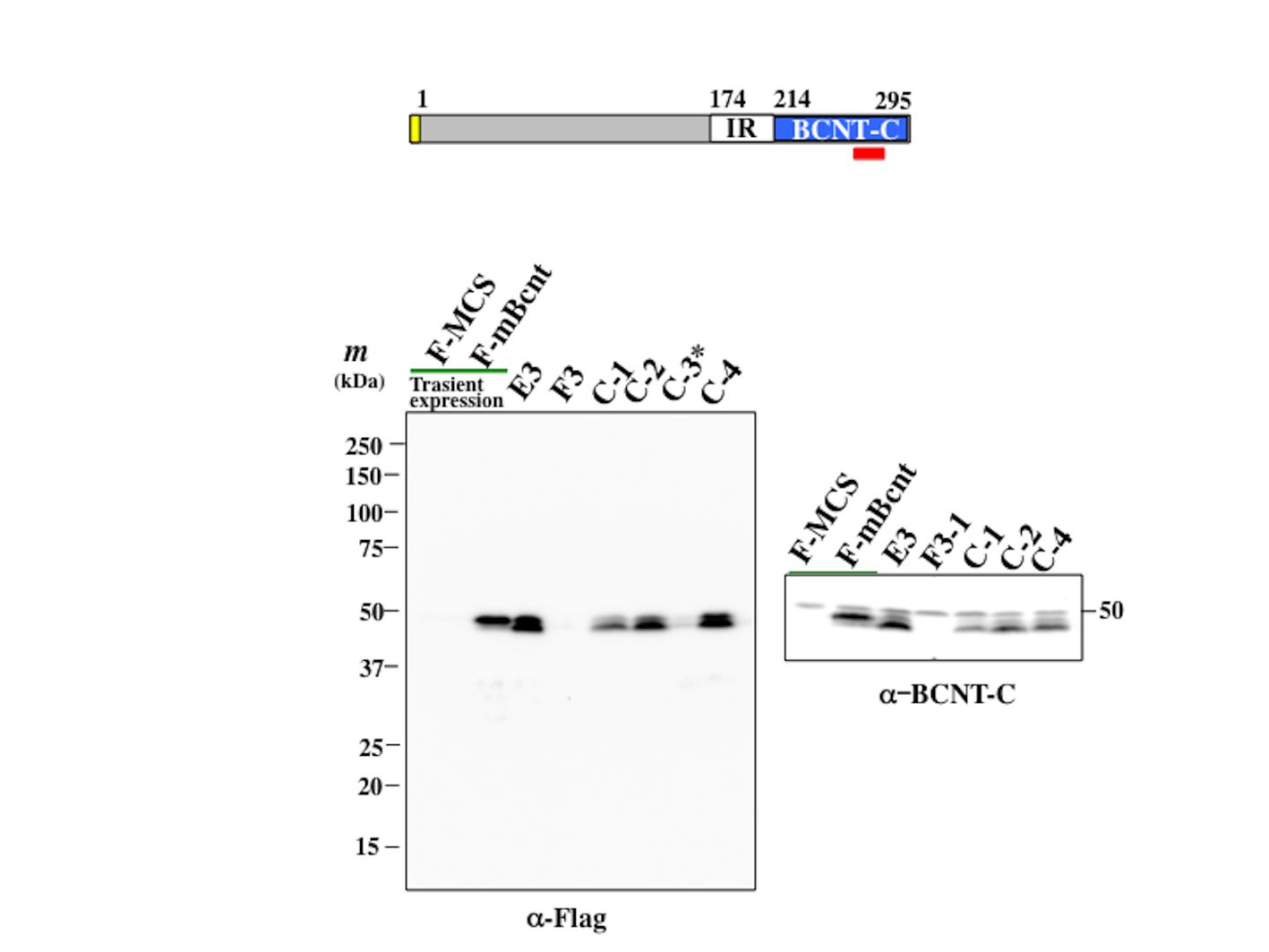

### Supplemental Figure 2

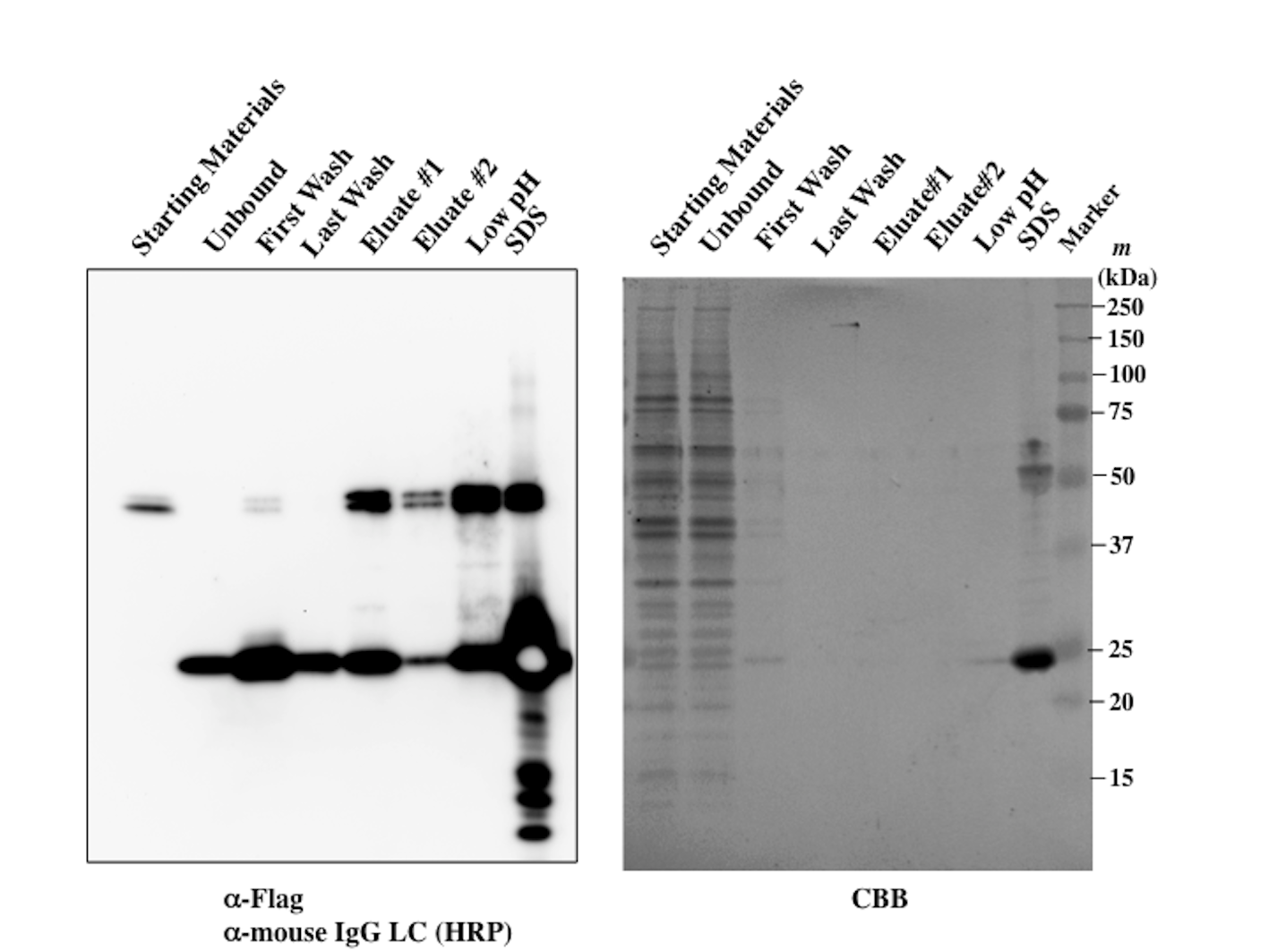

### Supplemental Figure 3

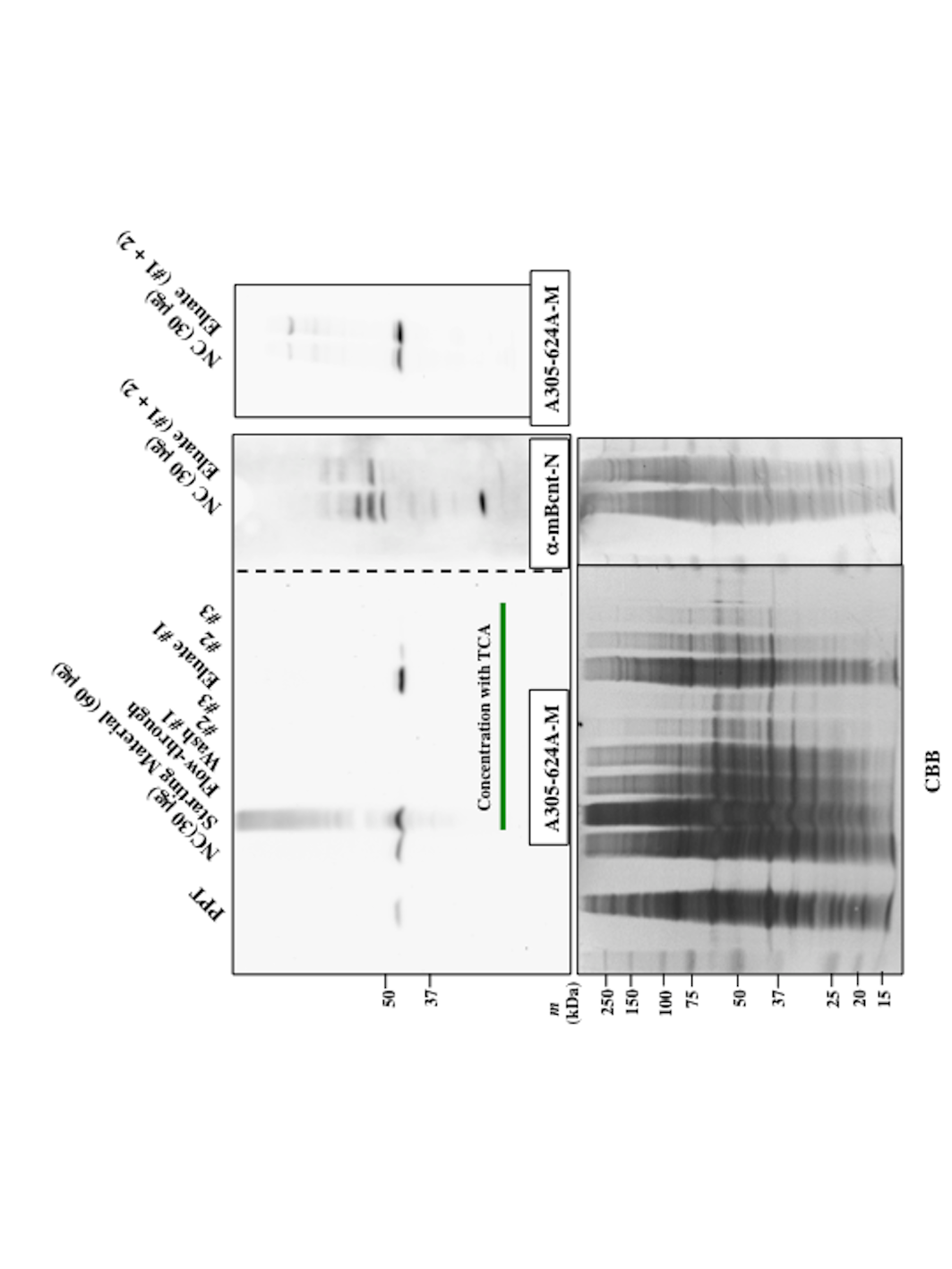

### Supplemental Figure 5

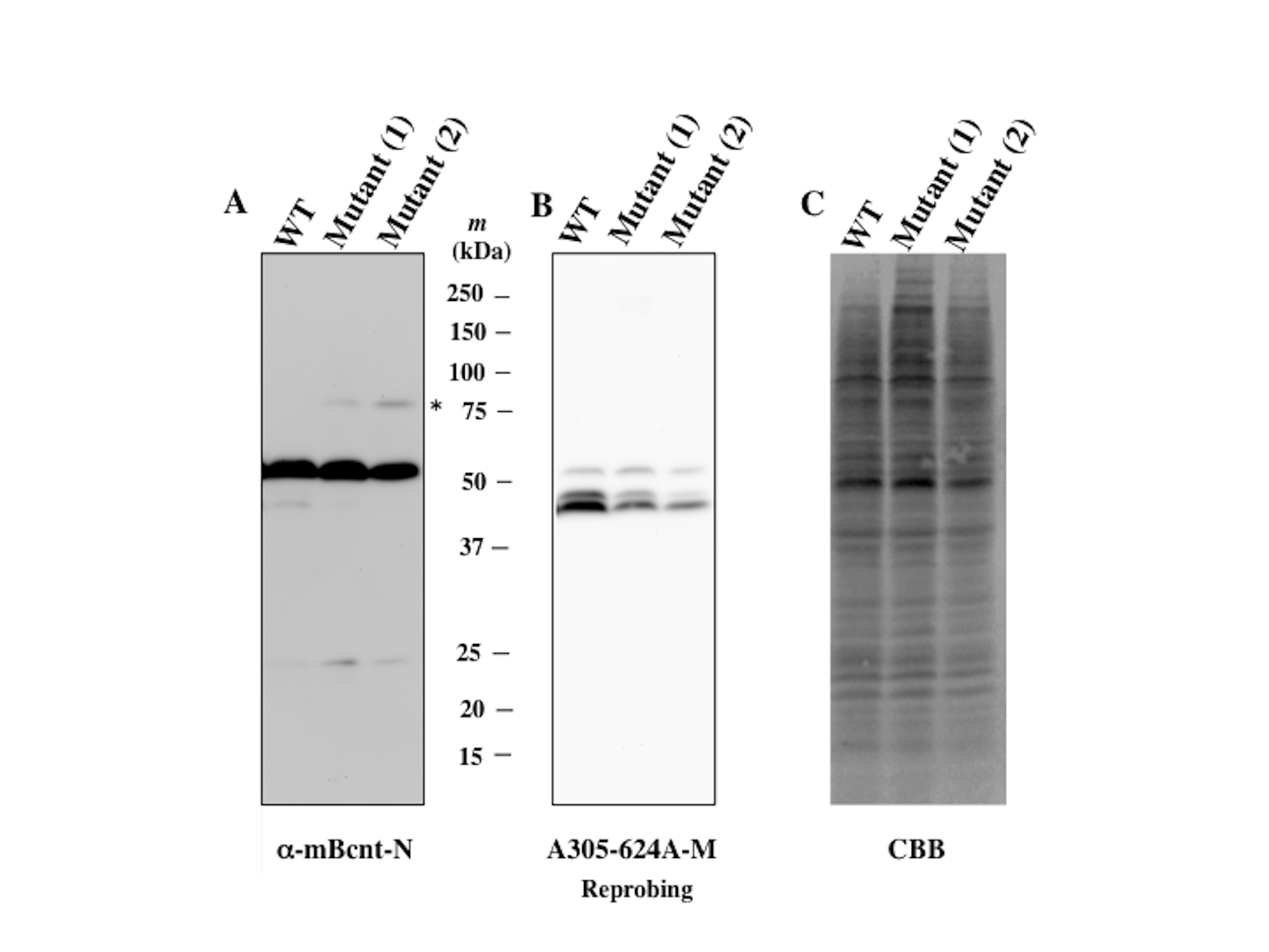

### Supplenetal Figure 4

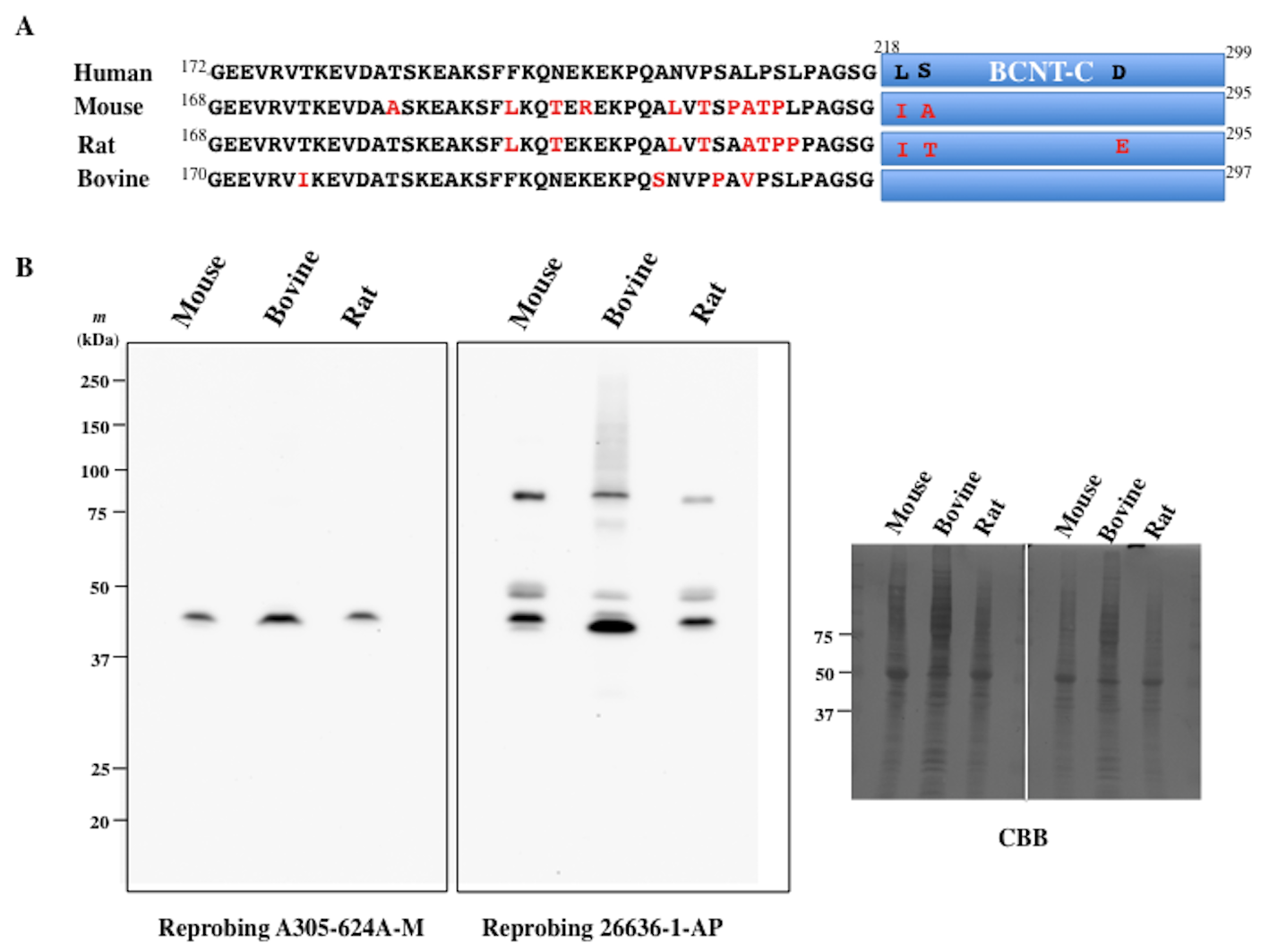
