## Supporting information on S1-S6 for "Assessment of a western blot signal for the Bcnt/Cfdp1, a tentative component of Srcap chromatin remodeling complex; trial to overcome off-target problems"

**S1 Fig. Screening of F-mBcnt expressing T-REx cell clones by western blot**.

Top panel: Molecular architecture of Flag-tagged mouse Bcnt/Cfdp1. The protein shown in a large outline comprises the acidic N-terminal region, Lys/Glu/Pro-rich 40 amino acids (named intramolecular repeat [IR], white box), and a highly conserved C-terminal region (BCNT-C domain, blue box) in addition to the Flag-tag at the N-terminus (yellow box). The numbers above the outline show the amino acid residues of mBcnt/Cfdp1. The underlined red bar presents the location of the immunogen for the generation of anti-BCNT-C Ab.

Cell extracts of several G418 resistant colonies from F-mBcnt transfectants and of MCS- or F-mBcnt transiently expressed cultures (all equivalent to ~ 5 x 10^4^ cells per lane) were subjected to western blot analysis wanti-Flag Tag Ab at 500 ng/mL (left panel) an-BCNT-C Ab at 1 μg/mL (right panel). C3* indicates a loss of the sample in applying to the gel (in left panel).

**S2 Fig. Isolation of F-mBcnt from T-REx clone E3 by anti-Flag Ab agarose bead.**

The supernatant of the E3 cell extract was mixed with anti-Fla-tag agarose beads and incubated. The bound fraction was washed, eluted with Flag (DYKDDDK) peptide sequentially (Eluate #1 and #2), and followed by glycine-HCl, pH 2.5 (Low pH). Finally, the agarose was boiled in SDS/PAGE sample buffer for 5 min (SDS). All of the samples were subjected to western blotting with anti-Flag Tag Ab (250 ng/mL) followed by HRP-conjugated anti-mouse IgG light-chain-specific Ab. Finally, the filter was stained with Coomassie Brilliant Blue (CBB, right panel).

**S3 Fig. Enrichment of bovine Bcnt/Cfdp1 by Phos-tag agarose.**

The supernatant fraction of the bovine placenta extract in RIPA buffer was mixed with Phos-tag agarose and incubated. After washing three times (Wash#1, 2, 3), the bound portion was sequentially eluted (Eluate #1, 2, 3), and each sample was concentrated with TCA. On the other hand, the pellet fraction, which contained 9-fold concentrated protein compared to the supernatant, was prepared (PPT). All of the samples were boiled in SDS/PAGE sample buffer and then subjected to western blotting analysis with A305-624-M at 1 μg/mL (left side) or anti-mBcntN Ab at 250 ng/mL (right side). Non-concentrated sample (NC) was also analyzed for reference. After obtaining their images, the filter that had been used with anti-mBcnt-N Ab was re-probed with A305-624-M at 1 μg/mL (rightmost upper panel). Finally, the filters were stained with Coomassie Brilliant Blue (CBB, bottom panel).

**S4 Fig. Confirmation of anti-mBcnt-N Ab specificity with A305-624-M or 26636-1-AP.**

(A) Alignment of amino acid sequences of Ab 26636-1-AP immunogens (172-299 of hBCNT/CFDP1 as shown at the top) among the counterparts of the mouse, rat, and bovine. Blue boxes indicate the BCNT-C domain. Red letters present the different amino acids corresponding to the human counterpart. It is noticed that bovine Bcnt/Cfdp1 is more similar to hBCNT/CFDP1 than mouse and rat counterparts, indicating a higher probability of immunoreaction of 26636-1-AP against the bovine than other counterparts. (B) Two filters used for evaluation of anti-mBcnt-N Ab specificity, as shown in Fig 5, were re-probed with A305-624-M at 1 μg/mL (left panel) or 26636-1-AP at 280 ng/mL (right panel) for confirmation of loading amounts of each immunoreactive Bcnt/Cfdp1. Finally, all filters were stained with Coomassie Brilliant Blue (CBB, bottom in right panel).

**S5 Fig. Identity of the ~45 kDa band detected with the two antibodies.**

Equal amounts of cell extract (20 μg protein) of vdR2-4 (WT) or Cfdp1-K1 (Mutant) cells serially passaged in the presence [Mutant (2)] or absence [Mutant (1)] of G418 and puromycin were subjected to western blot analysis with anti-mBcnt-N Ab (left panel). After stripping the filter was re-probed with A305-624A-M (middle panel). A red arrow indicates a signal of mouse Bcnt/Cfdp1. As described in the legends of Fig 6, a band detected with anti-mBcnt-N Ab at ~75 kDa (shown as *) is probably the fusion protein of a part of mBcnt (derived from exon 1-5) and hygromycin phosphotransferase. The filter was finally stained with Coomassie Brilliant Blue (CBB) to confirm the amounts of loading protein.

**S6 Fig. Isolation of Nono (p54rnb) by anti-His-tag Ab agarose.**

(A) Evaluation of anti-p54nrb Ab GTX101419 from Gene Tex. T-REx 293 Clone G11 cells were transfected cells with HA-P54nrb pcDNA3.1 (obtained from Dr. Atsushi Yokoyama). After 70 h, the cell lysate was prepared and centrifuged (25,000 x *g*, 30 min, 4 ℃). The protein ratio between fractions of the supernatant (SUP) and the particulate (PPT) was 8 : 1. Their constant amounts of protein (18 μg/lane) were subjected to western blot analysis with anti-p54nrb antibody (250 ng/mL). (B) Isolation of Nono by anti-His tag antibody-conjugated agarose beads. The supernatant of T-REx cell lysate was applied to anti-His-tag antibody-conjugated agarose beads, and after washing, the bound protein to agarose was incubated with His-tag peptide, followed by with glycine-HCl buffer (pH 2.5), and finally boiled in SDS/PAGE sample buffer. All of the samples were boiled in SDS/PAGE sample buffer and subjected to western blot analysis with anti-p54rnb antibody (250 ng/mL).

**Table titles**

**S１Appendix**

**Summary of whole transcriptome sequencing: different gene expression profiles.**

**S1 Table**

**Protease-digested fragments of the upper and the lower bands of F-mBcnt.**

**S2 Table**

**Whole expression profile in Cfdp-1 (Mutant) and vdR2-4 (Wild type) cells.**

**S3 Table**

**Differential expression profiles of *Bcnt/Cfdp1* flanking genes and internal control genes between Cfdp1-K1 and vdR2-4 cells.**

**S4 Table**

**Downregulation of *keratins* 8, 18, and 19 in Cfdp1-K1 cells.**

Other highlighted genes were listed based on reference [36].

**S5 Table**

**List of reagents and materials.**

**S6 Table**

**Oligonucleotide sequences for plasmid construction and mRNA examination.**
