## Supplemental for "Assessment of a western blot signal for the Bcnt/Cfdp1, a tentative component of Srcap chromatin remodeling complex; trial to overcome off-target problems"

Research use only

### *Mus musculus*
### Transcriptome Sequencing

### Report

March 2019

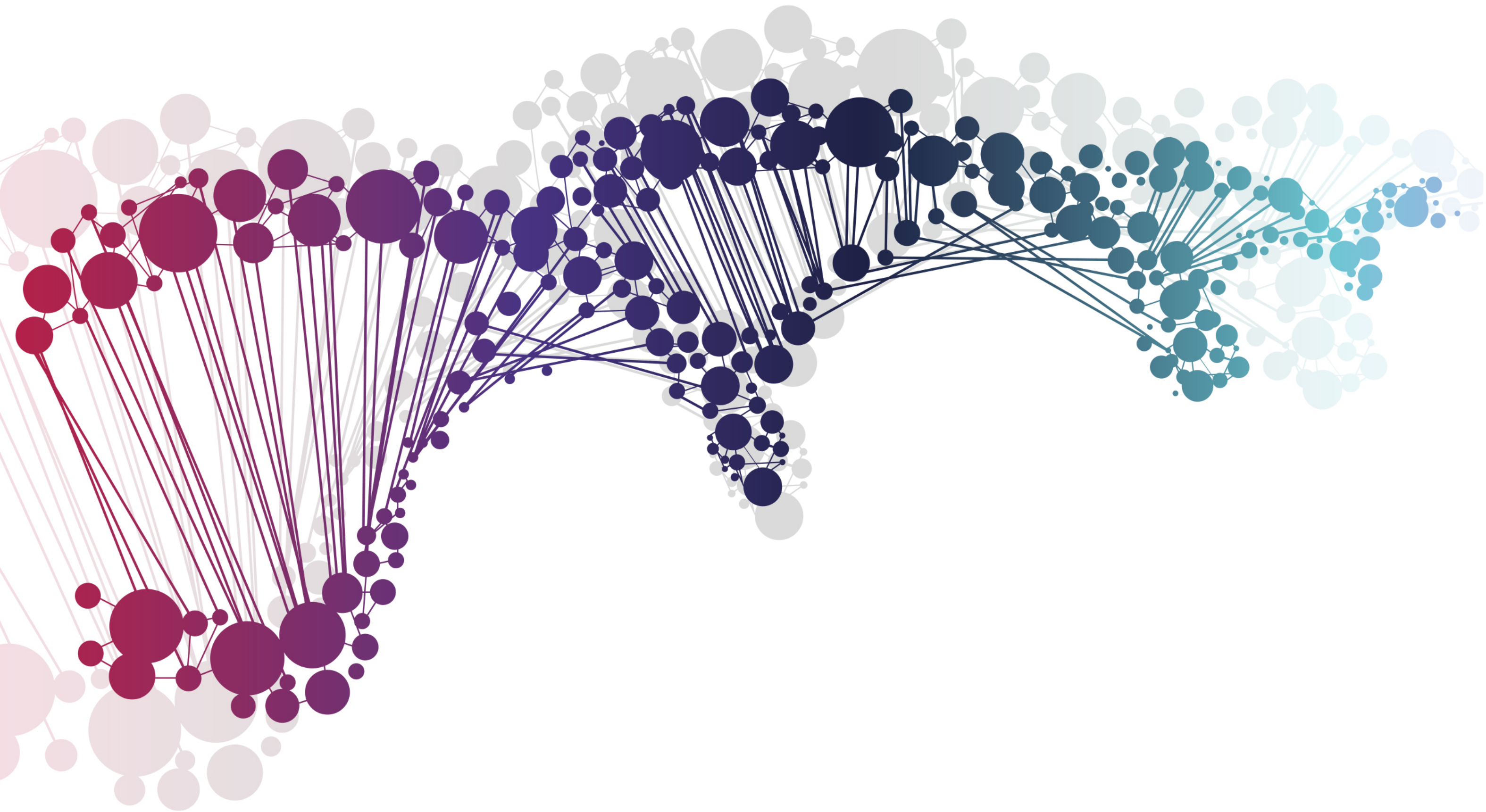

Humanizing Genomics
macrogen

ver. MGTR3.4_HS_mm10

COPYRIGHT © MACROGEN INC.

### Project Information

| | |
|---|---|
| Client Name | Macrogen Japan |
| Company/Institution | Macrogen Corp. Japan |
| Order Number | HN00101712 |
| Species | *Mus musculus* |
| Reference | UCSC mm10 |
| Annotation | RefSeq_2017_06_12 |
| Read Length | 101 |
| Number of Samples | 2 |
| Library Kit | TruSeq Stranded mRNA LT Sample Prep Kit |
| Library Protocol | TruSeq Stranded mRNA Sample Preparation Guide, Part # 15031047 Rev. E |
| Reagent | NovaSeq 6000 S4 Reagent Kit |
| Sequencing Protocol | NovaSeq 6000 System User Guide Document # 1000000019358 v02 |
| Type of Sequencer | NovaSeq |
| Sequencing Control Software | 1000000019358 v02 |

### Project Results Summary

In this study, *Mus musculus* whole transcriptome sequencing was performed in order to examine the different gene expression profiles, and to perform gene annotation on set of useful genes based on gene ontology pathway information.

Analyses were successfully performed on all 2 paired-ends samples. Figure 1 shows the throughput of raw data and trimmed data. Figure 2 shows the Q30 percentage (% of bases with quality over phred score 30) of each sample’s raw and trimmed data.

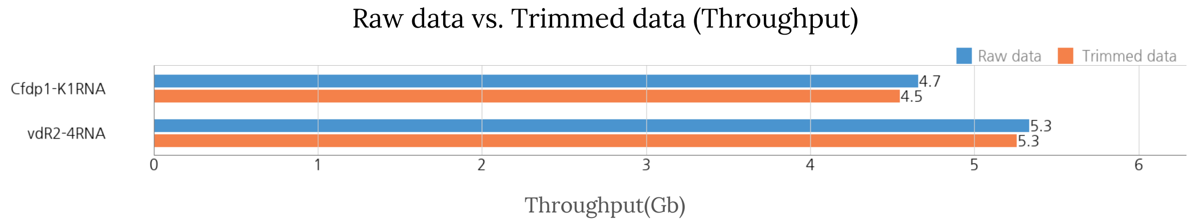


Figure 1. Throughput output of Raw and Trimmed data

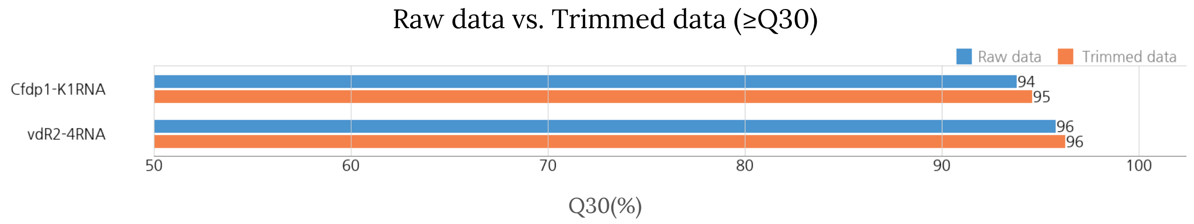


Figure 2. Q30 score of Raw and Trimmed data

Trimmed reads are mapped to reference genome with HISAT2. Figure 3 shows the overall read mapping ratio, the ratio of mapped reads to trimmed reads.

Overall read mapping ratio(%)

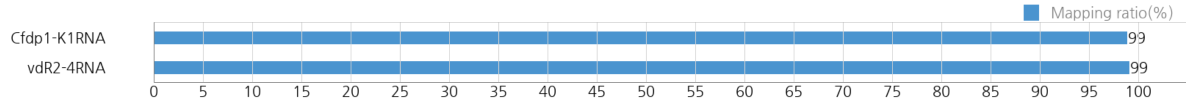


Figure 3. Overall read mapping ratio(%)

After the read mapping, Stringtie was used for transcript assembly. Expression profile was calculated for each sample and transcript/gene as read count and FPKM (Fragment per Kilobase of transcript per Million mapped reads).

DEG (Differentially Expressed Genes) analysis was performed on a comparison pair (Cfdp1-K1RNA_vs_vdR2-4RNA) as requested using FPKM. The results showed 694 genes which satisfied |fc|>=2 conditions in comparison pair.

Figure 4 shows the result of hierarchical clustering (distance metric= Euclidean distance, linkage method= complete) analysis. It graphically represents the similarity of expression patterns between samples and genes.

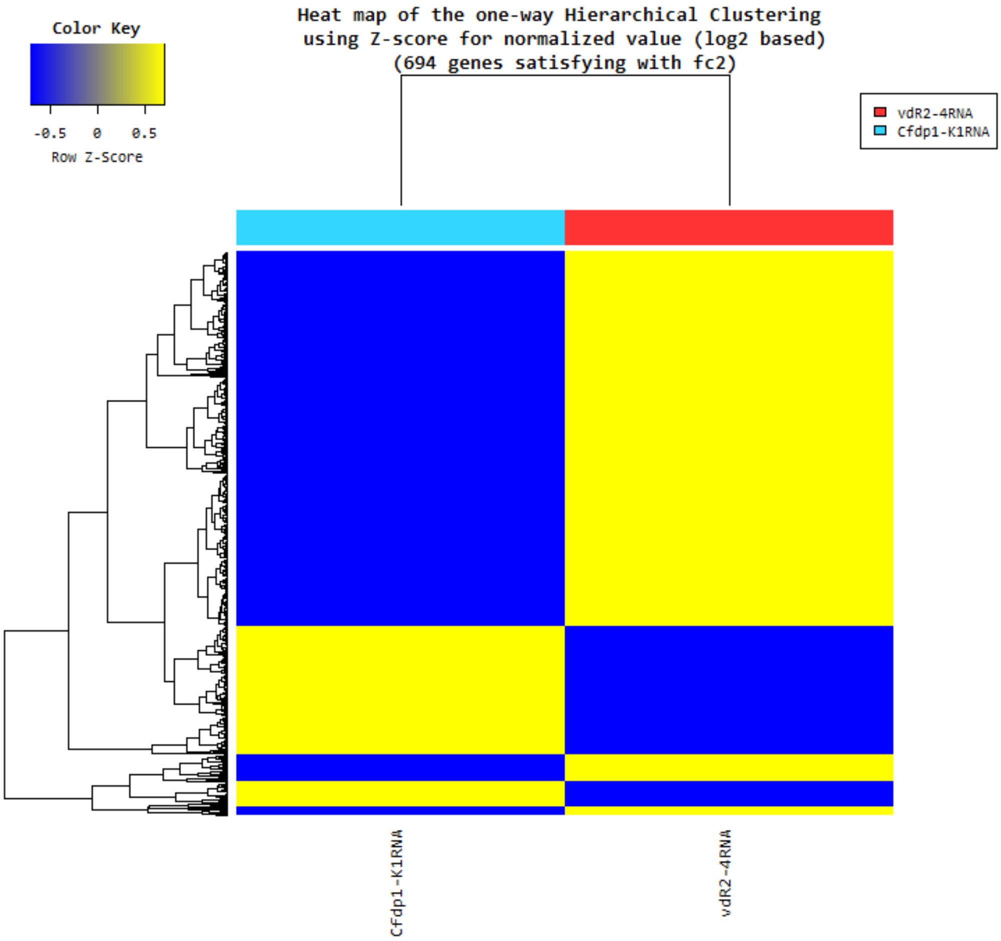


Figure 4. Heatmap for DEG list

DEG list was further analyzed with Gene Ontology (http://geneontology.org/) for gene set enrichment analysis per biological process (BP), cellular component (CC) and molecular function (MF). The Figure 5, 6 and 7 show the significant gene set by each category.

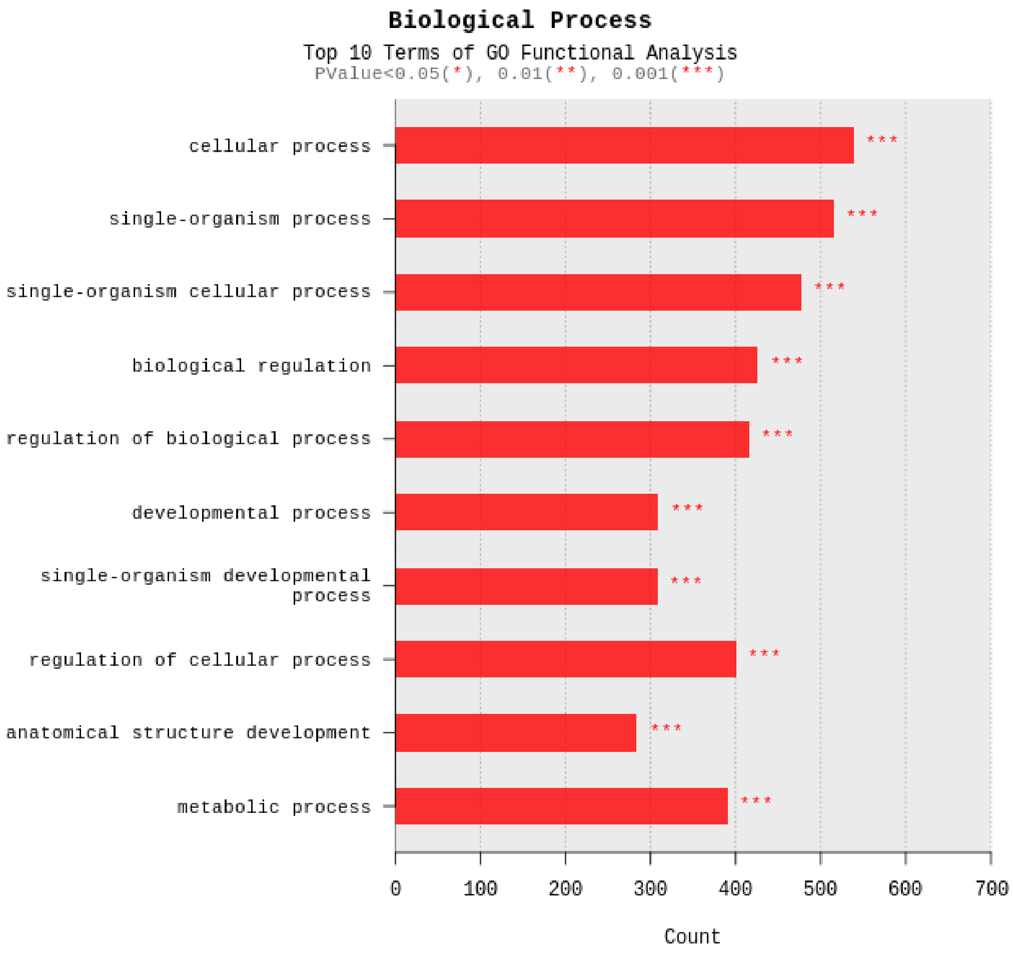


Figure 5. Gene Ontology terms related to Biological Process

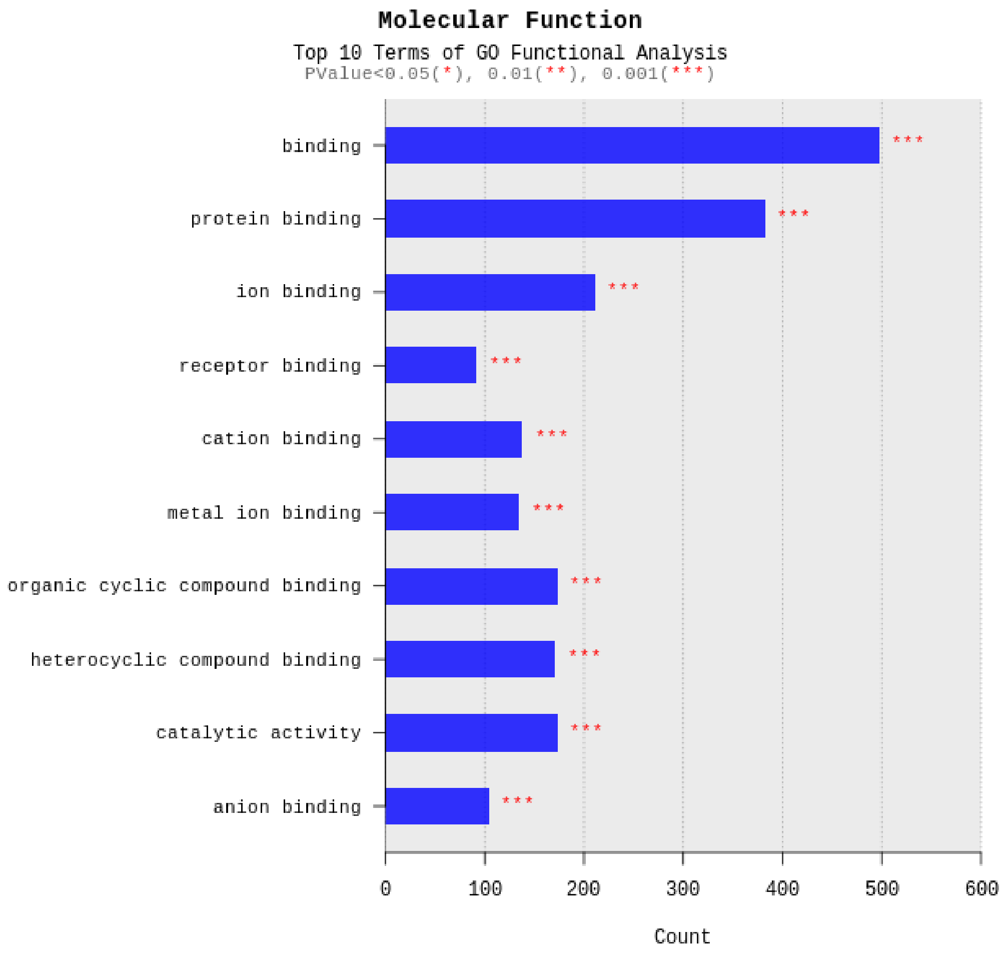


Figure 6. Gene Ontology Terms related to Molecular Function

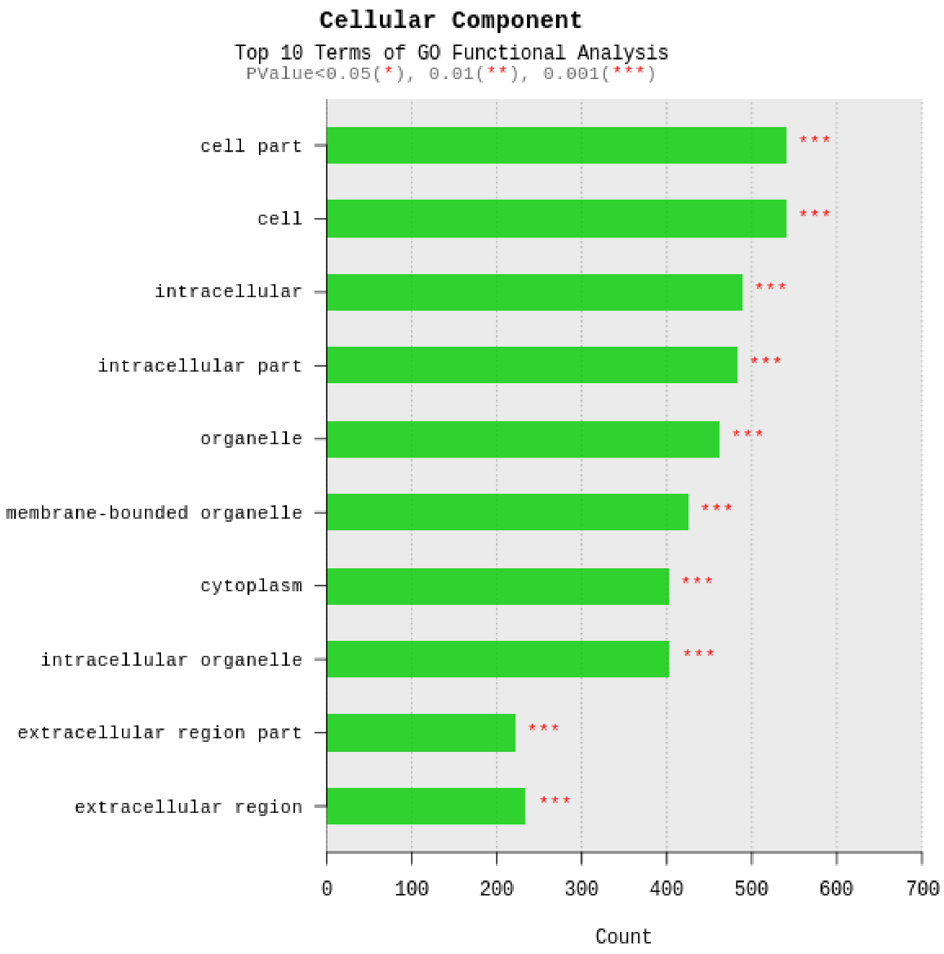


Figure 7. Gene Ontology Terms related to Cellular Component

### Table of Contents


Project Information 02

Project Results Summary 03

1. Experimental Methods and Workflow 10

2. Analysis Methods and Workflow 11

3. Summary of Data Production 12
   - 3. 1. Raw Data Statistics 12
   - 3. 2. Average Base Quality at Each Cycle 13
   - 3. 3. Trimming Data Statistics 14
   - 3. 4. Average Base Quality at Each Cycle after Trimming 15

4. Reference Mapping and Assembly Results 16
   - 4. 1. Mapping Data Statistics 16
   - 4. 2. Expression Profiling 17

5. Differentially Expressed Gene Analysis Results 19
   - 5. 1. Data Analysis Quality Check and Preprocessing 19
   - 5. 2. Differentially Expressed Gene Analysis Workflow 24
   - 5. 3. Significant Gene Results 25
   - 5. 4. GO Enrichment Analysis 30

6. Data Download Information 36
   - 6. 1. Raw Data 36
   - 6. 2. Analysis Results 36

7. Appendix 39
   - 7. 1. Phred Quality Score Chart 39
   - 7. 2. Programs used in Analysis 40
   - 7. 3. References 41

### 1. Experimental Methods and Workflow

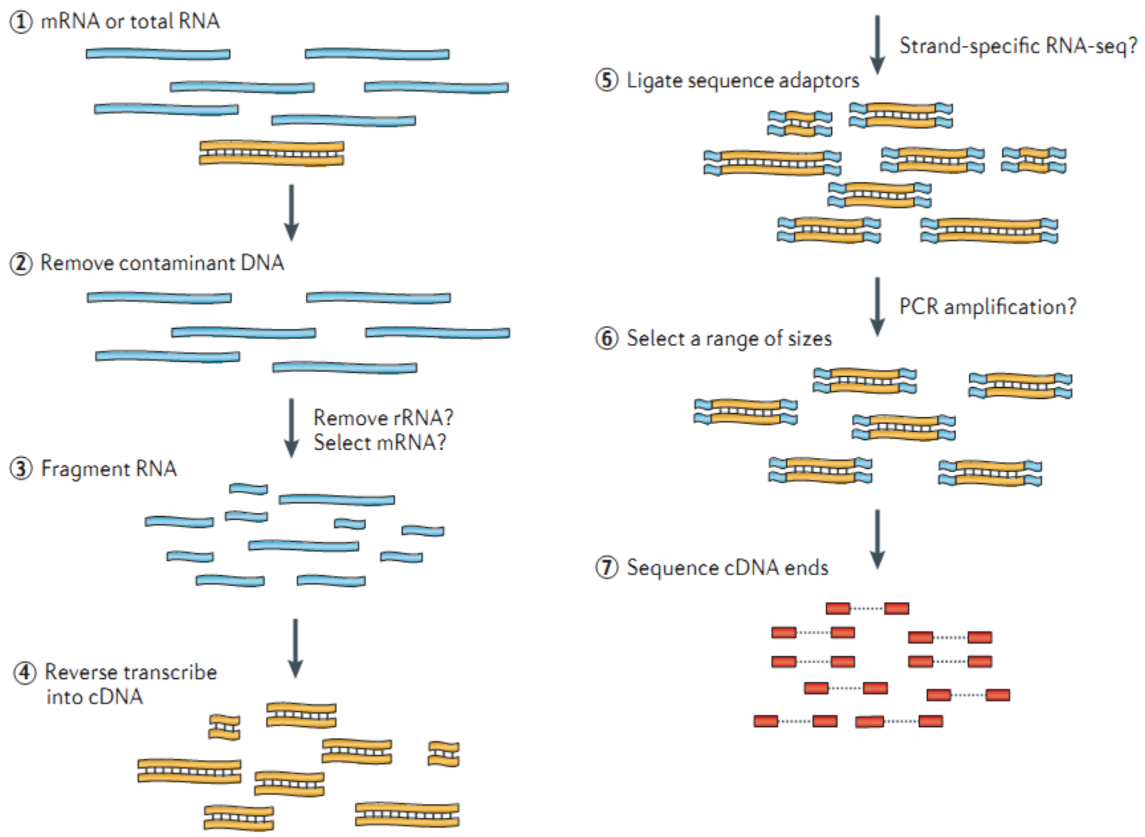


Figure 8. RNA Sequencing Experiment Workflow

REFERENCE Nat Rev Genet. 2011 Sep 7;12(10):671-82

1) Isolate the Total RNA from Sample of interest (Cell or Tissue).

2) Eliminate DNA contamination using DNase.

3) Choose an appropriate kit for library prep process depending on the types of RNA. For mRNA with poly-A tail, use mRNA purification kit; for non-coding RNAs, such as lincRNA, use ribo-zero RNA removal Kit to purify RNA of interest.

4) Randomly fragment purified RNA for short read sequencing.

5) Reverse transcribe fragmented RNA into cDNA.

6) Ligate adapters onto both ends of the cDNA fragments.

7) After amplifying fragments using PCR, select fragments with insert sizes between 200-400 bp. For paired-end sequencing, both ends of the cDNA is sequenced by the read length.

#### 2. Analysis Methods and Workflow

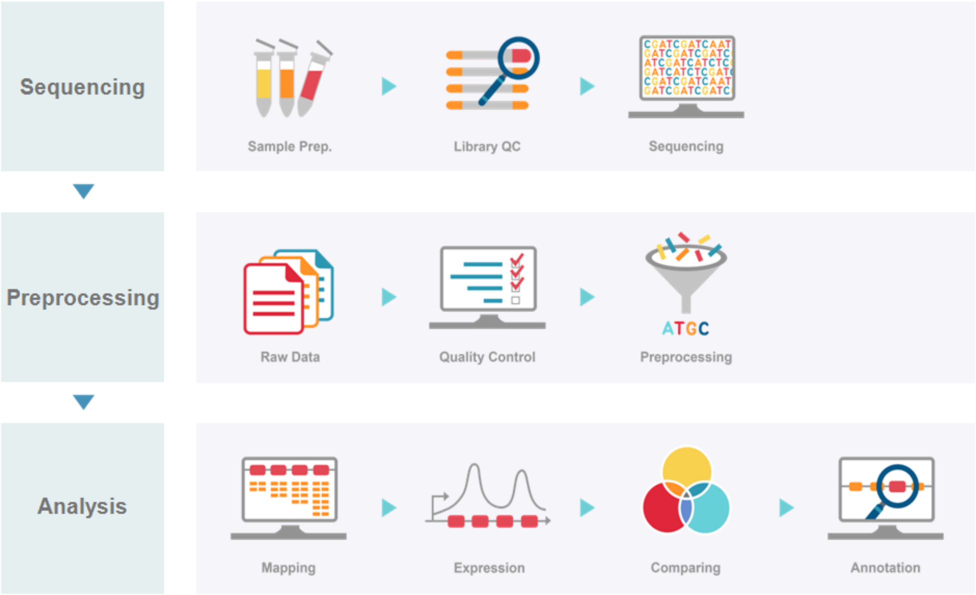


Figure 9. Analysis Workflow

1) Analyze the quality control of the sequenced raw reads. Overall reads' quality, total bases, total reads, GC (%) and basic statistics are calculated.

2) In order to reduce biases in analysis, artifacts such as low quality reads, adaptor sequence, contaminant DNA, or PCR duplicates are removed.

3) Trimmed reads are mapped to reference genome with HISAT2, splice-aware aligner.

4) Transcript is assembled by StringTie with aligned reads.

5) Expression profiles are represented as read count and normalization value which is based on transcript length and depth of coverage. The FPKM (Fragments Per Kilobase of transcript per Million Mapped reads) value or the RPKM (Reads Per Kilobase of transcript per Million mapped reads) is used as a normalization value.

6) In groups with different conditions, genes or transcripts that express differentially are filtered out through statistical hypothesis testing.

7) In case of known gene annotation, functional annotation and gene-set enrichment analysis are performed using GO and KEGG database on differentially expressed genes.

### 3. Summary of Data Production

#### 3. 1. Raw Data Statistics

(Refer to Path: result_RNAseq/Analysis_statistics/rawData/raw_throughput.stats)

The total number of bases, reads, GC (%), Q20 (%), Q30 (%) are calculated for 2 samples. For example, in Cfdp1-K1RNA, 46,100,372 reads are produced, and total read bases are 4.7Gbp. The GC content (%) is 49.57% and Q30 is 93.77%.

Table 1. Raw data stats

| Index | Sample id | Total read bases* | Total reads | GC (%) | Q20 (%) | Q30 (%) |
|---|---|---|---|---|---|---|
| 1 | Cfdp1-K1RNA | 4,656,137,572 | 46,100,372 | 49.57 | 97.87 | 93.77 |
| 2 | vdR2-4RNA | 5,331,793,636 | 52,790,036 | 50.52 | 98.66 | 95.75 |

(* Total read bases = Total reads x Read length)

- Total read bases: Total number of bases sequenced
- Total reads: Total number of reads
- GC (%): GC content
- Q20 (%): Ratio of bases that have phred quality score greater than or equal to 20
- Q30 (%): Ratio of bases that have phred quality score greater than or equal to 30

#### 3. 2. Average Base Quality at Each Cycle

(Refer to Path: Analysis_statistics/rawData/A_fastqc/)

The quality of produced data is determined by the phred quality score at each cycle. Box plot containing the average quality at each cycle is created with FastQC.

The x-axis shows number of cycles and y-axis shows phred quality score. Phred quality score 20 means 99% accuracy and reads over score of 20 are accepted as good quality.

LINK http://www.bioinformatics.babraham.ac.uk/projects/fastqc

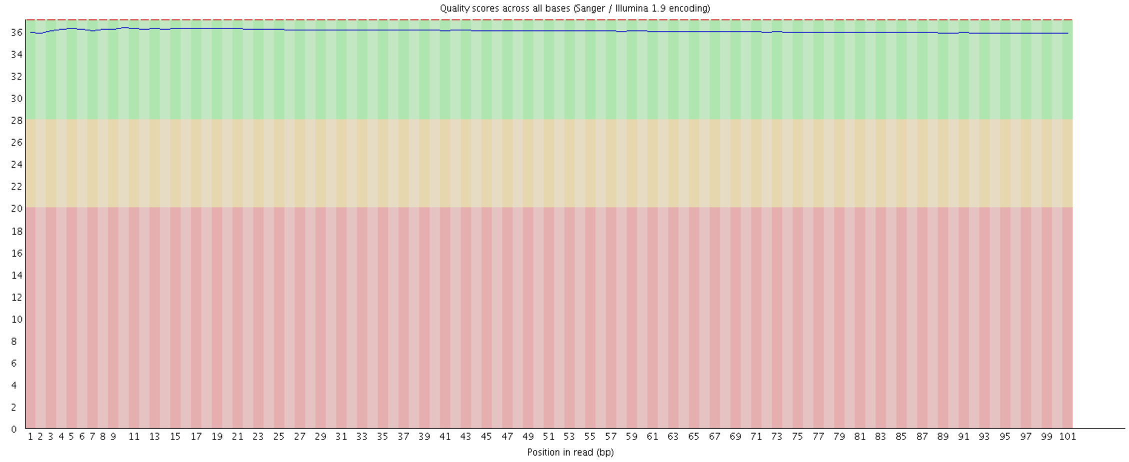


Figure 10. Read quality at each cycle of Cfdp1-K1RNA (read1)

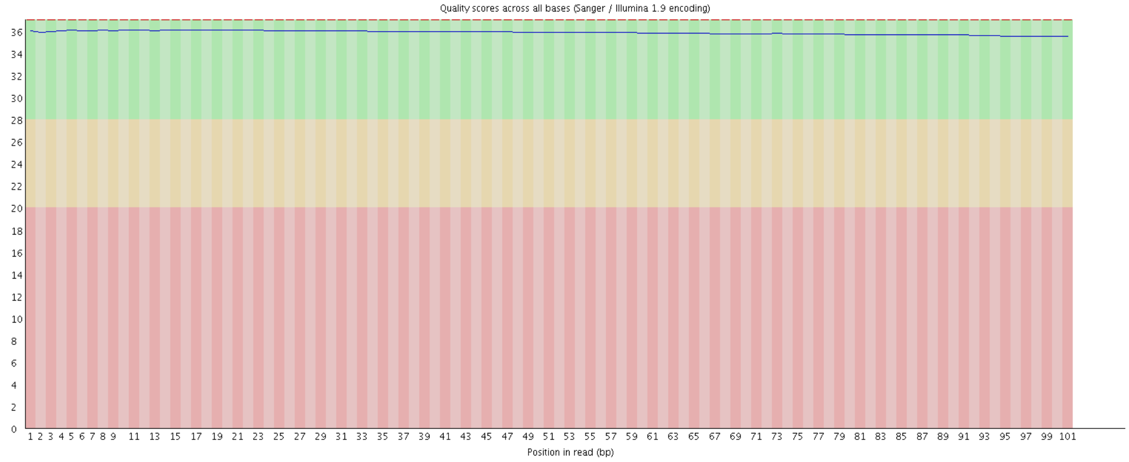


Figure 11. Read quality at each cycle of Cfdp1-K1RNA (read2)

- Yellow box: Interquartile range (25-75%) of phred score at each cycle
- Red line: Median of phred score at each cycle
- Blue line: Average of phred score at each cycle
- Green background: Good quality
- Orange background: Acceptable quality
- Red background: Bad quality

### 3. 3. Trimming Data Statistics

(Refer to Path: result_RNAseq/Analysis_statistics/trimmedData/trim_throughput.stats)

Trimmomatic program is used to remove adapter sequences and bases with base quality lower than three from the ends. Also using sliding window method, bases of reads that does not qualify for window size 4, and mean quality 15 are trimmed. Afterwards, reads with length shorter than 36bp are dropped to produce trimmed data.

Table 2. Trimming Data Stats

| Index | Sample id | Total read bases | Total reads | GC(%) | Q20(%) | Q30(%) |
|---|---|---|---|---|---|---|
| 1 | Cfdp1-K1RNA | 4,543,927,517 | 45,262,214 | 49.59 | 98.39 | 94.57 |
| 2 | vdR2-4RNA | 5,253,493,756 | 52,245,252 | 50.53 | 98.99 | 96.25 |

- Total read bases: Total number of read bases after trimming
- Total reads: Total number of reads after trimming
- GC (%): GC Content
- Q20 (%): Ratio of bases that have phred quality score greater than or equal to 20
- Q30 (%): Ratio of bases that have phred quality score greater than or equal to 30

#### 3. 4. Average Base Quality at Each Cycle after Trimming

(Refer to Path: result_RNAseq/Analysis_statistics/trimmedData/A_fastqc/)

Figure 12 and 13 show average base quality at each cycle after trimming.

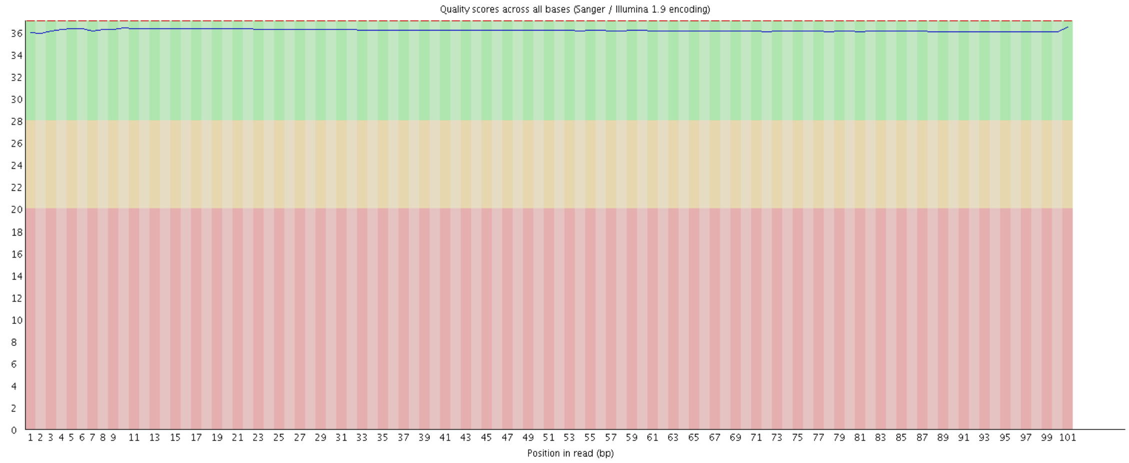


Figure 12. Average base quality of Cfdp1-K1RNA (read1) at each cycle after trimming

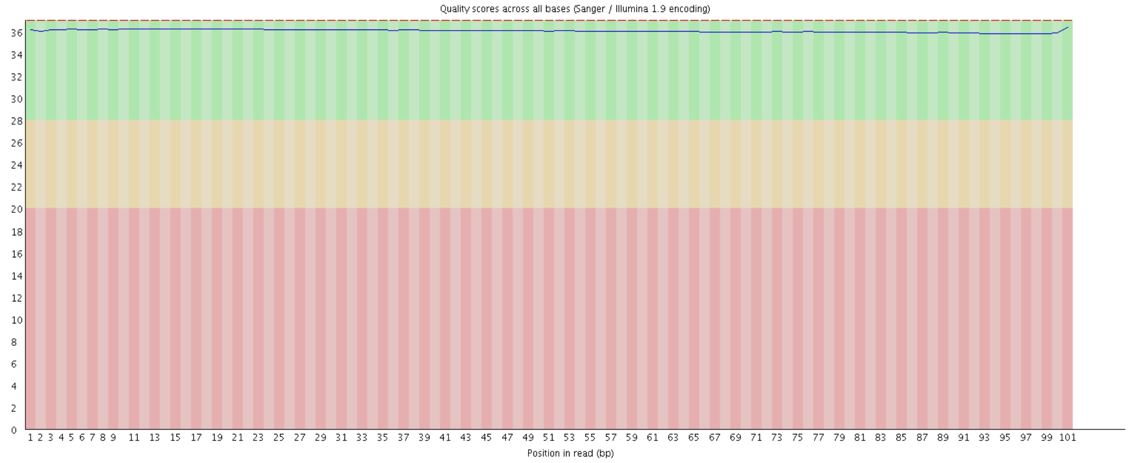


Figure 13. Average base quality of Cfdp1-K1RNA (read2) at each cycle after trimming

- Yellow box: Interquartile range (25-75%) of phred score at each cycle
- Red line: Median of phred score at each cycle
- Blue line: Average of phred score at each cycle
- Green background: Good quality
- Orange background: Acceptable quality
- Red background: Bad quality

### 4. Reference Mapping and Assembly Results

#### 4. 1. Mapping Data Statistics

(Refer to Path: result_RNAseq/Analysis_statistics/mapping.hisat.stats)

In order to map cDNA fragments obtained from RNA sequencing, UCSC mm10 was used as a reference genome. Table 3 shows the statistic obtained from HISAT2, which is known to handle spliced read mapping through Bowtie2 aligner. You can check number of processed reads, mapped reads.

Table 3. Mapped Data Stats

| Sample ID | # of processed reads | # of mapped reads (%) | # of unmapped reads (%) |
|---|---|---|---|
| Cfdp1-K1RNA | 45,262,214 | 44,725,812 (98.81%) | 536,402 (1.19%) |
| vdR2-4RNA | 52,245,252 | 51,767,696 (99.09%) | 477,556 (0.91%) |

- Processed reads: Number of cleaned reads after trimming
- Mapped reads: Number of reads mapped to reference
- Unmapped reads: Number of reads that failed to align

### 4. 2. Expression Profiling

Known genes and transcripts are assembled with StringTie based on reference genome model. After assembly, the abundance of gene/transcript is calculated in the read count and normalized value as FPKM (Fragments Per Kilobase of transcript per Million mapped reads) for a sample.

#### 4. 2. 1. Known Transcripts Expression Level

(Refer to Path: result_RNAseq_excel/Expression_profile/StringTie/ Expression_Profile.mm10.transcript.xlsx)

Table 4 is an example of known transcript expression level per sample in expression value. This result is obtained by -e option of StringTie does not consider novel transcript assembly.

Table 4. Known transcripts Expression Level (example)

| Transcript_ID | Gene_ID | Gene_Symbol | Description | Transcript_Locus | Trancript_Length | AM Read_Count | BM Read_Count | AM_FPKM | BM_FPKM |
|---|---|---|---|---|---|---|---|---|---|
| NM_001302545 | 14 | AAMP | angio associated migratory cell protei | chr2:219128852-219134 | 1835 | 898 | 987 | 12.220251 | 12.415353 |
| NM_001087 | 14 | AAMP | angio associated migratory cell protei | chr2:219128852-219134 | 1832 | 4678 | 6437 | 63.774269 | 81.140015 |
| NM_001166579 | 15 | AANAT | aralkylamine N-acetyltransferase, trar | chr17:74449433-744661 | 1913 | 46 | 30 | 0.599741 | 0.352587 |
| NR_110548 | 15 | AANAT | aralkylamine N-acetyltransferase, trar | chr17:74463630-744661 | 1082 | 9 | 9 | 0.192813 | 0.186779 |
| NM_001101 | 60 | ACTB | actin beta | chr7:5566779-5570232 | 1812 | 93591 | 129901 | 1290.007935 | 1655.640503 |
| NM_001161572 | 23764 | MAFF | MAF bZIP transcription factor F, trans | chr22:38597939-386125 | 2465 | 1 | 150 | 0.002107 | 1.397431 |
| NM_012323 | 23764 | MAFF | MAF bZIP transcription factor F, trans | chr22:38597939-386125 | 2439 | 1682 | 2109 | 17.222849 | 19.96483 |
| NM_001161574 | 23764 | MAFF | MAF bZIP transcription factor F, trans | chr22:38597939-386125 | 2372 | 0 | 0 | 0 | 0 |
| NM_001161573 | 23764 | MAFF | MAF bZIP transcription factor F, trans | chr22:38599027-386125 | 2223 | 44 | 25 | 0.485203 | 0.252227 |
| NM_001289905 | 23765 | IL17RA | interleukin 17 receptor A, transcript va | chr22:17565849-175965 | 8506 | 1303 | 975 | 3.825815 | 2.644646 |
| NM_014339 | 23765 | IL17RA | interleukin 17 receptor A, transcript va | chr22:17565849-175965 | 8608 | 3241 | 1998 | 9.402107 | 5.359576 |
| NR_028287 | 23766 | GABARAPL3 | GABA type A receptor associated pro | chr15:90889763-908926 | 1885 | 3 | 6 | 0.036076 | 0.073511 |
| NM_001017526 | 23779 | ARHGAP8 | Rho GTPase activating protein 8, trar | chr22:45148438-452586 | 1725 | 460 | 641 | 6.645803 | 8.576918 |
| NM_181335 | 23779 | ARHGAP8 | Rho GTPase activating protein 8, trar | chr22:45148438-452586 | 1632 | 1979 | 2405 | 30.27355 | 34.027134 |
| NM_001198726 | 23779 | ARHGAP8 | Rho GTPase activating protein 8, trar | chr22:45148438-452586 | 1528 | 84 | 59 | 1.366953 | 0.889118 |
| NM_030882 | 23780 | APOL2 | apolipoprotein L2, transcript variant al | chr22:36622255-366356 | 2545 | 559 | 1155 | 5.482551 | 10.474212 |
| NM_145637 | 23780 | APOL2 | apolipoprotein L2, transcript variant b | chr22:36622255-366360 | 2686 | 1212 | 0 | 11.260728 | 0 |

- Transcript_ID: Splicing variant (isoform/transcript)
- Gene_ID: Gene ID
- Gene_Symbol: Symbol of gene
- Gene_Description: Description of gene
- Transcript_Locus: Transript locus
- Transcript_Length: Transcript length
- [Sample Name]_Read_Count: Read count of a sample
- [Sample Name]_FPKM: FPKM normalized value of a sample

### 4. 2. 2. Known Genes Expression Level

(Refer to Path: result_RNAseq_excel/Expression_profile/StringTie/Expression_Profile.mm10.gene.xlsx)

Table 5 is an example of known gene expression level per sample in expression value. This result is obtained by -e option of StringTie does not consider novel transcript assembly.

Table 5. Known genes Expression Level (example)

| Gene_ID | Transcript_ID | Gene_Symbol | Description | AM Read_Count | BM Read_Count | AM_FPKM | BM_FPKM |
|---|---|---|---|---|---|---|---|
| 60 | NM_001101 | ACTB | actin beta | 93591 | 129901 | 1290.007935 | 1655.640503 |
| 70 | NM_005159 | ACTC1 | actin, alpha, cardiac muscle 1 | 20 | 6 | 0.1339 | 0.031949 |
| 175 | NM_000027,NM_001171988,NR_0 | AGA | aspartylglucosaminidase | 252 | 279 | 2.995219 | 3.071083 |
| 176 | NM_001135,NM_013227 | ACAN | aggrecan | 8 | 0 | 0.022519 | 0 |
| 177 | NM_001136,NM_001206929,NM_0 | AGER | advanced glycosylation end-product specif | 3332 | 3124 | 51.224842 | 44.355004 |
| 178 | NM_000028,NM_000642,NM_0006 | AGL | amylo-alpha-1, 6-glucosidase, 4-alpha-gluc | 4919 | 3679 | 16.662192 | 11.52329 |
| 191 | NM_000687,NM_001161766,NM_0 | AHCY | adenosylhomocysteinase | 12053 | 13891 | 129.59984 | 138.005572 |
| 245 | NR_002710,NR_120453 | ALOX12P2 | arachidonate 12-lipoxygenase pseudogene | 8 | 5 | 0.070872 | 0.041258 |
| 246 | NM_001140 | ALOX15 | arachidonate 15-lipoxygenase | 785 | 710 | 7.302354 | 6.108678 |
| 247 | NM_001039130,NM_001039131,NI | ALOX15B | arachidonate 15-lipoxygenase, type B | 6 | 0 | 0.049592 | 0 |
| 248 | NM_001631 | ALPI | alkaline phosphatase, intestinal | 13 | 3 | 0.098671 | 0.021092 |
| 249 | NM_000478,NM_001127501,NM_0 | ALPL | alkaline phosphatase, liver/bone/kidney | 9 | 19 | 0.085416 | 0.164094 |
| 250 | NM_001632 | ALPP | alkaline phosphatase, placental | 464 | 142 | 3.894943 | 1.098701 |
| 251 | NM_031313 | ALPPL2 | alkaline phosphatase, placental like 2 | 88 | 12 | 0.876858 | 0.106491 |
| 257 | NM_006492 | ALX3 | ALX homeobox 3 | 310 | 319 | 5.229297 | 4.975804 |
| 258 | NM_016519 | AMBN | ameloblastin | 0 | 0 | 0 | 0 |
| 259 | NM_001633 | AMBP | alpha-1-microglobulin/bikunin precursor | 0 | 0 | 0 | 0 |

- Gene_ID: Gene ID
- Transcript_ID: Splicing variant (isoform/transcript)
- Gene_Symbol: Symbol of gene
- Gene_Description: Description of gene
- [Sample Name]_Read_Count: Read count of a sample
- [Sample Name]_FPKM: FPKM normalized value of a sample

### 5. Differentially Expressed Gene Analysis Results

#### 5. 1. Data Analysis Quality Check and Preprocessing

There is a process that sorts differentially expressed gene among samples by FPKM value of known genes. In preprocessing, there are data quality and similarity checks among samples in case of biological replicates exist.

(Refer to Path: result_RNAseq_excel/DEG_result/Analysis_Result.html)

##### 5. 1. 1. Sample Information and Analysis Design

Total of 2 samples was used for analysis. For more information of samples and comparison pair, please refer to Sample.Info.txt file.

| Index | Sample.ID | Sample.Group |
|---|---|---|
| 1 | vdR2-4RNA | vdR2-4RNA |
| 2 | Cfdp1-K1RNA | Cfdp1-K1RNA |

Comparison pair and statistical method for each pair are shown below.

| Index | Test vs. Control | Statistical Method |
|---|---|---|
| 1 | Cfdp1-K1RNA vs. vdR2-4RNA | Fold Change, Hierarchical Clustering |

#### 5. 1. 2. DATA Quality Check

(Refer to Path: result_RNAseq_excel/DEG_result/Data Quality Check/)

For 2 samples, if more than one FPKM value was 0, it was not included in the analysis. Therefore, from total of 24,532 genes, 9,558 were excluded and only 14,974 genes were used for statistic analysis.

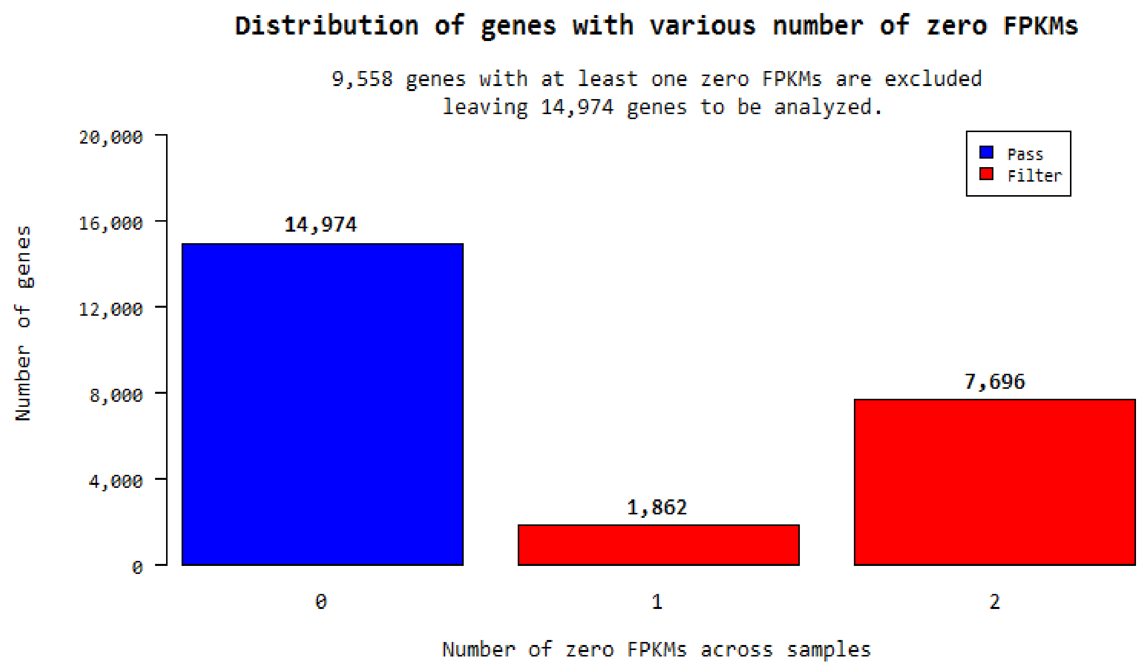


#### 5. 1. 3. Data Transformation and Normalization

To facilitate log2 transformation, 1 was added to the raw signal (FPKM). This process is performed because raw signals are scattered along wide range and most signals are concentrated on the low signal value, so log transformation reduces the range of the signals and produces more even data distribution. After log transformation, in order to reduce systematic bias, quantile normalization is used with preprocessCore' R library.

##### 5. 1. 3. 1. Boxplot of Expression Difference between samples.

Below boxplots show the corresponding sample's expression distribution based on percentile (median, 50 percentile, 75 percentile, maximum and minimum) based on raw signal (FPKM), Log2 transformation of FPKM+1 and Quantile Normalization.

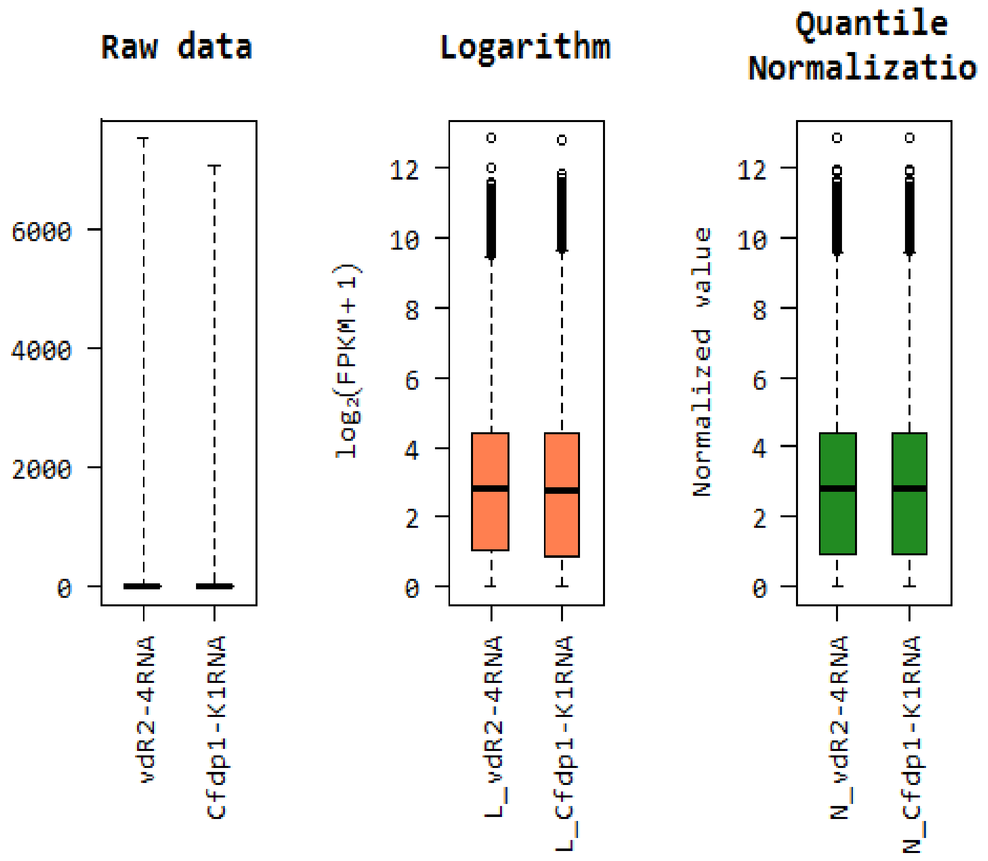

#### 5. 1. 3. 2. Expression Density Plot per sample

Below density plots show the corresponding samples expression distribution before and after of raw signal (FPKM), Log2 transformation of FPKM+1 and Quantile Normalization.

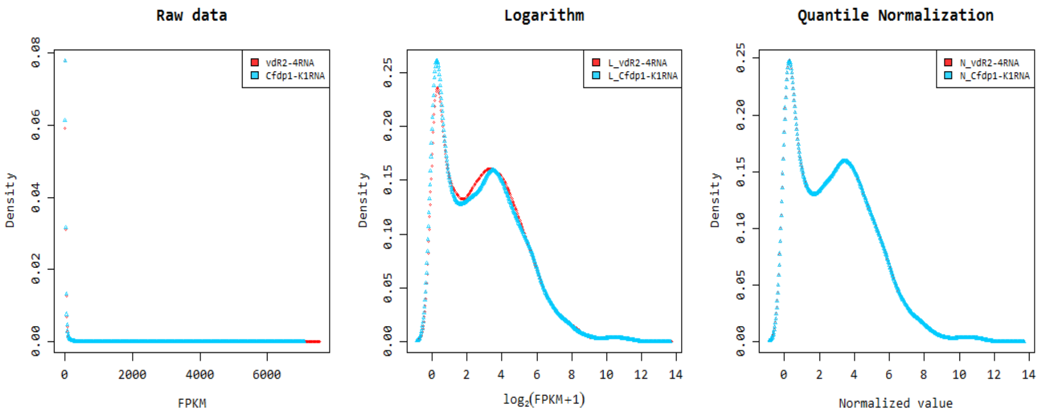

#### 5. 1. 4. Correlation Analysis between samples

The similarity between samples are obtained through Pearson's coefficient of the normalized value. For range: -1≤ r ≤ 1,the closer the value is to 1, the more similar the samples are.

Correlation matrix of all samples is as follows.

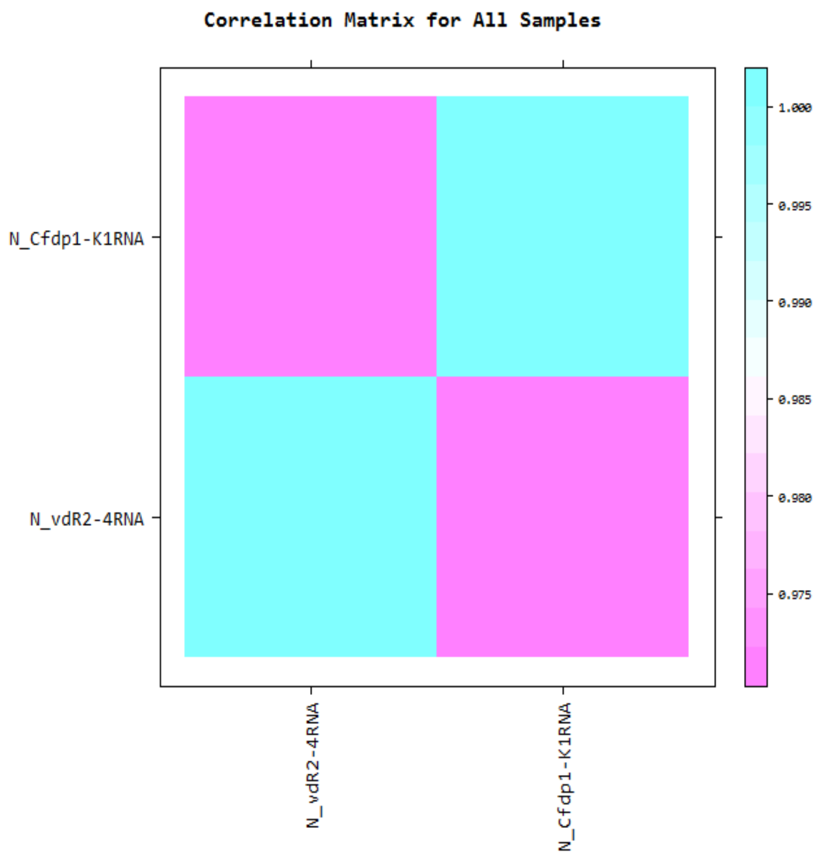

### 5. 2. Differentially Expressed Gene Analysis Workflow

Below shows the orders of DEG (Differentially Expressed Genes) analysis.

1) the FPKM value of known genes obtained through -e option of the StringTie were used as the original raw data.

- Raw data

(Refer to Path: result_RNAseq_excel/Expression_profile/StringTie/Expression_Profile.mm10.gene.xlsx)

: 24,532 genes, 2 samples

2) During data preprocessing, low quality transcripts are filtered. Afterwards, log2 transformation of FPKM+1 and quantile normalization are performed.

- Processed data

(Refer to Path: result_RNAseq_excel/DEG_result/data2.xlsx)

: 14,974 genes, 2 samples

3) Statistical analysis is performed using Fold Change per comparison pair. The significant results are selected on conditions of |fc|>=2.

- Significant data

(Refer to Path: result_RNAseq_excel/DEG_result/data3_fc2.xlsx)

: 694 genes

4) For significant lists, hierarchical clustering analysis is performed to group the similar samples and genes. These results are graphically depicted using heatmap and dendogram.

- Hierarchical Clustering (Euclidean Distance, Complete Linkage)

(Refer to Path: result_RNAseq_excel/DEG_result/Cluster image/)

5) For significant lists, gene-set enrichment analysis was performed based on gene ontology( http://geneontology.org/).

Please refer to the gomap_stat sheet and the gomap_genes sheet of data3 file.

Following result are provided.

- gomap_stat
- gomap_genes

#### 5. 3. Significant Gene Results

(Refer to Path: result_RNAseq_excel/DEG_result/Plots/)

These are DEG result of Cfdp1-K1RNA_vs_vdR2-4RNA meeting fc2 by example.

##### 5. 3. 1. Up, Down Regulated Count by Fold Change

Shows number of up and down regulated genes based on fold change of comparison pair.

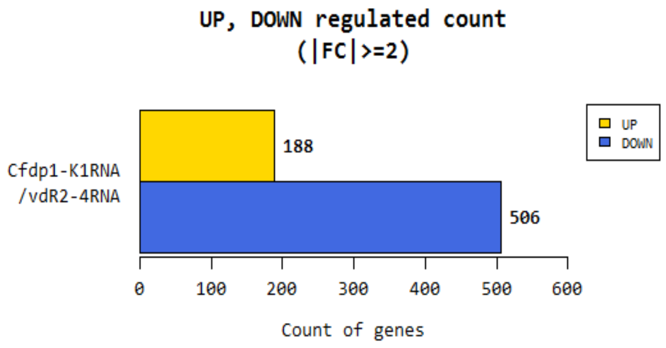

#### 5. 3. 2. Distribution of Expression Level between two groups

Shows distribution of normalized value of each group for comparison pair.

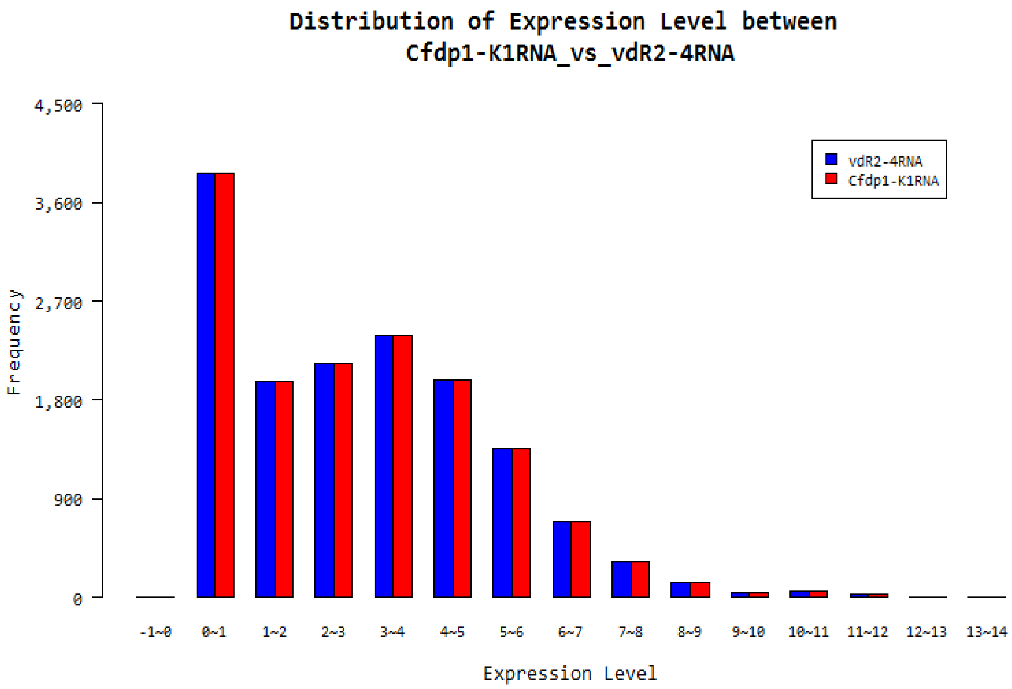

#### 5. 3. 3. Scatter Plot of Expression Level between two groups

Shows expression levels between comparison pair as a scatter plot. X-axis is control and Y-axis is average normalized value of the group.

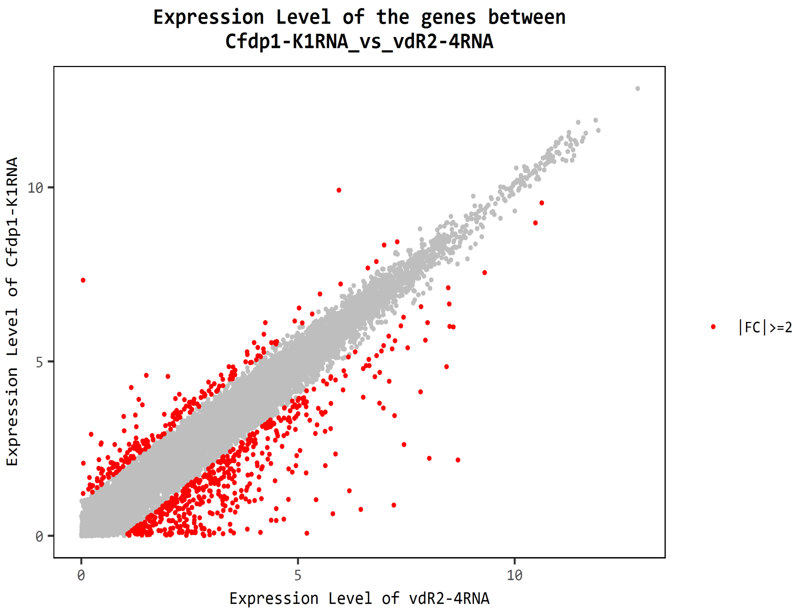

### 5. 3. 4. Volume Plot

Expression volume is defined as the geometric mean of two group's expression level. In order to confirm the genes that show higher expression difference compared to the control according to expression volume, volume plot is drawn. (X-axis: Volume, Y-axis: log2 Fold Change).

For example, even though fold change might be different by two-fold, the genes with higher volume may be more credible.

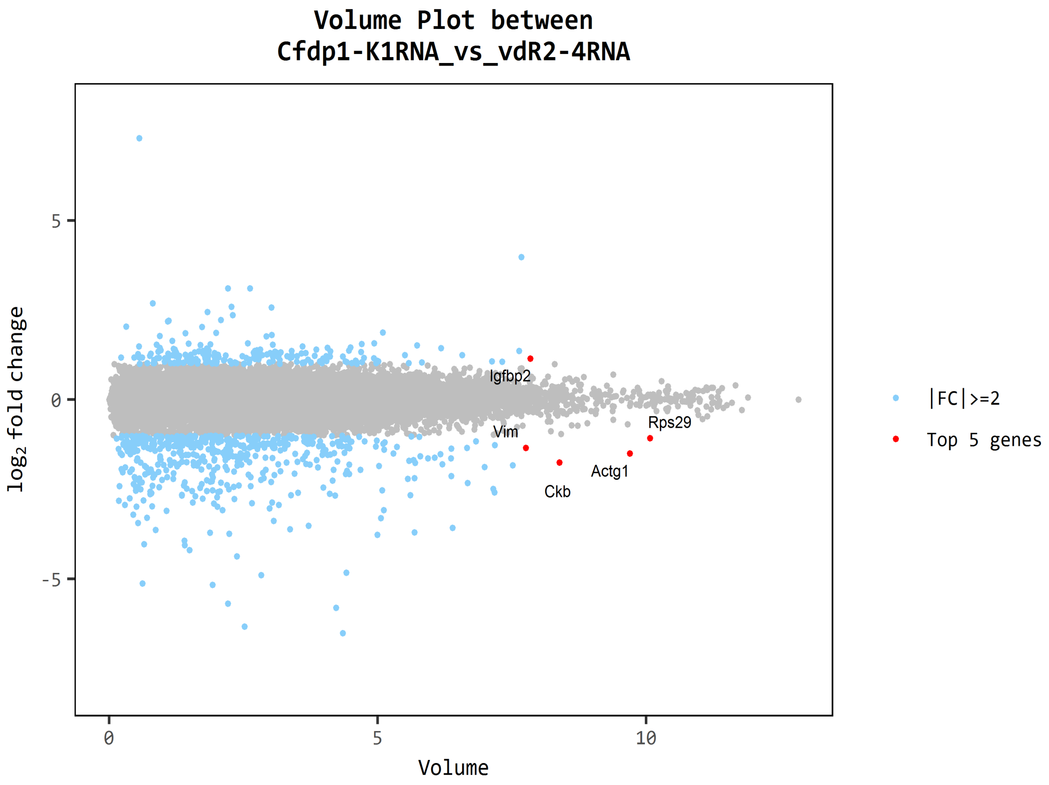


- Red dot: Top five genes by volume which satisfies, |fc|>=2

### 5. 3. 5. Hierarchical Clustering Analysis

(Refer to Path: result_RNAseq_excel/DEG_result/Cluster image/)

Heatmap shows result of hierarchical clustering analysis (Euclidean Method, Complete Linkage) which clusters the similarity of genes and samples by expression level (normalized value) from significant list.

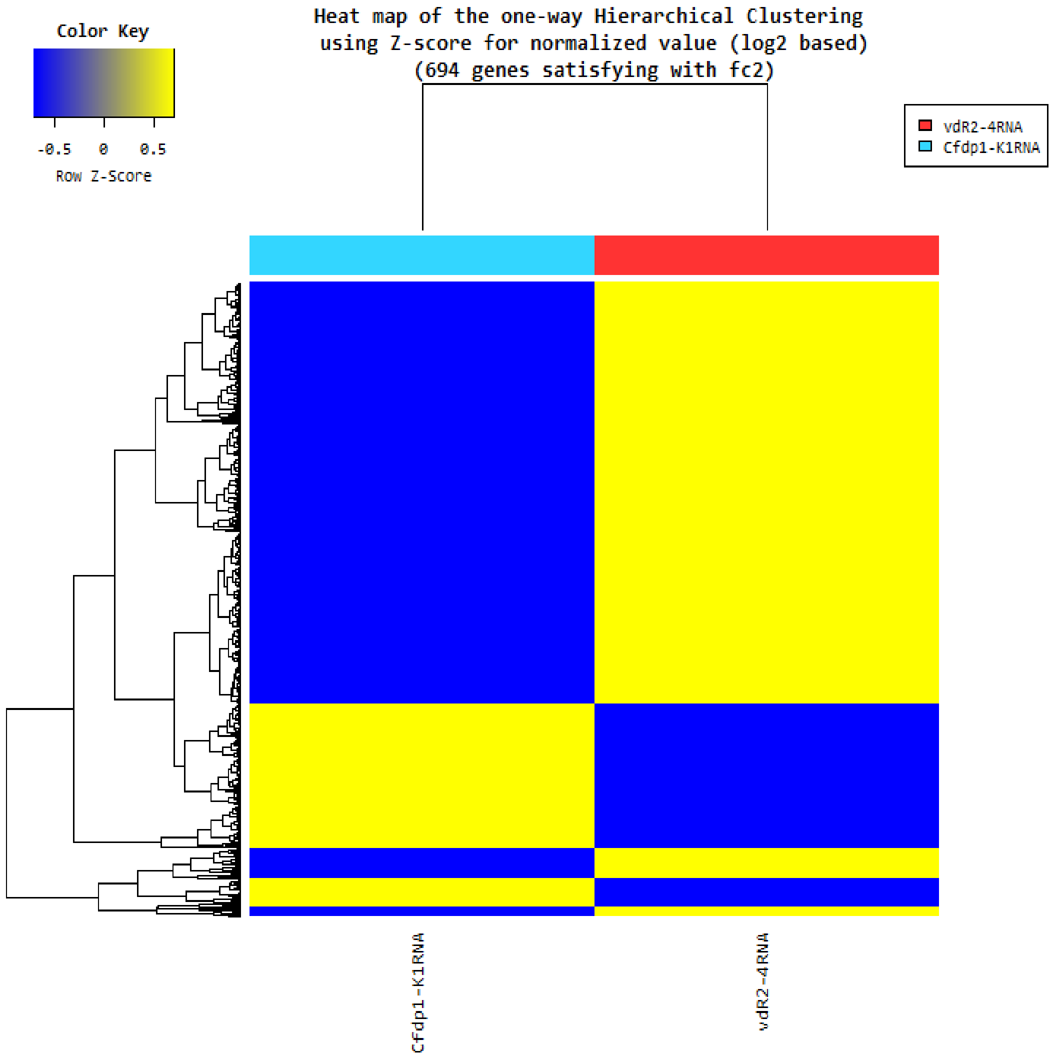

### 5. 4. GO Enrichment Analysis

(Refer to Path: result_RNAseq_excel/DEG_result/GO)

For Gene-Enrichment test which based on Gene Ontology (http://geneontology.org/) DB was conducted with significant gene list.

Progressing about 3 categories of GO. The gene or gene product, molecule associated with GO ID was summarized by parsing the ontology file and the annotation file (multispecies annotation provided by Uniprot, or the annotation provided by each type reference DB for the GO consortium) for the GO graph structure.

- Link for the ontology documentation: http://geneontology.org/page/ontology-documentation
- Link for the ontology files: http://geneontology.org/page/download-ontology
- Link for the annotation files: http://geneontology.org/page/download-annotations

The two results are provided for enrichment analysis.

- gomap_stat
- gomap_genes

### 5. 4. 1. gomap_stat Sheet

The result of associated gene and enrichment test was summarized by GO ID. The significance of specific GO ID in enrichment test with DEG set was calculated by modified fisher's exact test.

| Namespace | GOID | Term | count | Genes | Sig.NotIn.GO | Genome.In.GO | Genome.NotIn.GO | PValue | Bonferroni | FDR |
|---|---|---|---|---|---|---|---|---|---|---|
| cellular_component | GO:0005623 | cell | 736 | AAMDC, AARS, ABAT, ABCA3, ABCA7, AB | 205 | 115 | 59368 | 0 | 0 | 0 |
| cellular_component | GO:0044464 | cell part | 735 | AAMDC, AARS, ABAT, ABCA3, ABCA7, AB | 206 | 1286 | 58197 | 0 | 0 | 0 |
| biological_process | GO:0009987 | cellular process | 719 | AAMDC, AARS, ABAT, ABCA3, ABCA7, AB | 222 | 7204 | 52279 | 0 | 0 | 0 |
| biological_process | GO:0044699 | single-organism process | 684 | AAMDC, AARS, ABAT, ABCA3, ABCA7, AB | 257 | 5148 | 54335 | 0 | 0 | 0 |
| cellular_component | GO:0005622 | intracellular | 678 | AAMDC, AARS, ABAT, ABCA3, ABCA7, AB | 263 | 1147 | 58336 | 0 | 0 | 0 |
| cellular_component | GO:0043226 | organelle | 638 | AARS, ABAT, ABCA3, ABCA7, ABCB1, ABC | 303 | 1343 | 58140 | 0 | 0 | 0 |
| biological_process | GO:0044763 | single-organism cellular process | 631 | AAMDC, AARS, ABAT, ABCA3, ABCA7, AB | 310 | 1973 | 57510 | 0 | 0 | 0 |
| cellular_component | GO:0043227 | membrane-bounded organelle | 612 | AARS, ABAT, ABCA3, ABCA7, ABCB1, ABC | 329 | 4914 | 54569 | 0 | 0 | 0 |
| cellular_component | GO:0043229 | intracellular organelle | 576 | AARS, ABAT, ABCA3, ABCA7, ABCB9, ABC | 365 | 21 | 59462 | 0 | 0 | 0 |
| cellular_component | GO:0005737 | cytoplasm | 567 | AAMDC, AARS, ABAT, ABCA3, ABCA7, AB | 374 | 4904 | 54579 | 0 | 0 | 0 |
| biological_process | GO:0065007 | biological regulation | 555 | AAMDC, AARS, ABAT, ABCA7, ABCG1, AB | 386 | 94 | 59389 | 0 | 0 | 0 |
| cellular_component | GO:0043231 | intracellular membrane-bounded organelle | 541 | AARS, ABAT, ABCA3, ABCA7, ABCB9, ABC | 400 | 1895 | 57588 | 0 | 0 | 0 |
| biological_process | GO:0071704 | organic substance metabolic process | 524 | AAMDC, AARS, ABAT, ABCA7, ABCG1, AB | 417 | 2 | 59481 | 0 | 0 | 0 |
| biological_process | GO:0044237 | cellular metabolic process | 510 | AAMDC, AARS, ABAT, ABCA7, ABCG1, AB | 431 | 1195 | 58288 | 0 | 0 | 0 |
| biological_process | GO:0044238 | primary metabolic process | 507 | AAMDC, AARS, ABAT, ABCA7, ABCG1, AB | 434 | 18 | 59465 | 0 | 0 | 0 |
| biological_process | GO:0050794 | regulation of cellular process | 498 | AAMDC, AARS, ABAT, ABCA7, ABCG1, AB | 443 | 2 | 59481 | 0 | 0 | 0 |
| biological_process | GO:0050896 | response to stimulus | 447 | AARS, ABAT, ABCA3, ABCA7, ABCB1, ABC | 494 | 64 | 59419 | 0 | 0 | 0 |
| cellular_component | GO:0044444 | cytoplasmic part | 443 | AARS, ABAT, ABCA3, ABCA7, ABCB9, ABC | 498 | 7 | 59476 | 0 | 0 | 0 |
| cellular_component | GO:0016020 | membrane | 415 | AARS, ABCA3, ABCA7, ABCB1, ABCB9, AB | 526 | 601 | 58882 | 0 | 0 | 0 |
| cellular_component | GO:0044422 | organelle part | 413 | ABAT, ABCA3, ABCA7, ABCB9, ABCG1, AB | 528 | 87 | 59396 | 0 | 0 | 0 |
| cellular_component | GO:0044446 | intracellular organelle part | 407 | ABAT, ABCA3, ABCA7, ABCB9, ABCG1, AB | 534 | 16 | 59467 | 0 | 0 | 0 |
| biological_process | GO:0044260 | cellular macromolecule metabolic process | 379 | AAMDC, AARS, ABCA7, ABCG1, ABTB1, A | 562 | 14 | 59469 | 0 | 0 | 0 |
| biological_process | GO:0032501 | multicellular organismal process | 369 | AARS, ABAT, ABCA7, ABCG1, ACE, ACTA2, | 572 | 338 | 59145 | 0 | 0 | 0 |
| biological_process | GO:0044707 | single-multicellular organism process | 367 | AARS, ABAT, ABCA7, ABCG1, ACE, ACTA2, | 574 | 8 | 59475 | 0 | 0 | 0 |

- Namespace: 3 categories of Gene ontology (Cellular Component, Molecular Function, Biological Process)
- GOID: Gene ontology ID
- Term: Gene ontology term
- count: The number of unique genes associated with GO ID
- Genes: Associated genes with GO ID (connect by comma)
- Sig.NotIn.GO: The number of genes which are not associated GO ID
- Genome.In.GO: The number of genes associated with GO ID of total sample of gene species
- Genome.NotIn.GO: The number of genes which are not associated with GO ID of total sample of gene species
- PValue: Raw p-value was calculated by modified fisher's exact test
- Bonferroni: Adjusted p-value by Bonferroni
- FDR: Adjusted p-value by FDR

### 5. 4. 2. gomap_genes Sheet

The result of associated GO ID and DEG analysis was summarized based on gene. The GO ID which associated with specific gene was summarized with statistic such as fold change, p-value, volume, and normalized value.

| InID | OutID | GOID | Term | Namespace | PValue | Bonferroni | FDR | Gene_ID | Gene_Symbol | test/control.fc | test/control.volume | N_test | N_control |
|---|---|---|---|---|---|---|---|---|---|---|---|---|---|
| AAMDC | AAMDC | GO:0005488 | binding | molecular_function | 4.4834E-208 | 3.9544E-204 | 2.8864E-206 | 28971 | AAMDC | 1.899756 | 4.918387 | 5.885185 | 5.970135 |
| AAMDC | AAMDC | GO:0005515 | protein binding | molecular_function | 3.8301E-167 | 3.3782E-163 | 2.0108E-165 | 28971 | AAMDC | 1.899756 | 4.918387 | 5.885185 | 5.970135 |
| AARS | AARS | GO:0000049 | tRNA binding | molecular_function | 1 | 1 | 1 | 16 | AARS | 1.080088 | 6.531080 | 7.518749 | 7.629339 |
| AARS | AARS | GO:0000166 | nucleotide binding | molecular_function | 0.006121778 | 1 | 0.017222993 | 16 | AARS | 1.080088 | 6.531080 | 7.518749 | 7.629339 |
| AARS | AARS | GO:0001101 | response to acid chemical | biological_process | 1.61547E-30 | 1.42485E-26 | 1.98447E-29 | 16 | AARS | 1.080088 | 6.531080 | 7.518749 | 7.629339 |
| AARS | AARS | GO:0001882 | nucleoside binding | molecular_function | 4.5382E-146 | 4.0026E-142 | 2.0632E-144 | 16 | AARS | 1.080088 | 6.531080 | 7.518749 | 7.629339 |
| HNF1A | HNF1A | GO:0007267 | cell-cell signaling | biological_process | 1.20893E-98 | 1.06627E-94 | 3.63916E-97 | 6927 | HNF1A | -1.423774 | 3.386822 | 2.322587 | 2.153258 |
| HNF1A | HNF1A | GO:0007275 | multicellular organismal development | biological_process | 3.9889E-257 | 3.5182E-253 | 3.3829E-255 | 6927 | HNF1A | -1.423774 | 3.386822 | 2.322587 | 2.153258 |
| RARRES2 | RARRES2 | GO:0036211 | protein modification process | biological_process | 6.1358E-279 | 5.4118E-275 | 6.0131E-277 | 5919 | RARRES2 | -1.070216 | 6.260522 | 5.196256 | 4.595104 |
| VDR | VDR | GO:0003707 | steroid hormone receptor activity | molecular_function | 1.96245E-06 | 0.01730879 | 8.39418E-06 | 7421 | VDR | 1.664240 | 2.647718 | 3.457308 | 3.607864 |
| VDR | VDR | GO:0004871 | signal transducer activity | molecular_function | 9.73774E-71 | 8.58869E-67 | 2.27214E-69 | 7421 | VDR | 1.664240 | 2.647718 | 3.457308 | 3.607864 |
| VDR | VDR | GO:0004872 | receptor activity | molecular_function | 2.4367E-51 | 2.14917E-47 | 4.47744E-50 | 7421 | VDR | 1.664240 | 2.647718 | 3.457308 | 3.607864 |
| VDR | VDR | GO:0004879 | RNA polymerase II transcription factor ac | molecular_function | 2.87425E-07 | 0.002535084 | 1.31352E-06 | 7421 | VDR | 1.664240 | 2.647718 | 3.457308 | 3.607864 |
| VDR | VDR | GO:0005102 | receptor binding | molecular_function | 2.1529E-175 | 1.8988E-171 | 1.1868E-173 | 7421 | VDR | 1.664240 | 2.647718 | 3.457308 | 3.607864 |
| VDR | VDR | GO:0005488 | binding | molecular_function | 4.4834E-208 | 3.9544E-204 | 2.8864E-206 | 7421 | VDR | 1.664240 | 2.647718 | 3.457308 | 3.607864 |
| VDR | VDR | GO:0005496 | steroid binding | molecular_function | 1.3282E-09 | 1.17147E-05 | 7.29435E-09 | 7421 | VDR | 1.664240 | 2.647718 | 3.457308 | 3.607864 |
| VDR | VDR | GO:0005499 | vitamin D binding | molecular_function | 0.045949424 | 1 | 0.106960653 | 7421 | VDR | 1.664240 | 2.647718 | 3.457308 | 3.607864 |
| VDR | VDR | GO:0005515 | protein binding | molecular_function | 3.8301E-167 | 3.3782E-163 | 2.0108E-165 | 7421 | VDR | 1.664240 | 2.647718 | 3.457308 | 3.607864 |
| ZFP36 | ZFP36 | GO:0006952 | defense response | biological_process | 6.5961E-158 | 5.8178E-154 | 3.2321E-156 | 7538 | ZFP36 | 1.566431 | 2.191879 | 2.986505 | 3.233095 |
| ZFP36 | ZFP36 | GO:0006954 | inflammatory response | biological_process | 1.20554E-36 | 1.06328E-32 | 1.69313E-35 | 7538 | ZFP36 | 1.566431 | 2.191879 | 2.986505 | 3.233095 |
| ZFP36 | ZFP36 | GO:0007154 | cell communication | biological_process | 0 | 0 | 0 | 7538 | ZFP36 | 1.566431 | 2.191879 | 2.986505 | 3.233095 |
| ZFP36 | ZFP36 | GO:0007165 | signal transduction | biological_process | 0 | 0 | 0 | 7538 | ZFP36 | 1.566431 | 2.191879 | 2.986505 | 3.233095 |
| ZFP36 | ZFP36 | GO:0007275 | multicellular organismal development | biological_process | 3.9889E-257 | 3.5182E-253 | 3.3829E-255 | 7538 | ZFP36 | 1.566431 | 2.191879 | 2.986505 | 3.233095 |
| ZFP36 | ZFP36 | GO:0008152 | metabolic process | biological_process | 2.2907E-215 | 2.0204E-211 | 1.5423E-213 | 7538 | ZFP36 | 1.566431 | 2.191879 | 2.986505 | 3.233095 |

- InID: Iuput ID for GO enrichment analysis
- OutID: Mapping ID as gene symbol from input ID through GO enrichment analysis
- GOID: Gene ontology ID
- Term: Gene ontology term
- Namespace: 3 categories of gene ontology (Cellular Component, Molecular Function, Biological Process)
- PValue: Raw p-value was calculated by modified fisher's exact test
- Bonferroni: Adjusted p-value by Bonferroni
- FDR: Adjusted p-value by FDR

The bar plot below shows the results of the enrichment analysis based on Gene Ontology DB for significant genes.
(These plots were made based on gomap_stat result.)

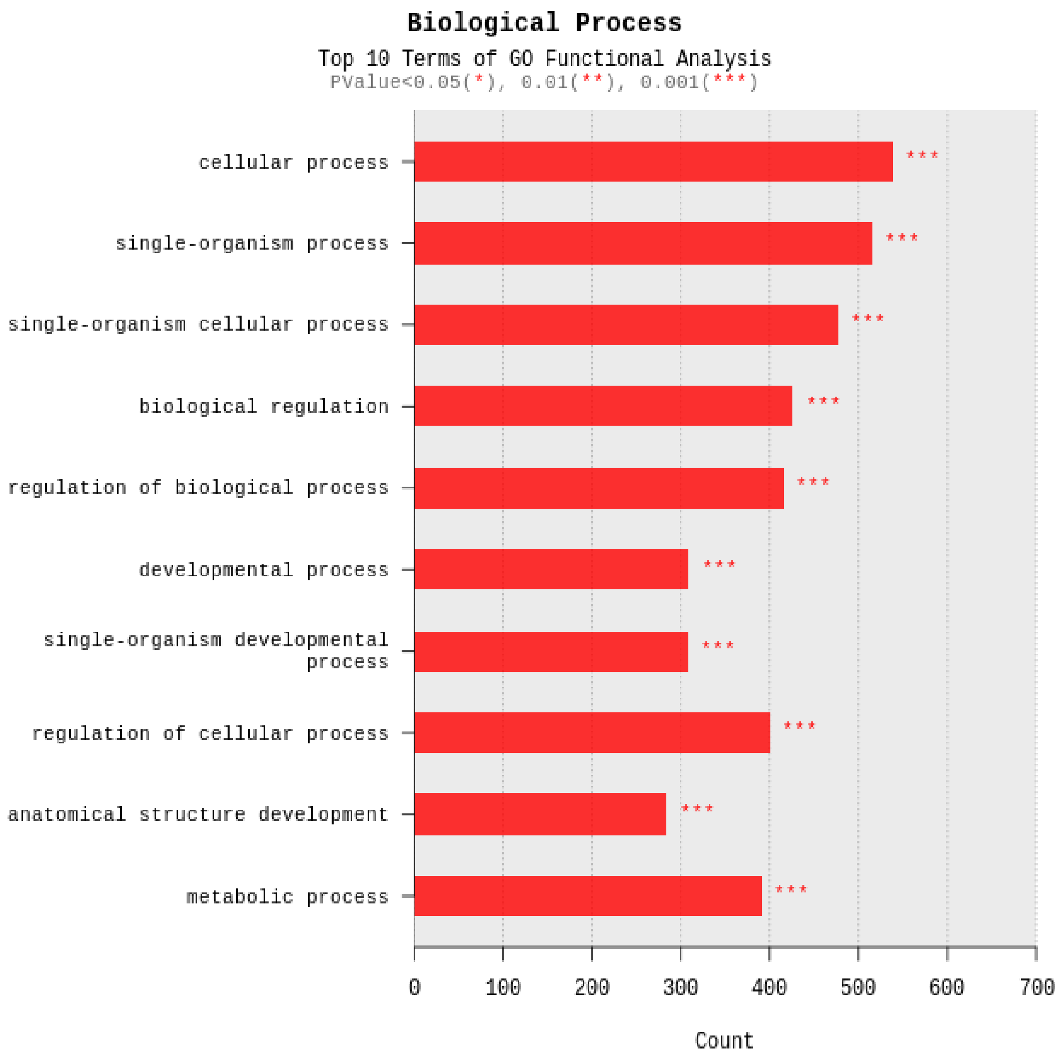

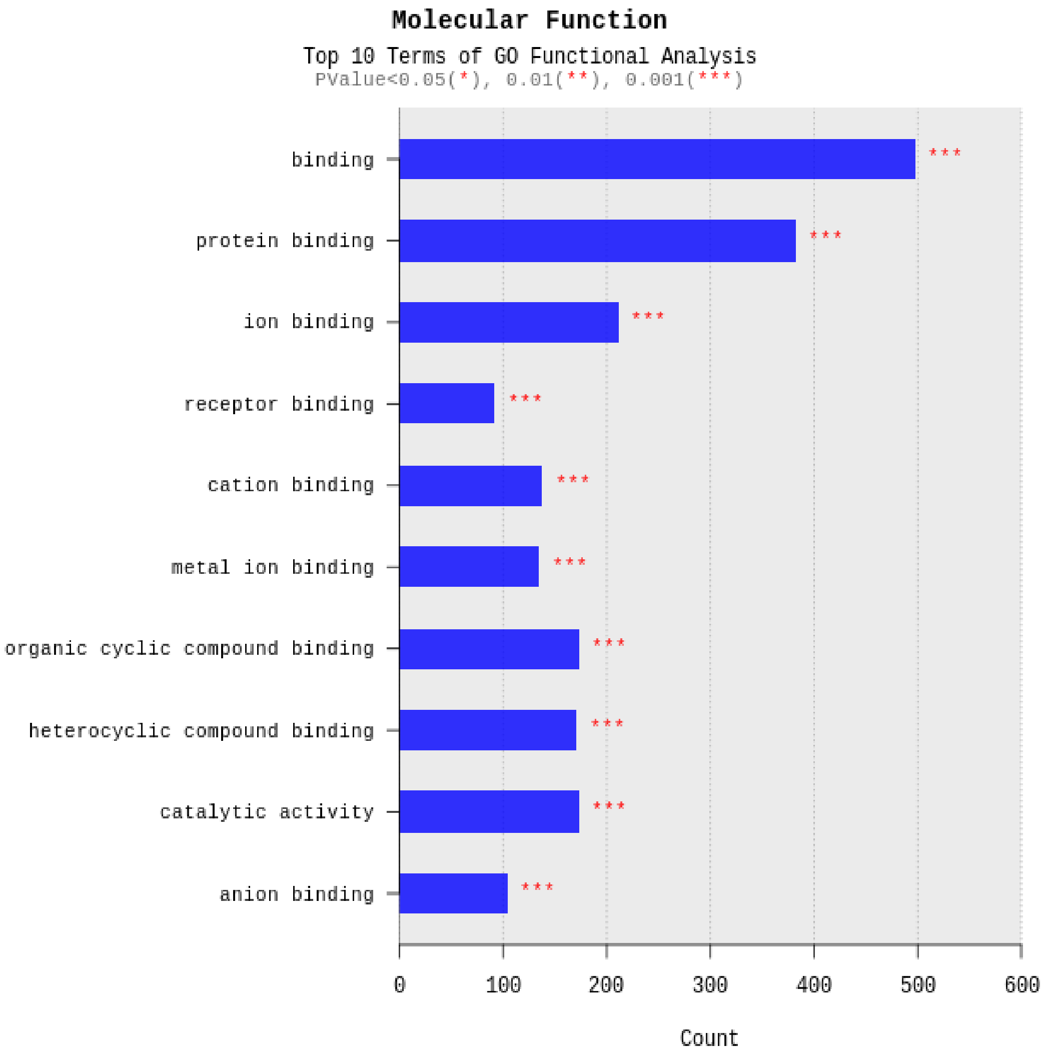
Molecular Function
Top 10 Terms of GO Functional Analysis
PValue<0.05(*), 0.01(**), 0.001(***)
binding
protein binding
ion binding
receptor binding
cation binding
metal ion binding
organic cyclic compound binding
heterocyclic compound binding
catalytic activity
anion binding
***
0
100
200
300
400
500
600
Count

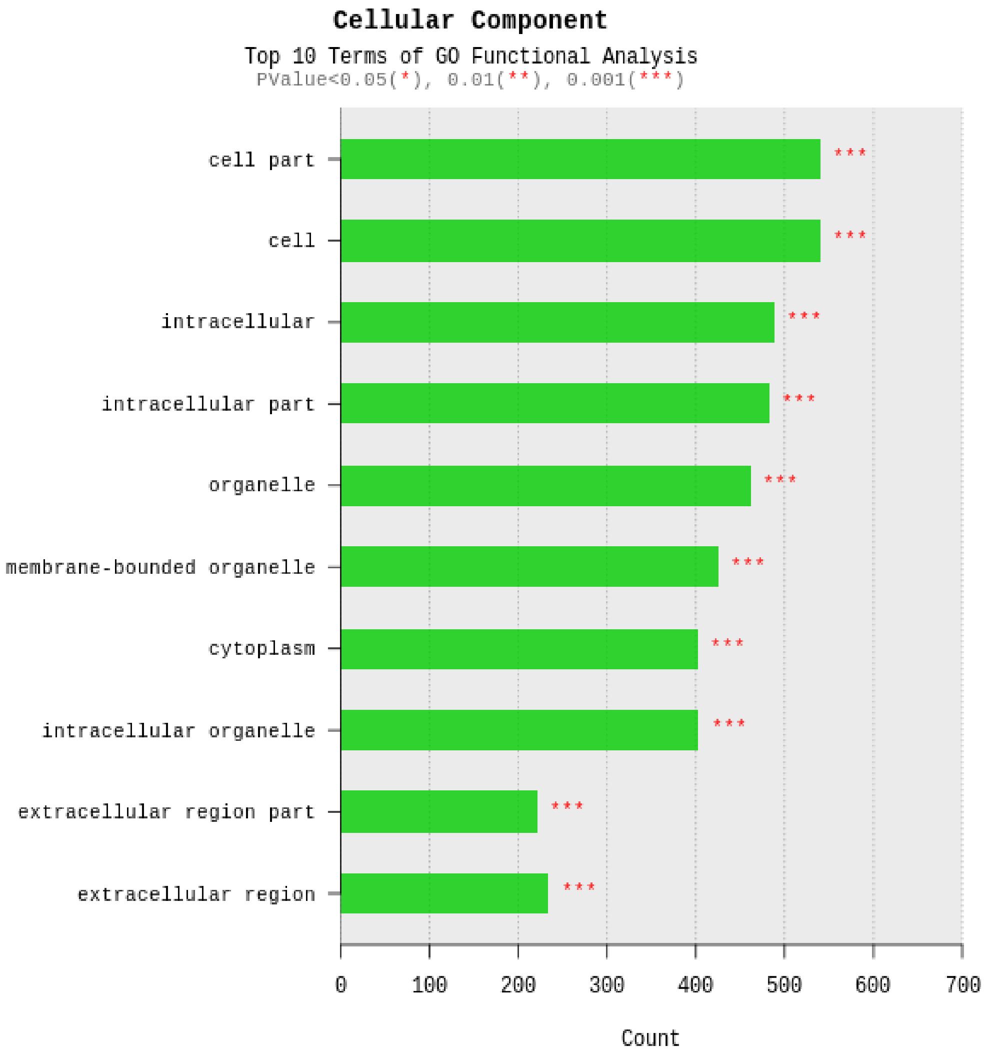

Cellular Component
Top 10 Terms of GO Functional Analysis
PValue<0.05(*), 0.01(**), 0.001(***)
cell part
cell
intracellular
intracellular part
organelle
membrane-bounded organelle
cytoplasm
intracellular organelle
extracellular region part
extracellular region
***
0
100
200
300
400
500
600
700
Count

### 6. Data Download Information

#### 6. 1. Raw Data

Raw data is the FASTQ file that isn't trimmed adapter sequence.

| Download link | File size | md5sum |
|---|---|---|
| Cfdp1-K1RNA_1.fastq.gz | 1.2G | 34d313aab0592c10d9a8c1181e14dfb5 |
| Cfdp1-K1RNA_2.fastq.gz | 1.23G | 68288067dba6036df06abc1a1fef372e |
| vdR2-4RNA_1.fastq.gz | 1.31G | a793383ab0387867c2d5edaae7627c38 |
| vdR2-4RNA_2.fastq.gz | 1.35G | ed193bf6585ada34d4a1d450d52971f1 |

- fastq.gz : This is a zip file of raw data used in analysis.
- md5sum : In order to verify the integrity of files, md5sum is used. If the values of md5sum are the same, there is no forgery, modification or omission.

#### 6. 2. Analysis Results

| Download link | File size |
|---|---|
| HN00101712_result_RNAseq.zip<br>(md5sum: 7f33ecd78a315ed837bf635cddbec896) | 30.45M |
| HN00101712_result_RNAseq_excel.zip<br>(md5sum: 621bddc7040d0df14b635ddd4db9ce28) | 14.35M |

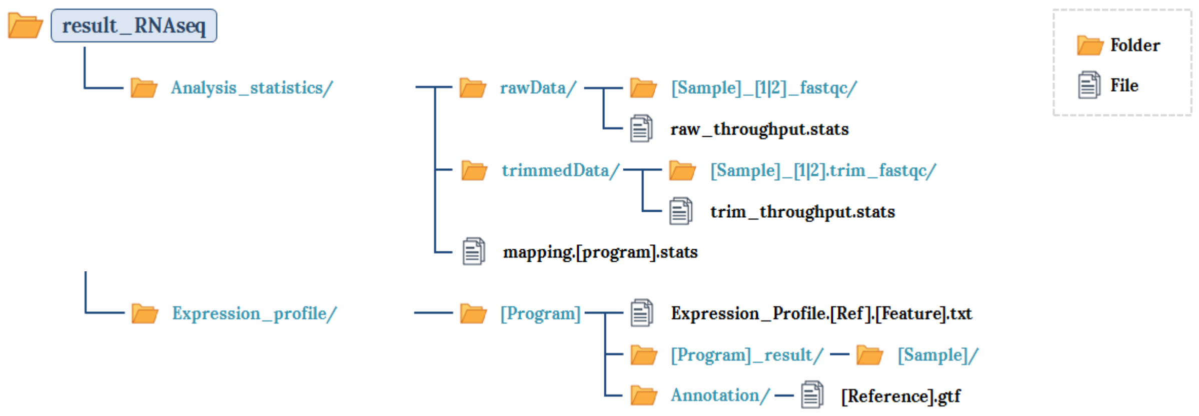
result_RNAseq
Analysis_statistics/
rawData/
[Sample]_[1|2]_fastqc/
raw_throughput.stats
trimmedData/
[Sample]_[1|2].trim_fastqc/
trim_throughput.stats
mapping.[program].stats
Expression_profile/
[Program]
Expression_Profile.[Ref].[Feature].txt
[Program]_result/
[Sample]/
Annotation/
[Reference].gtf
Folder
File

- Folder
- File

- result_RNAseq_excel
  - Expression_profile/
    - [Program]
      - Expression_Profile.[Ref].[Feature].xlsx
  - DEG_result/
    - Cluster image
    - Data Quality Check
    - GO
      - data3
      - data3_[comparison pair]
    - Plot
    - ...
    - data2.xlsx
    - data3_[filtering criteria].xlsx
    - data3_[comparison pair]_[filtering criteria].xlsx
    - Analysis_Result.html

⚠ The data retention period is three months,
or contact representative if you want longer retention period.

### 7. Appendix

#### 7. 1. Phred Quality Score Chart

Phred quality score numerically express the accuracy of each nucleotide. Higher Q number signifies higher accuracy. For example, if Phred assigns a quality score of 30 to a base, the chances of having base call error are 1 in 1000.

| Quality of phred score | Probability of incorrect base call | Base call accuracy |
|---|---|---|
| 10 | 1 in 10 | 90% |
| 20 | 1 in 100 | 99% |
| 30 | 1 in 1000 | 99.9% |
| 40 | 1 in 10000 | 99.99% |

Phred Quality Score Q is calculated with $-10\log_{10}P$, where P is probability of erroneous base call.

**Q-Score Binning**

Illumina NovaSeq sequencer groups quality scores into specific ranges, or bins, and assigns a value to each range. Q-scores is typically updated when significant characteristics of the sequencing platform changes, such as new hardware, software, or chemistry versions.

### 7. 2. Programs used in Analysis

#### 7. 2. 1. FastQC v0.11.7

LINK http://www.bioinformatics.babraham.ac.uk/projects/fastqc/

FastQC is a program that performs quality check on the raw sequences before analysis to make sure data integrity. The main function is importing BAM, SAM, FastQ files and providing quick overview on which section has problems. It provides such results as graphs and tables in html files.

#### 7. 2. 2. Trimmomatic 0.38

LINK http://www.usadellab.org/cms/?page=trimmomatic

Trimmomatic is a program that performs trimming depending on various parameters on illumina paired-end or single-end.

- ILLUMINACLIP: Cut adapter and other illumina-specific sequences from the read.
- SLIDINGWINDOW: Perform a sliding window trimming, cutting once the average quality within the window falls below a threshold.
- LEADING: Cut bases off the start of a read, if below a threshold quality.
- TRAILING: Cut bases off the end of a read, if below a threshold quality.
- CROP: Cut the read to a specified length.
- HEADCROP: Cut the specified number of bases from the start of the read.
- MINLEN: Drop the read if it is below a specified length.
- TOPHRED33: Change quality score to phred33.
- TOPHRED64: Change quality score to phred64.

#### 7. 2. 3. HISAT2 version 2.1.0, Bowtie2 2.3.4.1

LINK https://ccb.jhu.edu/software/hisat2/index.shtml

HISAT2 is a fast and sensitive alignment program for mapping next-generation sequencing reads to genomes. Its first implementation based on an extension of BWT for graphs, designed a graph FM index (GFM). In addition to using one global GFM index, HISAT2 uses a large set of small GFM indexes that collectively cover the whole genome (each index representing a genomic region of 56 Kbp, with 55,000 indexes needed to cover the human population). These small indexes (called local indexes), combined with several alignment strategies, enable rapid and accurate alignment of sequencing reads. This new indexing scheme is called a Hierarchical Graph FM index (HGFM).

#### 7. 2. 4. StringTie version 1.3.4d

LINK https://ccb.jhu.edu/software/stringtie/

StringTie is a fast and highly efficient assembler of RNA-Seq alignments into potential transcripts. It uses a novel network flow algorithm as well as an optional de novo assembly step to assemble and quantitate full-length transcripts representing multiple splice variants for each gene locus.

### 7. 3. References


1. BOLGER, Anthony M.; LOHSE, Marc; USADEL, Bjoern. Trimmomatic: a flexible trimmer for Illumina sequence data. Bioinformatics, 2014, btu170.
2. KIM, Daehwan; LANGMEAD, Ben; SALZBERG, Steven L. HISAT: a fast spliced aligner with low memory requirements. Nature methods, 2015, 12.4: 357-360.
3. LI, Heng, et al. The sequence alignment/map format and SAMtools. Bioinformatics, 2009, 25.16: 2078-2079.
4. PERTEA, Mihaela, et al. StringTie enables improved reconstruction of a transcriptome from RNA-seq reads. Nature biotechnology, 2015, 33.3: 290-295.
5. PERTEA, Mihaela, et al. Transcript-level expression analysis of RNA-seq experiments with HISAT, StringTie and Ballgown. Nature Protocols, 2016, 11.9: 1650-1667.

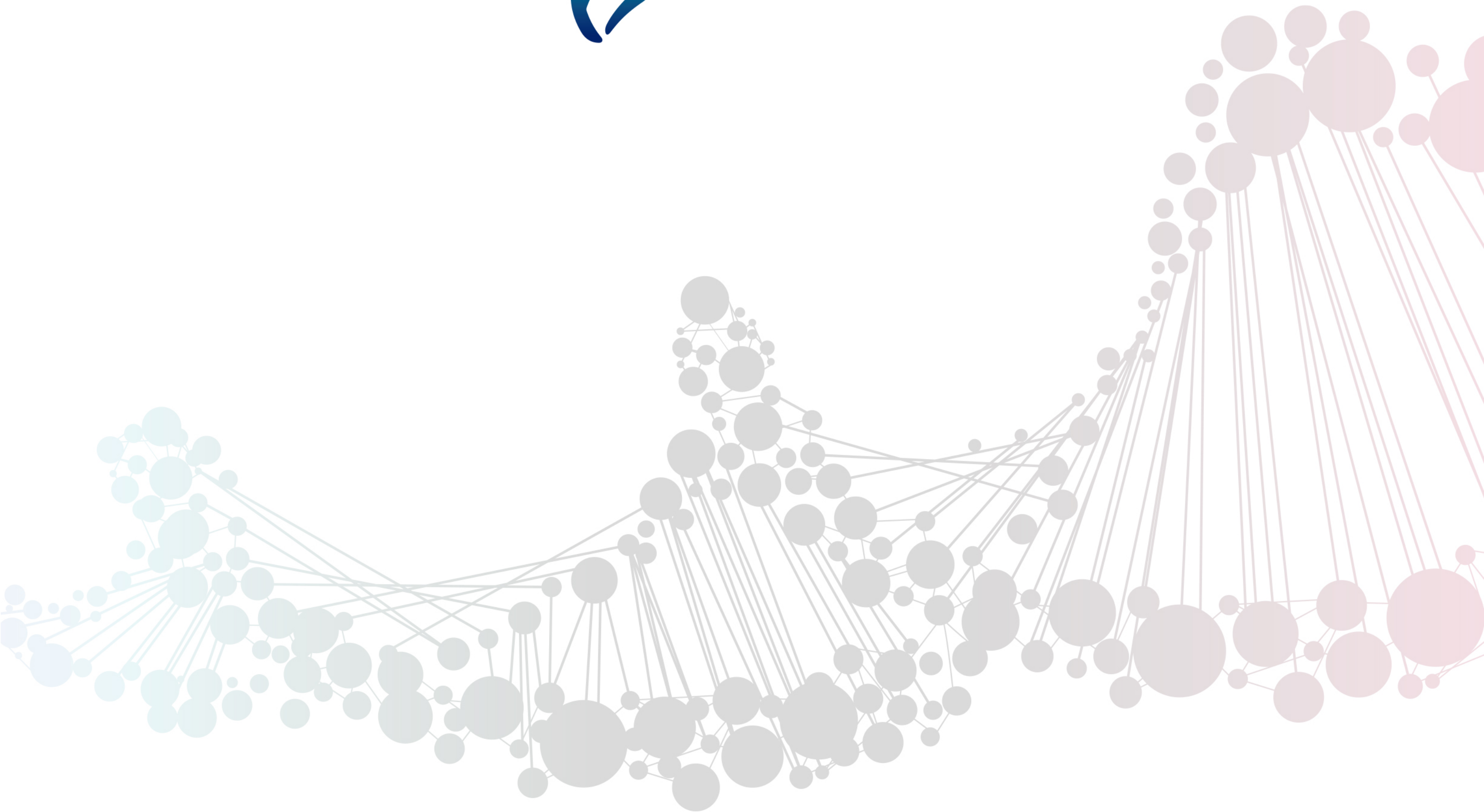


**Macrogen Inc.** | 08511 10F, 254 Beotkkot-ro, Geumcheon-gu, Seoul, Rep of Korea

**Contact us**
Site: www.macrogen.com | http://dna.macrogen.com
